# MuSpAn: A Toolbox for Multiscale Spatial Analysis

**DOI:** 10.1101/2024.12.06.627195

**Authors:** Joshua A. Bull, Joshua W. Moore, Shania M. Corry, Muyang Lin, Hayley L. Belnoue-Davis, Eoghan J. Mulholland-Illingworth, Simon J. Leedham, Helen M. Byrne

## Abstract

Advances in multiplex imaging and spatial omics have revolutionised spatial data generation in biology, revealing complex tissue organisation across multiple scales. However, methods for analysing these data have lagged behind, with fragmented, study-specific pipelines and limited guidance for tool selection. To address this, we introduce MuSpAn, a Multiscale Spatial Analysis package offering intuitive, flexible access to a wide range of mathematical tools - including spatial statistics, topological data analysis, geometry, and networks - within a unified framework. MuSpAn supports efficient data querying, is agnostic to imaging modality, and provides extensive documentation and community support. It enables users to create custom pipelines or conduct unbiased exploratory analyses. We demonstrate MuSpAn’s capacity to interrogate cross-compartmental cell interactions at multiple length scales in both normal and neoplastic tissue using mouse intestinal spatial transcriptomic datasets. Applied to a CMS4-like murine intestinal cancer model, MuSpAn identifies a continuum of fibroblastic functional phenotypes associated with discrete and coordinated fibroblast–immune interactions, highlighting its utility as a discovery tool across diverse biological contexts.

## Introduction

Coordinated interactions between diverse cell types define tissue architecture and aberrant disruption of these interactions is recognisably pathognomonic in disease. Advances in imaging technologies now enable these interactions to be mapped at unprecedented resolution, producing spatial datasets that span biological scales from subcellular features to entire organs [1–6]. Methods such as fluorescence in situ hybridization (FISH)-based imaging [7, 8], spatial transcriptomics [9], and spatial proteomics [10–13] generate rich, multiplexed views of tissue composition, yet this richness creates analytical bottlenecks that limit biological discovery.

Considerable progress has been made in addressing technical challenges such as storage, visualization, segmentation, and phenotyping, often with the help of Artificial Intelligence-based (AI) tools [14–21]. However, the central challenge remains: extracting meaningful biological insight from spatially complex data. Averaged metrics such as cell counts or densities obscure patterns of cellular positioning that often hold the key to biological function. For example, the prognostic significance of immune cells in cancer differs dramatically depending on whether they cluster at the tumour margin or infiltrate the lesion [22].

Crucially, cell signalling influences both cellular phenotypes and tissue organisation, and this ranges from local autocrine and paracrine pathways through to long range communication (e.g the endocrine system and extracellular vesicles) [23]. Resultant cell interactions occur across multiple spatial scales: from local, proximity-based interactions (e.g., antigen presentation) to formation of higher-order multicellular structures (e.g., tertiary lymphoid aggregates, intestinal crypts, hepatic lobules). Analyses focussed on a single scale risk overlooking essential biology, underscoring the need for multiscale methods that relate spatial patterns to underlying processes [24].

Most existing software tools focus on specific imaging modalities or biological questions, typically implemented as fixed, sequential pipelines [25–31]. For instance, Schürch et al. [25] used sliding windows to characterise neighbourhoods; SpottedPy applies Getis-Ord hotspot analysis to Visium data [26]; and SPIAT employs cross-K functions and clustering [27]. While powerful in their proposed contexts, it can be difficult to compare methods and select appropriate tools due to differences in programming language, imaging modality, and chosen analytical approaches. Few platforms support systematic comparison and integration of diverse methods; notably SquidPy provides a range of graph-based approaches with efficient data structures [32, 33]. Opportunities to combine complementary mathematical approaches therefore remain underexploited, despite improving feature detection [24].

To address these challenges, we introduce **MuSpAn**, a Python library for **Mu**ltiscale **Sp**atial **An**alysis. MuSpAn unifies more than 100 methods for quantifying spatial relationships across biological scales. Researchers can interactively compare analytical strategies tailored to their own questions, from cell–cell contacts to tissue geometry and topology. Critically, MuSpAn enables cross-scale integration, allowing insights at one level (e.g., subcellular expression) to inform analyses at another (e.g., tissue microenvironment). Its modular design supports incorporation of new methods, is underpinned by robust testing, and is accompanied by detailed tutorials, comprehensive documentation, and an active online user-developer community. Here, we demonstrate the capability of MuSpAn to interrogate cross-cell compartment interactions across length scales in normal and neoplastic mouse intestine. Together, these approaches illustrate how MuSpAn detects hierarchical organisation across scales - from single-cell positioning to tissue compartmentalisation - enabling applications such as immune niche mapping and structural feature discovery.

## Results

### MuSpAn provides a framework for quantitative spatial analysis at multiple scales

MuSpAn is a Python package for spatial analysis, applicable following segmentation and phenotype assignment (Fig. 1A). It accepts spatial coordinates (e.g., centroids, boundaries) and metadata (e.g., cell types, marker intensities, morphometrics), and is compatible with outputs from generic segmentation and phenotyping pipelines, or frameworks including SpatialData [19, 33], QuPath [14], and Xenium Explorer (10x). The package is supported by extensive documentation and tutorials (https://www.muspan.co.uk) and rigorous unit testing.

**Fig. 1:**
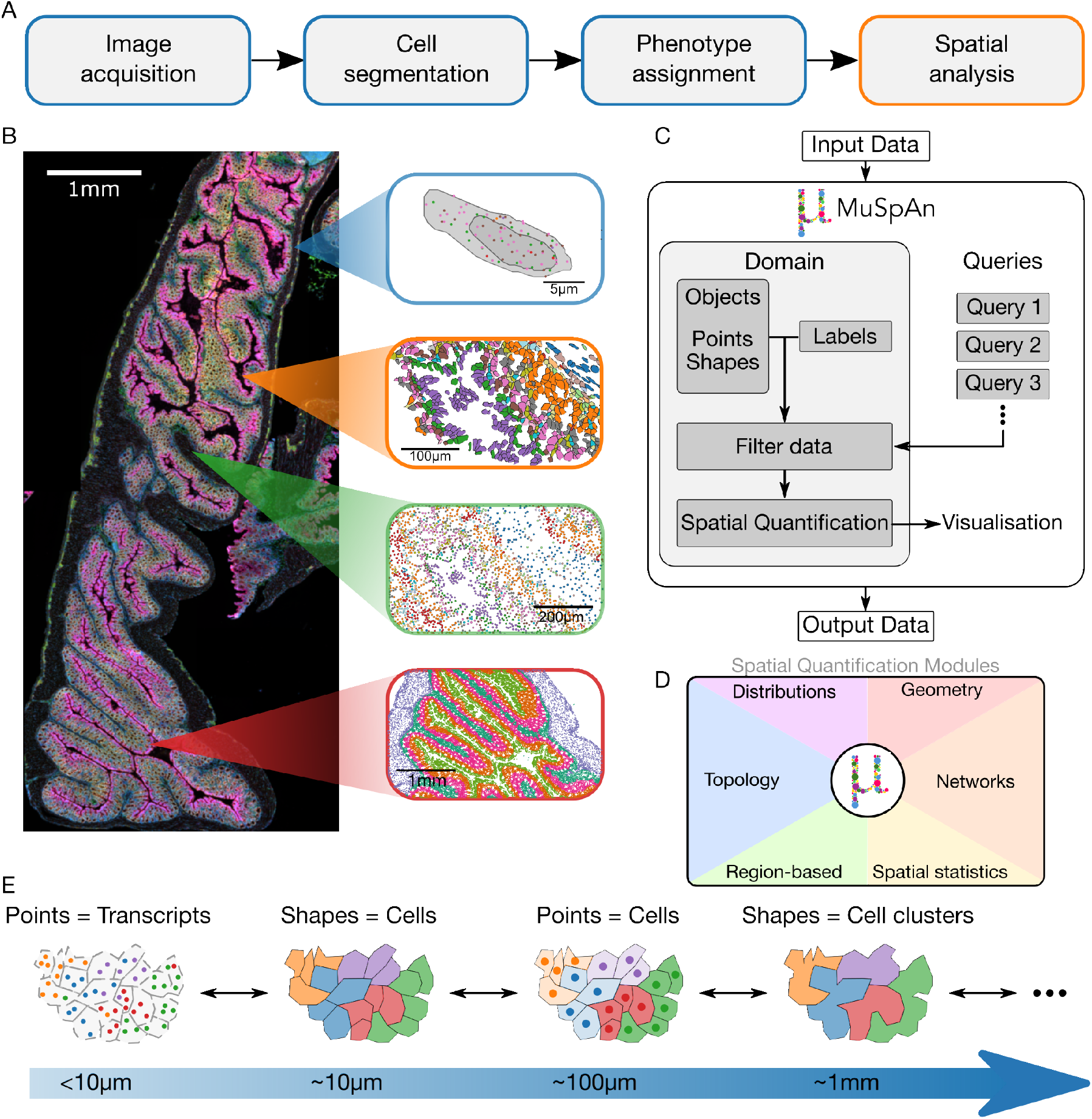
Overview of MuSpAn. A: Workflow for processing spatial biology. B: Healthy mouse colon from the Xenium platform, illustrating spatial structure across length scales. C: Schematic of the internal structure of MuSpAn. D: Spatial quantification methods are organised into six modules. E: MuSpAn represents data as geometric objects that can be defined at any length scale. Analytical methods can be applied at any scale, provided the objects are structurally compatible.

MuSpAn enables flexible and interactive analysis across length scales. For example, spatial transcriptomics data may describe subcellular transcript localisation (*<* 10µm), cell-cell contact (≈ 100µm), or tissue structures (≥ 1000µm) (Fig. 1B). Rather than focus on a fixed length scale, MuSpAn represents spatial data as geometric objects (points or shapes) which could reflect diverse spatial features (e.g., individual transcripts, cell centres, cell boundaries, user-defined annotations; Fig. 1C). Objects and associated metadata labels are stored within a single domain, which users may query to specify subsets of objects via spatial cues (e.g., ‘points inside a hypoxic region’, ‘cell boundaries within 50µm of the tumour invasive margin’), labels (e.g., ‘macrophages’, ‘cells highly expressing PD-L1’, ‘cellular neighbourhoods enriched with T cells’), or other object properties. Over 100 analysis methods are available to interrogate objects within a given region of interest (ROI, shape object), described in detail in the Supplementary Information. The methods are organised into six modules (Fig. 1D), and each can be applied to objects at any length scale. Complex objects can be simplified (e.g., shapes to centroids) to facilitate analyses at different scales, propagating embedded metadata and reducing computational cost (Fig. 1E).

Below, we apply MuSpAn to Xenium datasets of healthy mouse colon and a consensus molecular subtype 4 (CMS4)-like murine colorectal cancer model. The first example showcases MuSpAn’s ability to identify structural features across multiple scales in a publicly available tissue, while the second shows how it can be used to identify new hallmarks of disease using a bespoke Xenium panel and targeted analysis.

### Validation of MuSpAn in a healthy mouse colon exemplar

We demonstrate MuSpAn’s capacity for exploratory data analysis across length scales using an ‘off-the-shelf’ dataset of healthy mouse colon from the Xenium platform (10x). The gene panel and cell annotations represent a generic subset of cells whose behaviour is well characterised in the healthy colon. We show how some of the quantitative tools within MuSpAn can identify this expected cell behaviour, validating their use for spatial data analysis and showcasing some of the features outlined in Fig. 1. These include analysis of data across different length scales, calculating statistics within irregularly shaped regions, and the use of multiple complementary methods to support the same biological conclusions. First we consider cell-scale geometry and subcellular transcript localisation, then we progress up the spatial scales to characterise the whole tissue section.

### Cell geometry and subcellular transcript localisation

MuSpAn can analyse transcriptional polarisation and morphometrics at subcellular resolution, through characterisation of the spatial localisation of transcripts associated with individual Smooth

Muscle Cells (SMCs) and Epithelial Progenitor cells (PROG) in the normal mouse colon (Fig. 2). Within a small ROI (0.13mm^2^), we identified five transcripts preferentially expressed by PROG cells (*Ccl9, Sox9, Nupr1, Cldn2, Oit1*) and five preferentially expressed by SMCs (*Mylk, Myl9, Cnn1, Mgll, Mustn1*), using the segmentation mask to define the shapes of individual cell boundaries and nuclei for analysis. For representative cells, we quantify both cell geometry and transcript localisation within the nucleus and cytoplasm (Fig. 2B,C). For each metric, MuSpAn facilitates visualisation across these cell subtypes in the ROI (Fig. 2D-G) and quantification of differences in its distribution between SMC and PROG populations (Fig. 2H-K). The cell types have comparable areas, but we find significant differences in cell circularity and principal angle consistent with the tissue orientation of these different individual cell types. These results show that by interpreting individual transcripts as points MuSpAn can quantify subcellular transcript distributions and relate them to individual cell geometric features such as morphology, polarisation and tissue orientation.

**Fig. 2:**
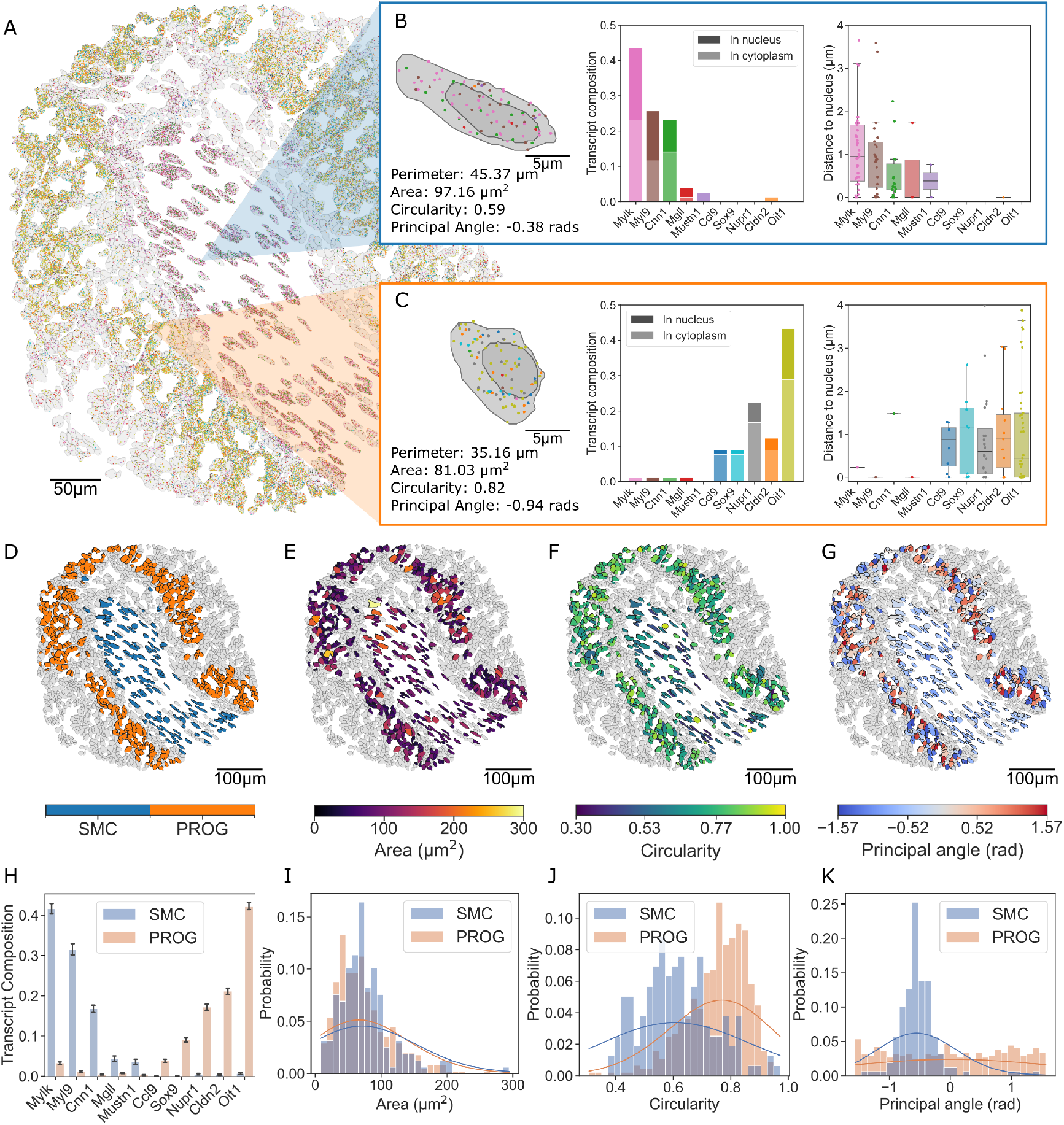
Quantification at the cell scale. A: ROI from healthy mouse colon. B, C: Morphological metrics and transcript summary statistics for a representative Smooth Muscle Cell (SMC; B) and Epithelial Progenitor cell (PROG; C). D-G: Cell summary labels visualised as heatmaps colouring the cell boundaries for SMCs and PROG cells (D: Celltype. E: Area. F: Circularity. G: Principal angle.) H: Proportion of selected transcripts within SMCs and PROG cells (mean, 95% CI). I-K: Distributions of morphological metrics from (F)-(G), by Celltype (metric, Wilcoxon rank-sum test p-value). I: area, *p* = 0.26. J: circularity, *p <* 10^−3^. K: principal angle, *p <* 10^−3^. Solid lines show Gaussian KDEs of continuous distributions.

### Quantification of cell proximity and interaction networks

Next, we used three complementary methods within MuSpAn to examine proximity based cell interaction networks within a larger ROI (≈ 0.38 mm^2^, Fig. 3A) in the normal mouse colonic mucosa. First, nearest-neighbour distances revealed strong (non random) proximity for SMCs and Type 1 Stromal cells (STR1) (10.9µm, Fig. 3B-C) within the intestinal lamina propria with contrasting exclusion for PROG cells to Colonocytes (COLs) (106.0µm, Fig. 3D-E) which were distributed differently along the epithelial vertical axis. *z*-scores from the Average Nearest Neighbour Index [27, 34] confirmed significance (*p <* 10^−3^). Secondly, contact networks (Fig. 3F) showed that SMCs were relatively isolated (low degree, Fig. 3G) but frequently connected to STR1s, consistent with the anticipated presence of these cell types in the lamina propria (Fig. 3H). Lastly, quadrat correlation analysis confirmed positive correlation for SMC-STR1 and exclusion for PROG-COL (Fig. 3I-K), and also identified other, non-random, positive and negative cell associations. Together, these data highlight MuSpAn’s ability to integrate multiple analytical methods to assess proximity based cell interactions. The convergence of multiple mathematical techniques in identifying the same cell relationships provides reassurance of biological relevance [24]. These tools can be used to rapidly identify significant cell interactions across heterogenous cell types within a tissue ecosystem. Applied in this way, MuSpAn acts as a hypothesis generating discovery tool to quantify local tissue cellular organisation and to begin to decipher the signalling networks that underpin key identified proximity-based cell interactions.

**Fig. 3:**
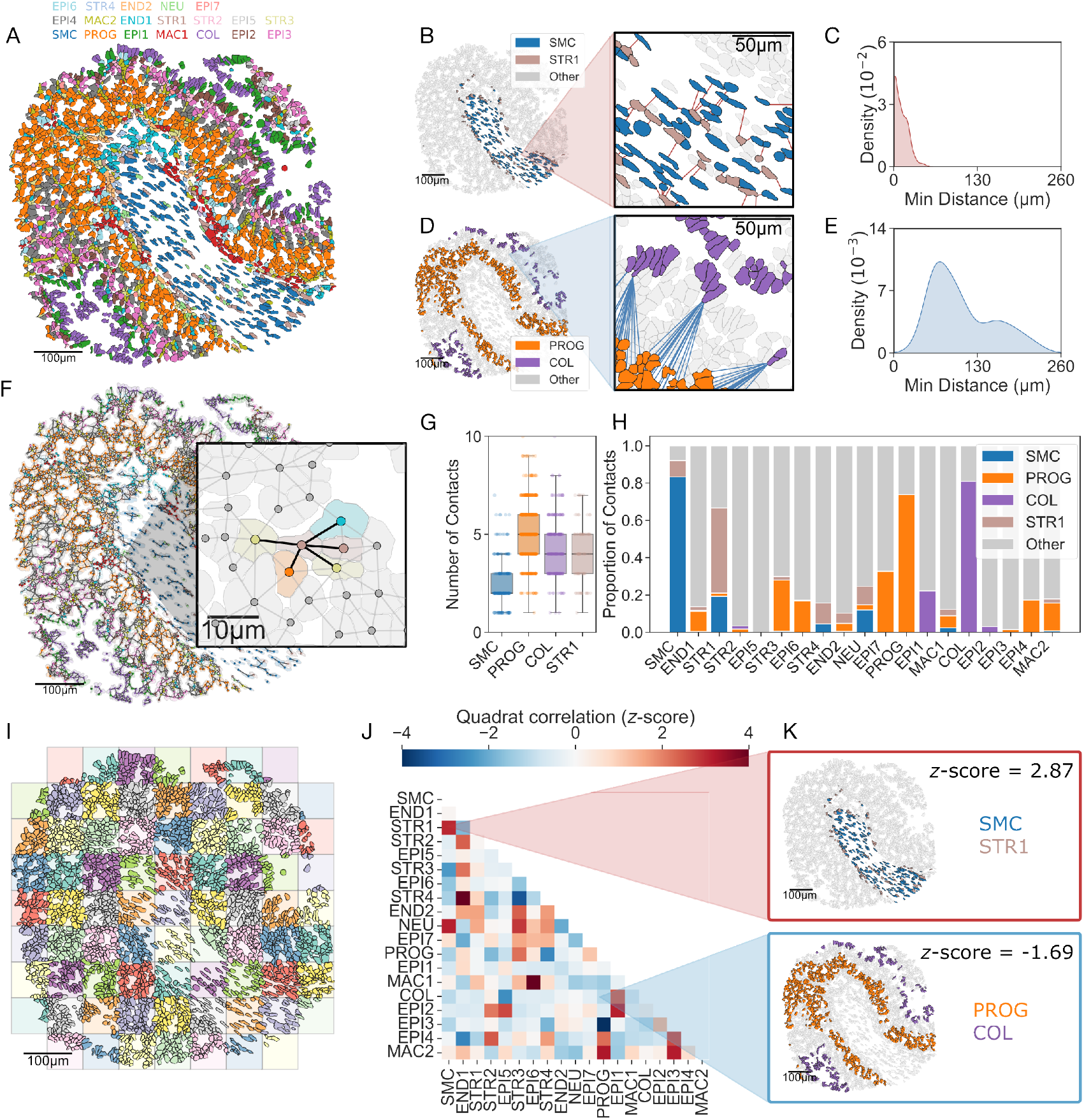
Quantification of cell-cell proximity. A: Cell boundaries coloured by cell type. B-E: Nearest neighbour distances for individual cells (B: SMC-STR1; D: PROG-COL) and their distributions (C: SMC-STR1, E: PROG-COL) F: Proximity network for the ROI in A, with nodes placed at cell centroids and edges present when cell boundaries are within 1µm. G: Degree distribution for selected cell types. H: Mean proportion of neighbours for selected cell types. I: Tiling of the ROI into 75µm *×* 75µm quadrats. J: Quadrat correlation matrix (QCM) for I. K: Locations of selected cell pairs, corresponding to colocalisation (SMC-STR1, *z* = 2.87) and exclusion (PROG-COL, *z* = −1.69)

### Assessment of multiscale spatial patterns

Having identified some proximity based cell interactions and exclusions, we next used two complementary techniques to assess the distribution and topology of epithelial (PROG) and smooth muscle (SMC) cell types. We consider a larger ROI (≈ 1.13 mm^2^, Fig. 4A) showing the spatial position of cell centroids within a section of normal colonic tissue architecture. The cross pair correlation function (cross-PCF) identified short to medium range clustering of PROG cells (Fig. 4B, *g >* 1 for *r <* 150µm) but exclusion of PROG-SMCs (Fig. 4B, *g <* 1 for *r <* 80µm), consistent with the discrete compartmentalisation of the epithelium and sub-crypt lamina propria. MuSpAn automatically corrected for boundary effects involving points located close to the boundary of the ROI (Fig. 4B insets). These results show how the cross-PCF can quantify both the strength and length scale of positive and negative cell associations. We also used persistent homology (PH), a form of topological data analysis (TDA), to identify loop-like (‘*H*_1_’) features in the distribution of PROG cells with an approximate radius of ≈ 25µm, corresponding to transected colonic crypts (Fig. 4C). PH captures birth and death radii of these features, consistent with intercellular spacing and crypt diameters, respectively. This analysis demonstrates the ability of PH to identify and quantify the organisation of multiple cell types into higher-order tissue structural features such as crypts, alveoli, or hepatic lobules which may be difficult to discern with conventional correlation analyses. The loop features may define normal tissue architecture which can be lost during disease progression, as seen in inflammation or neoplasia.

**Fig. 4:**
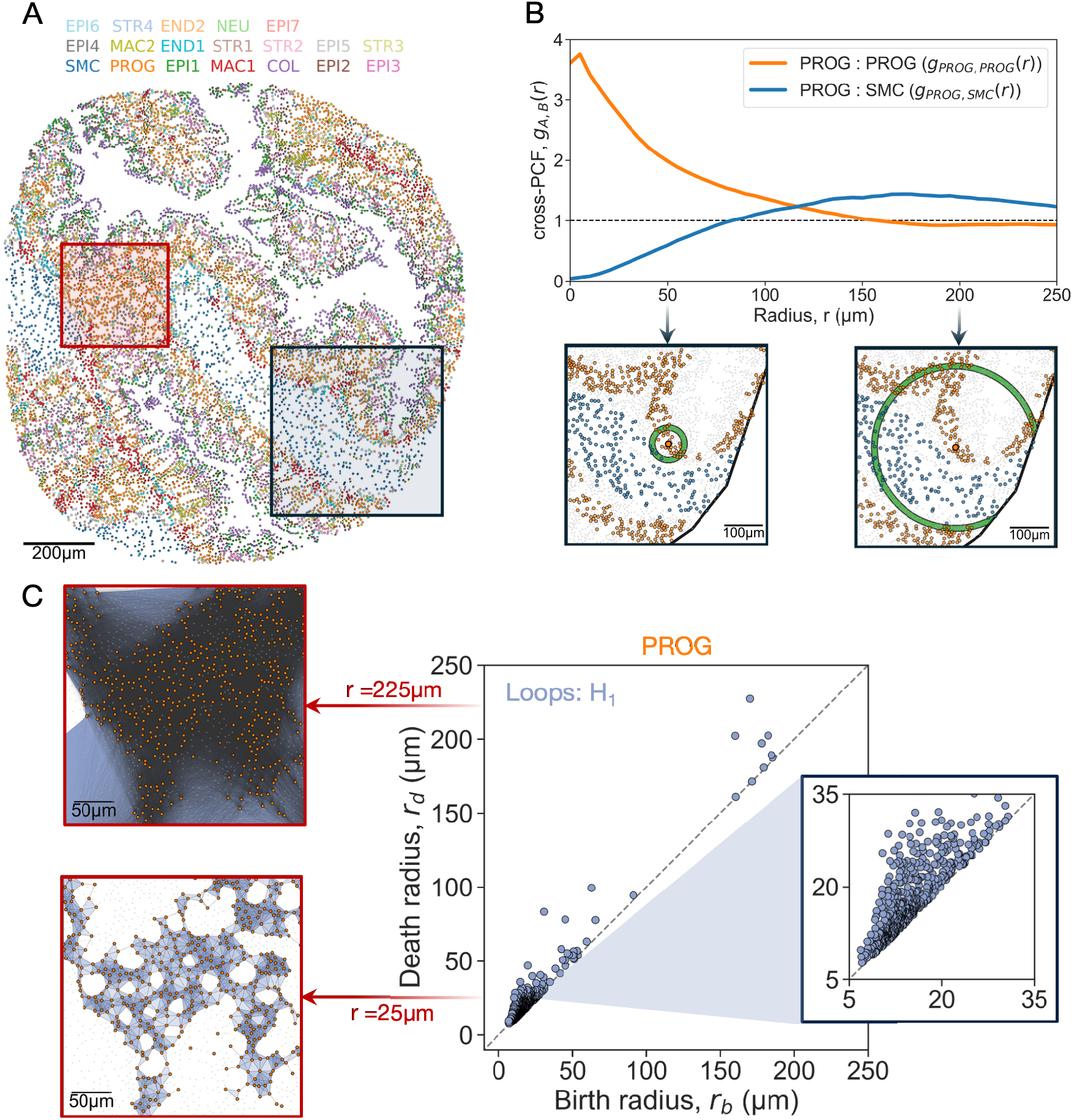
Quantification of cell-cell relationships across length scales. A: Cell centres coloured by cell type. B: Cross-PCFs for PROG-SMC (blue curve, exclusion) and PROG-PROG (orange curve, colocalisation). Annuli of radii 50µm and 200µm were used to calculate the cross-PCF, with automatic boundary correction (see insets, corresponding to black box in A). C: Persistence diagram of the VR filtration, showing loops (*H*_1_, purple). Stages involved in constructing the VR filtration on PROG cells (red box in A) with simplicial complexes at 25µm and 225µm are shown on left.

### Resolving tissue-scale organisation

We used three complementary methods to characterise the tissue organisational structures within the normal mouse colon at the level of the whole section (Fig. 5A). We used Delaunay graphs (≤ 30µm) to define cellular neighbourhoods; assigning each cell centre to one of five distinct microenvironments (MEs) according to the composition of its 3-hop neighbourhood (Fig. 5B-E) [25, 35, 36]. Contiguous regions of cells assigned to the same ME were converted into shapes to conduct an adjacency permutation test (APT) which revealed ordered transitions from the serosal wall through to the epithelial cell compartment (Fig. 5E–G).

**Fig. 5:**
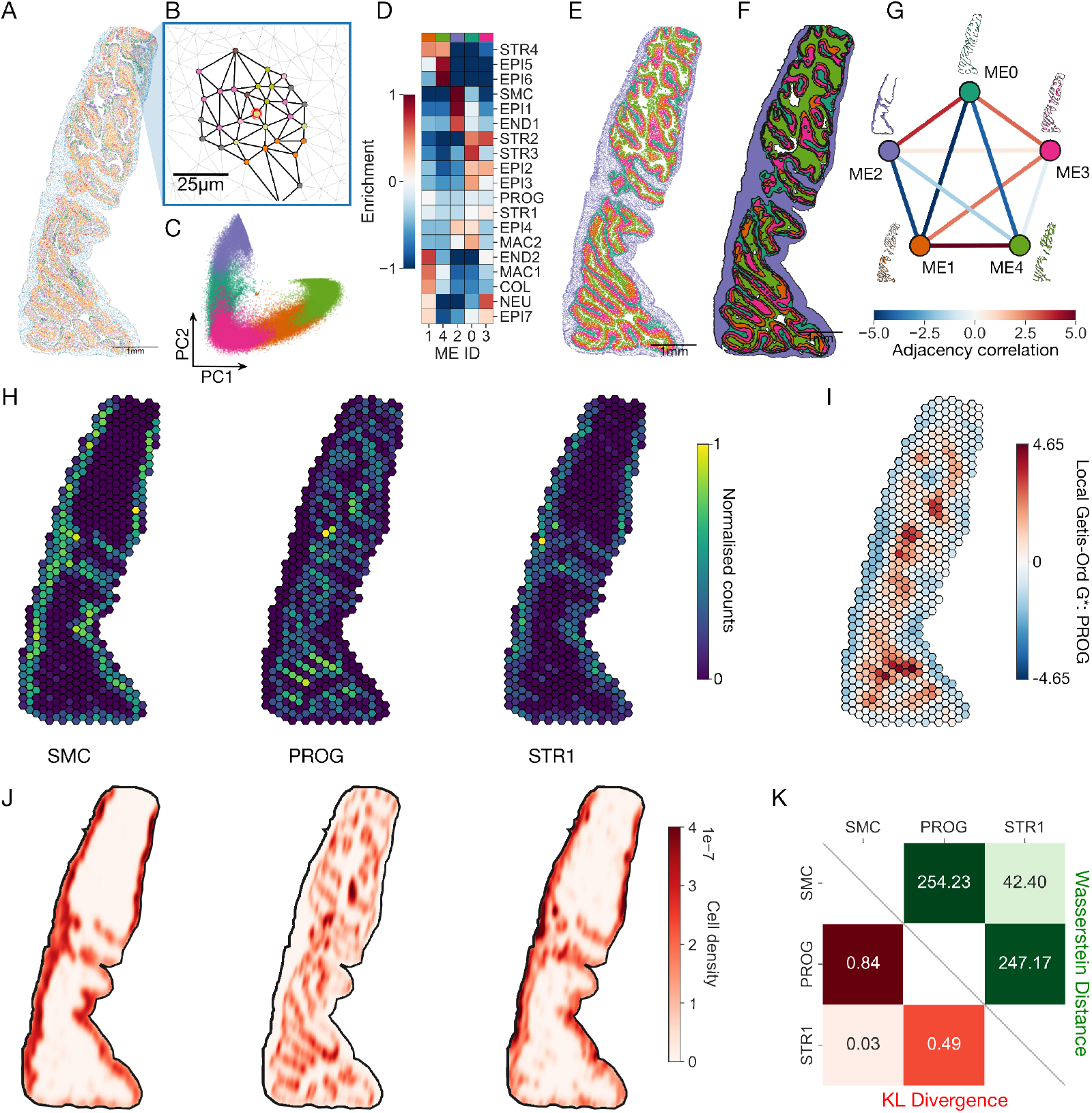
Quantification of tissue scale features. A-G: Microenvironment analysis. H-I: Region-based analysis. J-K: Distribution-based analysis. A: Large ROI from the healthy mouse dataset, cell centres coloured by cell type. B: 3-hop neighbourhood centred on the red-outlined cell. C: PCA of 3-hop neighbourhood compositions, with cells assigned microenvironment (ME) IDs via *k*-means clustering (*k* = 5). D: Cell type composition of the 5 microenvironments. E: Cell centres coloured by ME ID. F: Shapes formed by grouping spatially contiguous cell centres with the same ME ID. G: Adjacency correlation of shapes with different ME IDs. H: Cell counts of selected cell types within a hexagonal lattice (100µm sides). I: Getis-Ord^*^ for PROG cells. J: Kernel density estimates (KDEs) of selected cell types. K: KL-divergence (red, lower triangle) and Wasserstein distance (green, upper triangle) between the KDEs in J.

Secondly, we compartmentalised the ROI into a hexagonal lattice of side length 100µm. Cell counts within each hexagon (Fig. 5H) highlighted peripheral enrichment of muscularis cell types SMC and STR1 and central hotspots of PROG, detected by Local Getis-Ord^*^ (Fig. 5I).

Lastly, we use kernel density estimation (KDE) to generate smooth maps of distributions (Fig. 5J). KL divergence and Wasserstein distance quantified overlap and divergence of cell population distributions (Fig. 5K).

These methods each demonstrate how relationships between individual cells can define tissue-scale cell compartments, and show how MuSpAn can quantify the distribution and interactions between tissue structures across whole tissue sections.

The previous examples characterise healthy mouse intestinal tissue, within which the interactions we have described are well known. We now consider a bespoke Xenium panel for a murine colorectal cancer model, designed to characterise interactions between fibroblasts and immune cell subtypes. We show how MuSpAn can be leveraged to investigate specific cell relationships and uncover new biological insight.

### Multiscale spatial profiling of fibroblast–immune interactions in murine intestinal neoplasia

Having demonstrated the capacity of MuSpAn to conduct exploratory analyses of multiscale, cross compartment cell interactions in normal mouse colon tissue, we next used it for focussed analysis of pathological cancer-associated fibroblast-immune cell interactions in murine intestinal neoplasia. We undertook analysis of a bespoke 480-gene Xenium spatial transcriptomics panel on a dataset of intestinal tumours arising in *Villin-CreER*^*T2*^*;Apc*^*fl/fl*^*;Kras*^*G12D/+*^*;Trp53*^*fl/fl*^ mice (hereafter AKPT) (*n* = 4), as they generate stromal rich, metastatic, CMS4 cancers [37]. A multiscale workflow (Fig. 6A) was constructed, analysing immune context around cancer-associated fibroblast subtypes (Fig. 6B-E) at three scales: direct cell–cell contact (*<* 10µm, Fig. 6F-H), immune neighbourhood composition (*<* 60µm, Fig. 6I-M), and pairwise spatial correlations (*<* 500µm, Fig. 6N-P).

**Fig. 6:**
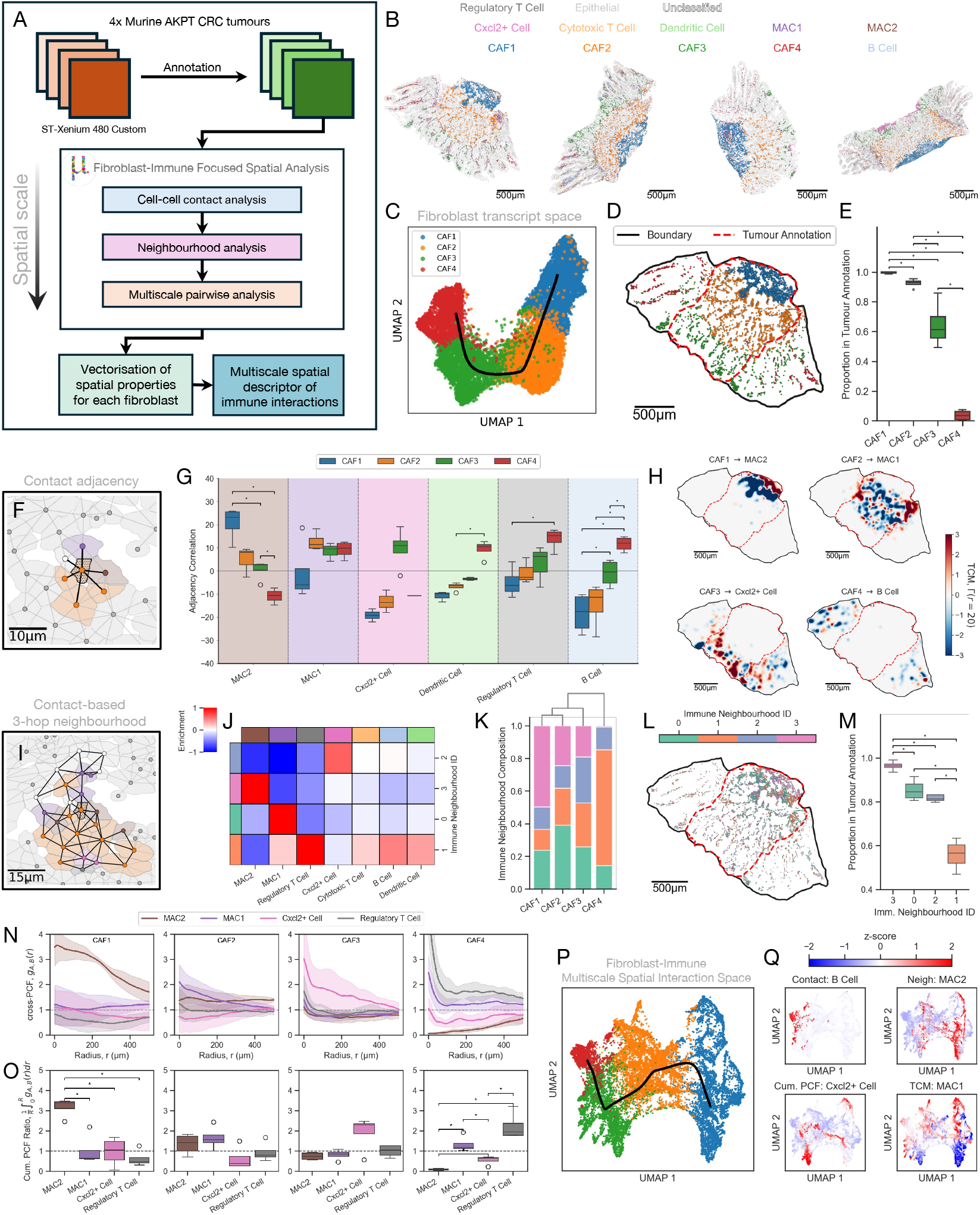
Multiscale spatial correlation of fibroblasts and immune cells in murine AKPT CRC tumours. A: Spatial analysis workflow overview. B: Cell boundaries in the murine AKPT colorectal tumour samples, coloured by cell type. C: UMAP projection of fibroblast transcriptional profiles (*n* = 16, 512). Black line represents inferred trajectory through transcript clusters. D: Fibroblasts cell centres in a representative tumour sample, with sample boundary (black) and pathologically annotated tumour region (red dashes). E: Proportion of fibroblasts within the tumour annotation across all samples (*: *p <* 0.05, Mann-Whitney). F-H: Cell-cell contact analysis. F: 1-hop neighbourhood on segmented cell boundaries (connectivity distance *<* 1.5µm), coloured by cell type. G: Adjacency permutation test (APT) results describing fibroblast correlation with immune cell subtypes (*: *p <* 0.05, Mann-Whitney). H: Topographical correlation maps (TCMs) quantifying short range (*<* 20µm) colocalisation between fibroblasts and immune cells. I-M: Neighbourhood analysis. I: 3-hop neighbourhood on cell boundaries (connectivity distance *<* 1.5µm), coloured by cell type. J: Enrichment matrix for immune cells in 3-hop neighbourhoods centred around a fibroblast (*k*-means clustering, *k* = 4, square-root-transformed *z*-scores). K: Distribution of immune neighbourhoods around each fibroblast subtype, coloured by neighbourhood ID. L: Fibroblast cell centres coloured by immune neighbourhood ID (annotations as in D). M: Proportion of fibroblasts with each immune neighbourhood classification within the tumour annotation (*: *p <* 0.05, Mann-Whitney). N-O: Multiscale analysis of fibroblast-immune spatial relationships. N: Cross-PCFs between selected fibroblast-immune cell pairs (mean, 95% confidence intervals). O: Cumulative PCF ratios summarising the first 150µm of each cross-PCF, 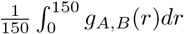, for each fibroblast–immune cell pair (*: *p <* 0.05, Mann-Whitney). P: UMAP projection summarising multiscale spatial immune interaction metrics across all fibroblasts, coloured by fibroblast type (line represents inferred trajectory). Q: UMAP projection coloured by four selected spatial interaction features (*z*-score): 1-hop adjacency with B cells, abundance of Macrophage Type 2 cells in 3-hop neighbourhoods, cumulative cross-PCF ratio with Cxcl2+ Cells and TCM values with Macrophage Type 1 cells.

Cell annotation identified four transcriptionally distinct cancer-associated fibroblast subtypes, spanning a trajectory from fibrotic CAF1 to myofibroblast-like CAF4 [38] (Fig. 6C). Their spatial distributions reflected this progression: CAFs 1–3 localised inside annotated tumour regions, while CAF4 cells were enriched outside the tumour periphery (Figs. 6D–E).

At the cell–cell contact scale, proximity networks (≤ 1.5µm) defined 1-hop fibroblast neighbourhoods (Fig. 6F). Adjacency Permutation Tests (APT) revealed subtype-specific interactions: CAF1 fibroblasts with Spp1+ macrophages (MAC2), CAF3 with Cxcl2+ cells, and CAF4 with B cells and dendritic cells (Fig. 6G; Fig. E10I). Topographical Correlation Maps (TCM, *r <* 20µm) highlighted regional variation in cell interactions: for example, CAF2-MAC1 contacts were concentrated at the invasive front, but absent in the tumour bulk (Fig. 6H, Fig. E10H).

At the neighbourhood scale (≈ 60µm), 3-hop immune environments around fibroblasts were clustered by immune cell composition, yielding four neighbourhoods dominated by MAC1, MAC2, Cxcl2+ cells, or a lymphoid mixture including regulatory T cells (Tregs) respectively (Figs. 6I–J; Appendix E). CAF1 cells were surrounded by MAC2-dense neighbourhoods (51%), while CAF4 cells were strongly associated with Treg cell environments (67%) (Fig. 6K). Spatial differences in the localisation of the different immune neighbourhoods were evident: macrophage- and Cxcl2+-rich neighbourhoods were localised within the tumour mass, while adaptive cell-enriched neighbourhoods mapped to the periphery (Figs. 6L–M).

At the tissue scale (≈ 500µm), cross-PCFs confirmed these associations: CAF1 cells positively correlated with MAC2 cells, and were excluded from Treg cells, whereas CAF4 cells co-localised with Tregs and were excluded from MAC2 cells (Figs. 6N–O).

To integrate findings across scales, we constructed an 87-dimensional summary vector for each fibroblast, consisting of vectorised metrics describing adjacency, neighbourhoods, and correlation (see Methods). There was clear segregation of fibroblast subtypes in this vector space, with spatial association profiles aligning with transcriptional states (Fig. 6P). Trajectory inference further supported a continuum of fibroblast–immune interactions, mirroring the transcriptional trajectory.

This analysis demonstrates MuSpAn’s capacity to reveal structured stromal–immune organisation across spatial scales. Transcriptionally and spatially distinct cancer associated fibroblast populations generate discrete immune niches, with CAF1 located in central tumour regions which are macrophage-rich and CAF4 cells concentrated in adaptive immune cell-enriched regions outside the tumour periphery. These patterns recapitulate hallmark features of CMS4 tumours, including immunosuppressive macrophage enrichment, fibroblast heterogeneity, and compartmentalised immune exclusion. By integrating spatial data from direct contact to tissue-wide co-localisation, MuSpAn uncovers coordinated stromal–immune organisation, reinforcing stromal architecture as a driver of immune evasion and therapy resistance [39, 40].

### Enabling reproducible and customisable spatial analysis

MuSpAn provides a flexible toolbox which can be applied in many ways and contexts. Central to this is MuSpAn’s ability to easily reproduce existing data analysis pipelines, proposed for specific biological contexts and data types. Further, MuSpAn makes extending or adapting such pipelines trivial. This is demonstrated in the case studies in the preceding sections, which have adapted existing pipelines for exploratory data analysis (in the healthy tissue example) and for biologically-driven pipeline development (in the AKPT tissue example). Further case studies are presented in the Supplementary Information (Appendix A), where we reproduce previous data analysis workflows on datasets from COVID-19 lung tissue, head and neck cancer, and colorectal cancer tissue microarrays. These examples further demonstrate the platform agnostic nature of MuSpAn, with datasets from co-registered IHC slides, CODEX, and imaging mass cytometry. The flexibility of MuSpAn’s “library” approach differs from commonly used “pipeline” approaches to spatial data analysis, and, as such, care must be taken to ensure that pipelines constructed using MuSpAn are relevant to the research question of interest. An in-depth discussion of key factors to consider when developing an effective biologically-driven pipeline is present in the Supplementary Information (Appendix B).

## Discussion

Spatial biology enables precise mapping of cell populations, their interactions, and structural arrangements within complex tissues. By quantifying intercellular organisation, spatial analysis has become a powerful driver of biological discovery. Yet, the diversity of imaging modalities, scales of analysis, and analytical frameworks makes it difficult to extract consistent biological insights. Existing tools often specialise in single imaging platforms or narrow questions, leaving researchers without flexible options to compare or integrate methods across scales.

MuSpAn addresses this gap by offering a versatile, multiscale platform for spatial analysis in a modality-agnostic manner, compatible with diverse imaging and omics technologies. It provides streamlined access to over 100 methods drawn from geometry, topology, network theory, spatial statistics, optimal transport, and probability. By allowing analyses at one scale to inform those at another, MuSpAn helps quantify biological complexity that may otherwise be overlooked. Here, we have demonstrated the application of MuSpAn for analysis of spatial datasets from the subcellular level through to tissue structural organisation in mouse normal and neoplastic intestinal datasets. We have highlighted the convergent use of diverse mathematical tools to identify and quantify cell morphology, distribution and cross-compartmental interactions, and demonstrated the power of this for biological discovery.

MuSpAn emphasises flexibility, supporting both detailed exploratory analyses in discovery biology and scalable, high-throughput workflows. Users can easily benchmark and cross-validate complementary methods within the same dataset, overcoming the limitations of relying on any single analytic approach and providing confidence that identified cell associations are genuine. Comprehensive online resources, including documentation, detailed tutorials, and scientific case studies, support spatial data exploration and foster community engagement (https://www.muspan.co.uk). MuSpAn’s user-focussed design permits most statistics to be executed and visualised with a single line of code, while its *query* system is intuitive for routine use while enabling highly customisable spatial data filtering for advanced investigations. These features combine to maximise flexibility and foster innovation, empowering researchers to rapidly develop and refine bespoke spatial analysis pipelines.

Beyond exploratory analysis, MuSpAn complements the growing role of AI in spatial biology. While AI excels at pattern recognition, it often lacks interpretability [41]. MuSpAn delivers directly interpretable metrics that provide mechanistic insight (e.g., describing cell–cell contact, exclusion zones, morphology). Metrics can be vectorised for integration into AI workflows, bridging interpretable quantification and predictive modelling (an approach successfully applied in TDA [42, 43]).

Looking ahead, MuSpAn is positioned to address challenges in integrating spatial data with other modalities including single-cell RNA sequencing, as well as with patient metadata to link spatial patterns to clinical outcomes. Its architecture supports future extensions to 3D and longitudinal datasets, enabling spatio-temporal analyses that capture dynamic tissue reorganisation. By combining accessibility, mathematical depth, and multiscale capability, MuSpAn bridges the gap between advanced imaging technologies and interpretable analytics, and complements the wider community of tools for exploratory spatial biology [14, 20, 32, 44]. Alongside these tools, MuSpAn empowers researchers to move beyond qualitative mapping toward mechanistic, quantitative, and multiscale biological insight.

## Methods

### MuSpAn

#### Overview

MuSpAn provides a general framework for conducting spatial analyses. In this section, we summarise its structure, and explain how this facilitates flexible and targeted spatial analyses which can be applied to any imaging modality or data source. We emphasise that MuSpAn is designed for spatial analysis, and not for preprocessing of raw imaging data. Cell or object identification, region annotation, and labelling should be conducted prior to spatial analysis, usingavailable pipelines or image analysis software (e.g., Xenium explorer, Qupath, Halo, Deepcell). MuSpAn imports array-like data directly within Python — by reading CSV files with libraries like Pandas, or using Numpy arrays - and provides additional ‘quality of life’ tools to import annotations or labels from software such as QuPath or Xenium Explorer.

In the following sections, we outline MuSpAn’s design, key features, and how it enables highly flexible spatial analysis. While many features are not user-facing, and therefore do not require user understanding, understanding the infrastructure supporting MuSpAn can aid more advanced use.

##### Summary

All MuSpAn analyses begin with the *domain*, a class which includes data, labels and metadata. The domain can be viewed as a container to which data is added; this standardises the internal representation of the data so they can be interactively queried and/or analysed by the user. This ensures that analysis methods are not called on incompatible data (e.g., requesting the area of point-like data), that data can easily be filtered for subsequent analysis, and permits straightforward expansion of the spatial analysis toolbox.

Spatial data are stored within the domain as *objects*, which represent distinct biological entities as points or shapes - the interpretation ofobjects can vary. For example, both transcripts and cell centres are represented as point objects, while cell boundaries, manually specified annotations, and region of interest boundaries are shape objects. Each object can have metadata associated as *labels*, which can be used to isolate populations of interest for statistical analysis. By ensuring all data within a domain are held in a standardised format, the data can be easily interrogated using individual quantitative analysis methods, and results visualised within MuSpAn, or returned as raw data for subsequent visualisation or onward use outside of MuSpAn.

##### Domain

A MuSpAn *domain* is a class which standardises the structure of spatial data and metadata, and can be thought of as a container within which objects for analysis are located. Biologically, it represents a single area containing all relevant data for analyses within the same spatial frame of reference, such as a region of interest within an image where all cell (or transcript) locations are represented within the same coordinate system. The domain is enclosed by a continuous closed *boundary*, which defines a simply connected space that fully contains all objects.

When adding new objects to the domain outside of the existing boundary, the boundary of the domain updates by default to be the axis aligned bounding box surrounding all objects; however, the boundary can be easily changed to any non-self intersecting polygon which encloses all objects, such as the convex hull, an alpha shape that surrounds the objects, or an annotation made in external software (for instance, the boundaries of the domains shown in Figs. 2-5 are annotations drawn within Xenium explorer).

The domain also contains helpful metadata, such as colourmaps for visualisation of objects under different criteria (e.g., for colouring objects according to associated labels), or caches to ensure computationally expensive calculations such as distances between objects or network representations of the data can be retained where needed.

##### Objects

MuSpAn *objects* represent biological entities as either *points* or *shapes*, each assigned a unique Object ID. Points are defined by (*x, y*) coordinates, while shapes are non-self-intersecting polygons defined by ordered coordinate lists. Shapes can include complex features like internal voids, allowing them to represent diverse regions such as cell boundaries, tumour zones, or areas for exclusion in spatial analyses.

Each object’s Object ID links it to metadata, including labels, spatial properties, networks, distances, and other features. Objects can be added or removed from a domain and may store pre-calculated attributes, such as area, perimeter, bounding circles for efficient distance calculations, or relationships to other objects (e.g., parent-child links created through transformations like shape decomposition). The spatial distance between two objects is defined as the minimum Euclidean distance, accommodating both point-like and shape-like objects. For shapes, distances can be calculated using the centroid or via the nearest part of the relevant line segment in the shape boundary.

Objects can belong to one or more collections, which are named subsets of objects (e.g., ‘cell boundaries’, ‘transcripts’, or ‘damaged tissue’). A collection denotes a subset of objects, and is one way of referring to multiple objects at once. Collections have no restrictions on names or overlapping membership.

##### Labels

The main mechanism by which MuSpAn associates metadata with an object is through the use of *labels*. Each object may be assigned any number of different labels, which may be categorical (discrete) or numeric (continuous). For example, in Fig. 2D-G, cells are shape objects that have been assigned a categorical label ‘Celltype 1’ or ‘Celltype 2’, as well as numerical labels ‘Area’, ‘Principal angle’ and ‘Circularity’. Labels can be assigned to some subset of objects (for instance, all objects in one collection), either based on metadata obtained from outside of MuSpAn (e.g., labelling cells from multiplex immunohistochemistry according to their mean pixel intensity of a marker of interest) or from within MuSpAn (e.g., labelling cells according to the result of some spatial analysis of interest). In Fig. 2D-G, the celltype label is based on the preprocessing steps conducted in Xenium explorer, while the other labels are derived from geometric summary statistics calculated for each cell within MuSpAn and subsequently added to the shapes. Labels can be used for analysis or visualisation, or as part of the calculation of other spatial metrics.

##### Moving between spatial scales

As indicated in Fig. 1D, a key way of moving between spatial scales in MuSpAn is by defining new objects based on existing ones. Starting from the lowest spatial scale added to the domain (in our examples, point objects representing transcript locations), we can move up spatial scales by combining transcript locations into a single cell object (using the segmentation mask if available, or by finding, for instance, the convex hull or *α*-shape of the points). Converting a shape into a single point (e.g., turning a cell boundary into a cell centroid) also moves up spatial scales by ‘simplifying’ a collection of more complex spatial information into a single object. These processes effectively lower the resolution of the object or objects in question (moving from a set of points to a single shape, or from a detailed shape boundary to a single centroid), which can then be considered in the context of other objects at the same spatial scale without becoming computationally infeasible.

Data can be transferred across spatial scales through the use of labels, which allow a ‘coarse grained’ object to contain information about the ‘fine grained’ objects from which is has been derived - for instance, a cell centroid could be labelled according to the circularity or area of the parent cell boundary object. This allows features computed at one scale to be inherited by the new ‘child’ objects derived from the ‘parent’. Parent-child relationships are stored within the domain as new objects are created, which allows easy construction of ancestry trees for multiscale information retrieval.

##### Queries

MuSpAn provides powerful query tools to filter and analyse objects based on specific properties, enabling both straightforward and advanced data interrogation. While most users benefit from this infrastructure automatically, advanced users can directly create and use queries to extract complex data subsets using the `query` module. A query in MuSpAn specifies criteria for selecting objects. It comprises three components:

1. a property of interest (e.g., the label ‘Celltype’);
2. a relation (e.g., ‘is equal to’);
3. a comparison value (e.g., ‘T cell’).

These components would generate a query that returns any objects with a ‘Celltype’ label equal to ‘T cell’. This query is evaluated at runtime to produce a boolean array (i.e., True for matching objects, False otherwise). Once created, queries can be reused across various functions, including spatial analysis and visualisation tools.

Users can construct queries explicitly or rely on MuSpAn’s intuitive syntax for common cases. For example, passing the tuple (‘Celltype’, ‘T cell’) to a function will automatically lead to its interpretation as a query relating to the ‘Celltype’ label. Advanced queries can combine multiple criteria using query containers, which support logical operations (e.g., AND, OR). These containers allow nesting of conditions, enabling highly detailed filtering. For example, a query container might select all T cells within 20 µm of a tumour boundary, or even more complex combinations of spatial relationships, label values, and even summary statistics.

Queries persist in the Python workspace, allowing consistent use across analysis and visualisation methods. While beginner users can rely on MuSpAn’s default behaviour and ‘behind the scenes’ use of the query infrastructure to conduct basic analyses, leveraging the query system provides a powerful way to explore and analyse data efficiently. Tutorials and examples of advanced queries are provided on the MuSpAn website.

##### Visualisation

While not primarily a visualisation package, MuSpAn leverages Python’s matplotlib package [45] to provide easy access to data visualisation tools through MuSpAn’s `visualise` module. In particular, this permits straightforward visualisation of the domain and objects within it, coloured according to labels, collections, or other properties of interest. Domain visualisation accepts query arguments to specify the objects to plot or the metadata used to colour the output, permitting easy publication-ready plotting of complex subsets of objects; almost all the panels in this manuscript are generated using the visualisation tools available within MuSpAn, and detailed tutorials showing how this is achieved are available on the MuSpAn website and alongside this work.

MuSpAn can also be used to natively plot the outputs of selected spatial analysis methods. Every MuSpAn spatial analysis method returns the output as raw data that can be passed by the user to other analysis or visualisation tools; however, this data can also be passed to dedicated functions within the `visualise` module to straightforwardly generate plots. These functions can also be called automatically using the `visualise_output` keyword argument associated with most spatial analysis methods. While the `visualise` module provides customisation via matplotlib, it is generally recommended that users use raw outputs for creating custom visualisation.

### Quantitative analyses

Central to MuSpAn are the quantitative analyses it provides. All of the methods available within MuSpAn v1.2.0 have been previously described in the academic literature, and so are not described in detail here; a complete list of implemented methods, with references to their use in spatial biology where possible, can be found in the Supplementary Information. Where possible, MuSpAn leverages implementations of individual methods from established discipline-specific Python packages [46–48]. The spatial analysis methods provided within MuSpAn are divided between six submodules, named according to the underlying mathematical framework within which the analysis methods are contained. These modules are:

- **Distribution** - containing methods through which spatial data is considered as a continuous distribution, such as a density function.
- **Geometry** - containing methods that describe the morphology or other characteristics of shapes.
- **Networks** - containing methods through which data is represented as spatially embedded networks.
- **Region Based** - containing methods which represent data within spatially-averaged subregions of the region of interest.
- **Spatial Statistics** - containing methods from the field of spatial statistics, which compare observations of point data against statistical null models.
- **Topology** - containing methods from the field of topological data analysis, which describes data in terms of topological invariants.

We reiterate here that each method within these modules is designed to investigate different types of spatial relationships. As such, techniques from any module can be applied at any length scale. To generate pipelines that are robust to potential error introduced by viewing data through the lens of one particular methodology, we suggest that methods from across multiple submodules should be employed where possible (see the Supplementary Information, Appendix B).

#### Specifying populations

Each of the functions available within MuSpAn for spatial analysis accepts different classes of objects; for example, spatial statistics such as the cross-PCF require two point populations to be defined, while a geometric statistic such as convexity is only defined for shape-like (polygonal) objects. Each statistic allows the population (or populations) to be specified when calculated, for instance by using the MuSpAn `query` module described above. This permits subsets of the data to be isolated and analysed very easily, and for the same population to be passed to multiple functions in series, making pipeline construction simple.

#### Specifying analysis boundaries

Many statistics are dependent not only on the population of objects they consider, but also on the boundaries of the analysis. As a simple (non spatial) example, the density of a fixed number of cells varies depending on the size of the region in which they are measured. MuSpAn permits analysis boundaries to be specified via queries in the same way as populations are. Users can specify a set of shapes that define the boundaries of a calculation region, or of shapes defining regions that should be excluded from analysis. MuSpAn converts these requests on-the-fly into simply connected spaces for analysis, and where appropriate adjusts statistical analyses to account for boundary corrections.

Where no boundary is specified, analyses will use the domain boundary. See Appendix A in the SI for examples of spatial analysis within shape boundaries.

### Spatial analysis methods overview

MuSpAn v1.2.0 (used for the analyses in this manuscript) includes a diverse collection of 40+ quantitative analysis methods which have been described in the scientific literature. Table 1 summarises the spatial methods used in the exemplar analyses shown in Figures 2–6, and provides references containing detailed methodological descriptions (for additional information see the SI, Appendix D). A complete list of all spatial analysis methods implemented in MuSpAn v1.2.0 is provided in the SI (Appendix C). In addition, detailed information on all 150+ MuSpAn functions can be found at https://docs.muspan.co.uk.

**Table 1:** Overview of spatial analysis methods featured in Figures 2–6. We indicate each method’s module within the MuSpAn package and provide a concise description. For full details on the definitions of these methods, please see the references provided. References for standard methods are omitted (denoted by ‘-’).

| Figure | Method<br>(MuSpAn module) | Summary | Ref |
| --- | --- | --- | --- |
| 2 | Circularity<br>(`geometry`) | A measure of how similar the shape of a polygon is to a circle, with values $\approx 1$ indicating shapes that are closer to perfectly round. | - |
| 2 | Principal angle<br>(`geometry`) | The angle of the longest line through the center of a polygon that best represents its overall orientation, often used to describe a shape’s direction or alignment. | [49] |
| 2 | Label composition<br>(`summary_statistics`) | The count of a specific label category, normalised relative to the total number of labels in the region. | - |
| 3 | Average nearest neighbour index (ANNI)<br>(`spatial_statistics`) | A statistical measure used to determine whether a spatial distribution of points is clustered, random, or dispersed by comparing observed and expected nearest neighbour distances. | [34] |
| 3 | Proximity network<br>(`networks`) | A network that connects nodes based on their spatial distance. | [50] |
| 3 | Quadrat correlation matrix (QCM)<br>(`spatial_statistics`) | A measure of correlation between the counts of different cell types within a set of regions. | [51] |
| 4 | Cross pair correlation function (cross-PCF)<br>(`spatial_statistics`) | A measure of spatial aggregation between pairs of objects across length scales. | [52] |
| 4 | Vietoris-Rips filtration<br>(topology) | A method in topological data analysis that constructs a nested sequence of simplicial complexes by connecting points within increasing distances, capturing the shape and features of a dataset across scales. | [46] |
| 5 | Delaunay network<br>(networks) | A network where nodes in a plane are connected to form triangles such that no point lies inside the circumcircle of any triangle. | [50] |
| 5 | $k$-hop neighbourhood<br>(networks) | The set of all nodes that can be reached from a given node within $k$ steps or connections. | - |
| 5 | Neighbourhood clustering<br>(networks) | Group similar neighbourhoods together based on the enrichment of labels within the neighbourhoods. | [50] |
| 5 | Adjacency permutation test<br>(networks) | A statistical method that assesses the significance of node label adjacency by randomly permuting node labels and comparing the observed property to the distribution from the permutations. | [53] |
| 5 | α-shape<br>(helpers) | A method to define the shape of a set of points in space by connecting points that are close to each other, using a parameter $\alpha$ to determine the length scale captured. | - |
| 5 | Hexgrid<br>(region.based) | Generate a mesh of regular hexagons covering the requested space. | - |
| 5 | Local Getis-Ord*<br>(spatial.statistics) | A statistic that identifies spatial clusters by measuring the degree of local spatial association for a variable, highlighting areas of high or low values relative to their surroundings. | [54] |
| 5 | Kernel density estimation (KDE)<br>(distribution) | A non-parametric statistical method for estimating the probability density function of a random variable by smoothing data points using a kernel function. | [47] |
| 5 | Wasserstein distance<br>(distribution) | A metric for comparing probability distributions by measuring the optimal transport cost required to move one distribution to another. | [28] |
| 5 | KL-divergence<br>(distribution) | A measure of the difference between two probability distributions by quantifying the amount of information lost when approximating one distribution with another. | [55] |
| 6 | Topographical correlation map (TCM)<br>(`spatial_statistics`) | A Local Indicator of Spatial Association (LISA) which quantifies the pairwise correlation between two populations of points and is visualised locally as a heatmap. | [56] |
| 6 | Ripley’s K-function<br>(`spatial_statistics`) | A measure of pairwise spatial correlation between two populations of points, based on comparing the observed numbers of points within a minimum pairwise distance of one another. | [52] |
| 6 | Empty space function<br>(`spatial_statistics`) | A function which characterises empty spaces by repeatedly sampling random locations and measuring the distance to the nearest point. | [52] |

### Datasets

#### Healthy murine colon - Xenium

Spatial analysis was conducted on murine colon tissue stained with the Xenium Mouse Tissue Atlas Panel (10x Genomics, USA), using dataset ‘Fresh Frozen Mouse Colon with Xenium Multimodal Cell Segmentation’ sourced from the 10x Genomics website and available under a CC BY license (available from https://www.10xgenomics.com/datasets/fresh-frozen-mouse-colon-with-xenium-multimodal-cell-segmentation-1-standard). Fresh-frozen tissue arrays were prepared from 8-week-old male C57 mice (Charles River Laboratories, USA). Tissue samples were processed according to the Xenium protocols for fresh-frozen tissues, following the Xenium In Situ for Fresh Frozen Tissues - Tissue Preparation Guide (CG000579) and Fixation & Permeabilisation Guide (CG000581). Staining procedures involved probe hybridisation, washing, ligation, amplification, and cell segmentation staining, carried out in alignment with the Xenium In Situ Gene Expression with Cell Segmentation Staining User Guide (CG000749). Post-instrument processing, including hematoxylin and eosin (H&E) staining, followed the Xenium In Situ Gene Expression - Post-Xenium Analyser H&E Staining Protocol (CG000613). The Xenium Mouse Tissue Atlas Panel, pre-designed by 10x Genomics, includes 379 genes selected to represent major cell types and optimised for spatial gene expression profiling in mouse tissues.

Transcript clustering was provided with the Xenium experiment, which assigned 25 distinct cluster IDs to cells using a graph-based approach (Leiden clustering). Cluster IDs 1-19 were considered in our demonstrative analysis due to low observations of cells with Cluster IDs 20-25. The 19 retained clusters were subsequently annotated on the basis of the differential expression matrix provided by 10x Genomics (Table 2).

**Table 2:**
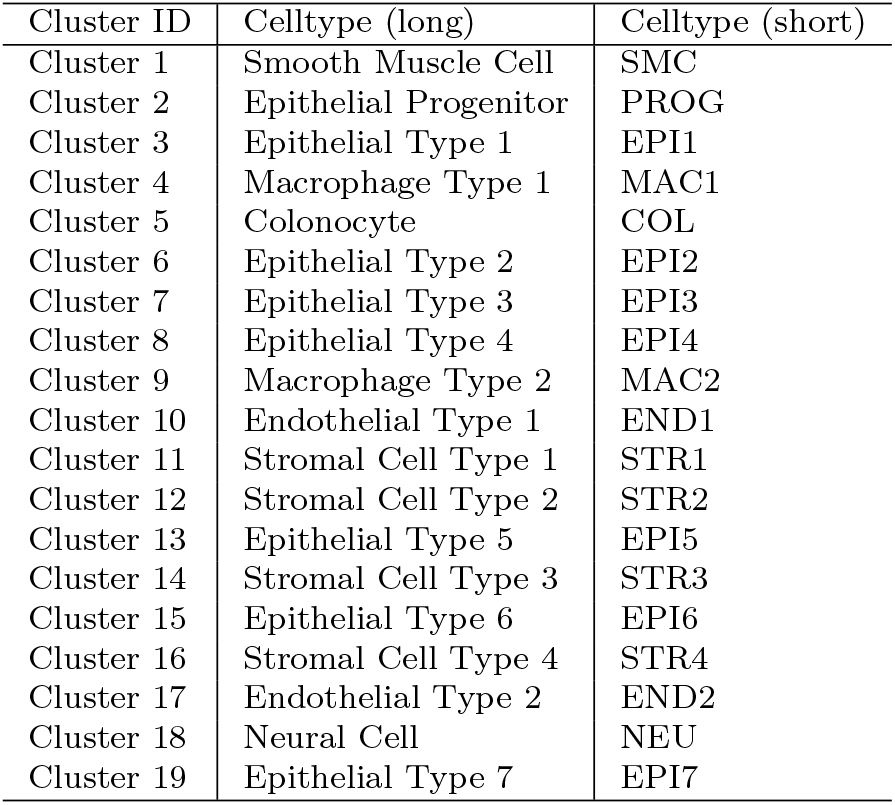
Transcript-defined cell clusters with corresponding full and abbreviated cell type annotations from the healthy murine colon Xenium dataset.

| Cluster ID | Celltype (long) | Celltype (short) |
| --- | --- | --- |
| Cluster 1 | Smooth Muscle Cell | SMC |
| Cluster 2 | Epithelial Progenitor | PROG |
| Cluster 3 | Epithelial Type 1 | EPI1 |
| Cluster 4 | Macrophage Type 1 | MAC1 |
| Cluster 5 | Colonocyte | COL |
| Cluster 6 | Epithelial Type 2 | EPI2 |
| Cluster 7 | Epithelial Type 3 | EPI3 |
| Cluster 8 | Epithelial Type 4 | EPI4 |
| Cluster 9 | Macrophage Type 2 | MAC2 |
| Cluster 10 | Endothelial Type 1 | END1 |
| Cluster 11 | Stromal Cell Type 1 | STR1 |
| Cluster 12 | Stromal Cell Type 2 | STR2 |
| Cluster 13 | Epithelial Type 5 | EPI5 |
| Cluster 14 | Stromal Cell Type 3 | STR3 |
| Cluster 15 | Epithelial Type 6 | EPI6 |
| Cluster 16 | Stromal Cell Type 4 | STR4 |
| Cluster 17 | Endothelial Type 2 | END2 |
| Cluster 18 | Neural Cell | NEU |
| Cluster 19 | Epithelial Type 7 | EPI7 |

#### AKPT murine colorectal cancer - Xenium

We utilised the *Villin-CreER*^*T2*^*;Apc*^*fl/fl*^*;Kras*^*G12D/+*^*;Trp53*^*fl/fl*^*;Tgfβr1*^*fl/fl*^ (*n* = 2) or *Villin-CreER*^*T2*^*;Apc*^*fl/+*^*;Kras*^*G12D/+*^*;Trp53*^*fl/fl*^*;Tgfβr1*^*fl/fl*^ (*n* = 2) (AKPT) murine model of colorectal cancer. Driver alleles were initiated as follows: A is *Apc*^*fl/fl*^, K is *Kras*^*G12D*^, P is *p53*^*fl/fl*^, T is *Tgfβr1*^*fl/fl*^. This combination recapitulates the genetic and microenvironmental features of aggressive, stromal rich, colorectal cancer.

Gut preparations were washed in PBS, fixed overnight in 10% neutral buffered formalin and then transferred to 70% ethanol before processing for embedding. Formalin-fixed gut sections were rolled into Swiss Rolls, pinned and placed in a histology cassette. Specimens were processed using a Histomaster machine (Bavimed). Processed samples were embedded in paraffin wax using a paraffin embedding station (EG1150H, Leica). 5µm formalin-fixed, paraffin-embedded tissue sections were used on Xenium slides. Slides were baked at 42^∘^C for 3 hours. The sections stained following the manufacturers protocol with a custom 480 panel of genes with Xenium v1 chemistry.

Seurat (v5.2.0) was used in R (v4.3.3) to annotate four AKPT murine CRC Xenium samples using transcript profiles, which were merged, normalised, and clustered, with epithelial-rich clusters identified by Epcam and Cdh1 expression and removed. The remaining non-epithelial cells were reclustered at higher resolution to identify fibroblast (*Pdgfra*+) and immune (*Ptprc*+) populations, with classification based on scaled aggregate gene expression across 22 clusters. Immune clusters were further subtyped using a marker-based heatmap (ComplexHeatmap v2.18.0), resulting in a low-resolution annotation of 12 immune cell populations. For further information on sample annotation, see Appendix E.

### Multiscale spatial profiling of fibroblast-immune interactions in colorectal tumours

To investigate fibroblast–immune interactions across spatial scales, we applied a series of spatial statistical and network-based analyses using the MuSpAn. For each fibroblast type of interest, we quantified spatial context of immune interactions at three spatial scales: (i) direct cell–cell contact, (ii) neighbourhood composition, and (iii) multiscale (0-500µm) spatial association.

At the cell-cell contact level, we built proximity networks from cell boundaries (contact distance ≤1.5µm) to define immediate (1-hop) adjacency immune cells and computed an *Adjacency Permutation Test* (APT) to identify correlations between neighbouring fibroblasts and immune cells. We also computed *Topographical Correlation Maps* (TCMs) to quantify the spatial significance of resultant co-localised pairs from the APT. At an intermediate scale, extended (3-hop) neighbourhoods centred on the selected fibroblasts in the proximity network were computed and clustered by immune composition using *k*-*means* with *k* = 4, selected to yield distinct and interpretable immune microenvironments around each fibroblast. For larger length scales, we used the *cross–pair correlation function* (cross-PCF) to quantify spatial associations between fibroblast and immune populations from 0 to 500µm.

The above measurements were aggregated into an 87-dimensional feature vector for each fibroblast across all samples. For each fibroblast, the features include:

- Number of direct cell–cell contacts with each immune subtype (7);
- Number of cells within a 3-hop contact neighbourhood with each immune subtype (7);
- TCM value at the fibroblast’s location with respect to each immune subtype at *r* = 30µm (7)
- Contribution of fibroblast to the respective cross-PCF from all fibroblasts to each immune and fibroblast subtype at 0, 100, 200, 300, 400µm (55);
- Cumulative cross-PCF ratio up to *R* = 150µm for each immune and fibroblast subtype (11).

Feature vectors were *z*-score normalised and summarised medians were computed for each fibroblast type. Pairwise distances between fibroblast type median summary vectors were used to construct a weighted minimum spanning tree (WMST), which was projected into UMAP space to visualise an inferred trajectory of fibroblast–immune states. For an extended description of methods, see Appendix E.

## Code Availability

MuSpAn is freely available for non-commercial use via the MuSpAn website (https://www.muspan.co.uk/get-the-code) under an Academic Use Licence. Commercial users should contact Oxford University Innovation via their software store (https://process.innovation.ox.ac.uk/software/p/21795/multiscale-spatial-analysis-toolbox/1). Comprehensive user guides including installation, documentation, and detailed tutorials are available via https://docs.muspan.co.uk. Scripts that reproduce the analysis conducted in this paper can be found at https://github.com/joshwillmoore1/Supporting_material_muspan_paper. A public Github repository for issues tracking, feature suggestion, and community discussion, is available at https://github.com/joshwillmoore1/MuSpAn-Public.

## Data Availability

The ‘healthy mouse colon’ dataset is available from 10x Genomics (https://www.10xgenomics.com/datasets/fresh-frozen-mouse-colon-with-xenium-multimodal-cell-segmentation-1-standard). The synthetic dataset used in Fig. B9 can be loaded within MuSpAn via the command `muspan.datasets.load_example_domain(‘Synthetic-Points-Architecture’)`. All MuSpAn domains generated and used can be found at https://doi.org/10.5281/zenodo.17176282.

## Acknowledgements

The authors would like to thank the many researchers (experimentalists, bioinformaticians, mathematicians, and others) involved in testing the MuSpAn package for their valuable feedback in implementing improvements in the usability and performance of the package. Many thanks to Ling-Pei Ho, Praveen Weeratunga, Chris Pugh, and Philip Macklin for collaborations which led to data shown in Section A of the Supplementary Information. The authors would also like to thank Oxford University Innovation for their help in releasing MuSpAn. JAB and JWM were supported by Cancer Research UK (CRUK) grant number CTRQQR-2021 \ 100002, through the Cancer Research UK Oxford Centre. SJL was supported by CRUK Program Grant (DRCNPG-Jun22 \ 100002) and the MRC Mouse Genetics Network. EJM was supported by the Lee Placito Medical Research Fellowship (University of Oxford). For the purpose of open access, the authors have applied a CC BY public copyright licence to any author accepted manuscript arising from this submission.

## Author Contributions

JAB and JWM were responsible for software development and implementation. JAB, JWM, HMB, and SJL conceptualised the project. SMC, ML, HLB-D and EJM-I were responsible for data curation. All authors wrote and reviewed the manuscript, and approved the final version.

## Appendix A Application of MuSpAn to other imaging modalities

We demonstrate how MuSpAn can be used to reproduce and extend existing data analysis pipelines across diverse imaging modalities. To illustrate its versatility, we present three case studies:

- Head and neck cancer tissue, imaged via stacked, registered IHC slides [S1];
- COVID-19 infected lung tissue, imaged with Imaging Mass Cytometry [S2];
- Colorectal cancer tissue microarrays, imaged using the CODEX platform [S3].

In each case, MuSpAn replicates published analyses and augments them with additional methods (Fig. A1). Note that each pipeline could be indefinitely extended straightforwardly using alternative MuSpAn components; these examples are chosen to complement the methods used in previous studies. Code is available in the Supplementary Information GitHub repository: https://github.com/joshwillmoore1/Supporting_material_muspan_paper. Further examples, including applications to Visium data, are provided on the MuSpAn website: https://docs.muspan.co.uk/latest/tutorials.html.

### A.1 Macrophage localisation in head and neck cancer (IHC)

We use MuSpAn to analyse stacked IHC images from a resected human head and neck cancer sample [S1, S4] (Fig. A2). The dataset comprises four consecutive tissue slices stained with PanCK (epithelial cells), CAIX (hypoxia), pimonidazole (severe hypoxia), and CD68 (macrophages) (Fig. A2A). These slides were registered to form a single composite image (Fig. A2B). The ROI used in this analysis is available as an exemplar domain within MuSpAn and can be loaded via `muspan.datasets.load_dataset(‘Macrophage-Hypoxia-ROI’)` (Fig. A2C). Macrophage centres are represented as point objects, while PanCK, pimonidazole, and CAIX boundaries define shape objects.

In the original study [S1], the pair correlation function (PCF) and J-function were used to assess macrophage infiltration into unannotated tumour nests. Using MuSpAn, we replicate these analyses and extend them by using MuSpAn’s `include_boundaries` and `exclude_boundaries` keywords to compare stroma and tumour regions. Additionally, we compute kernel density estimates (KDEs) to visualise macrophage density, both across the entire ROI and within separate tissue compartments.

**Fig. A1:**
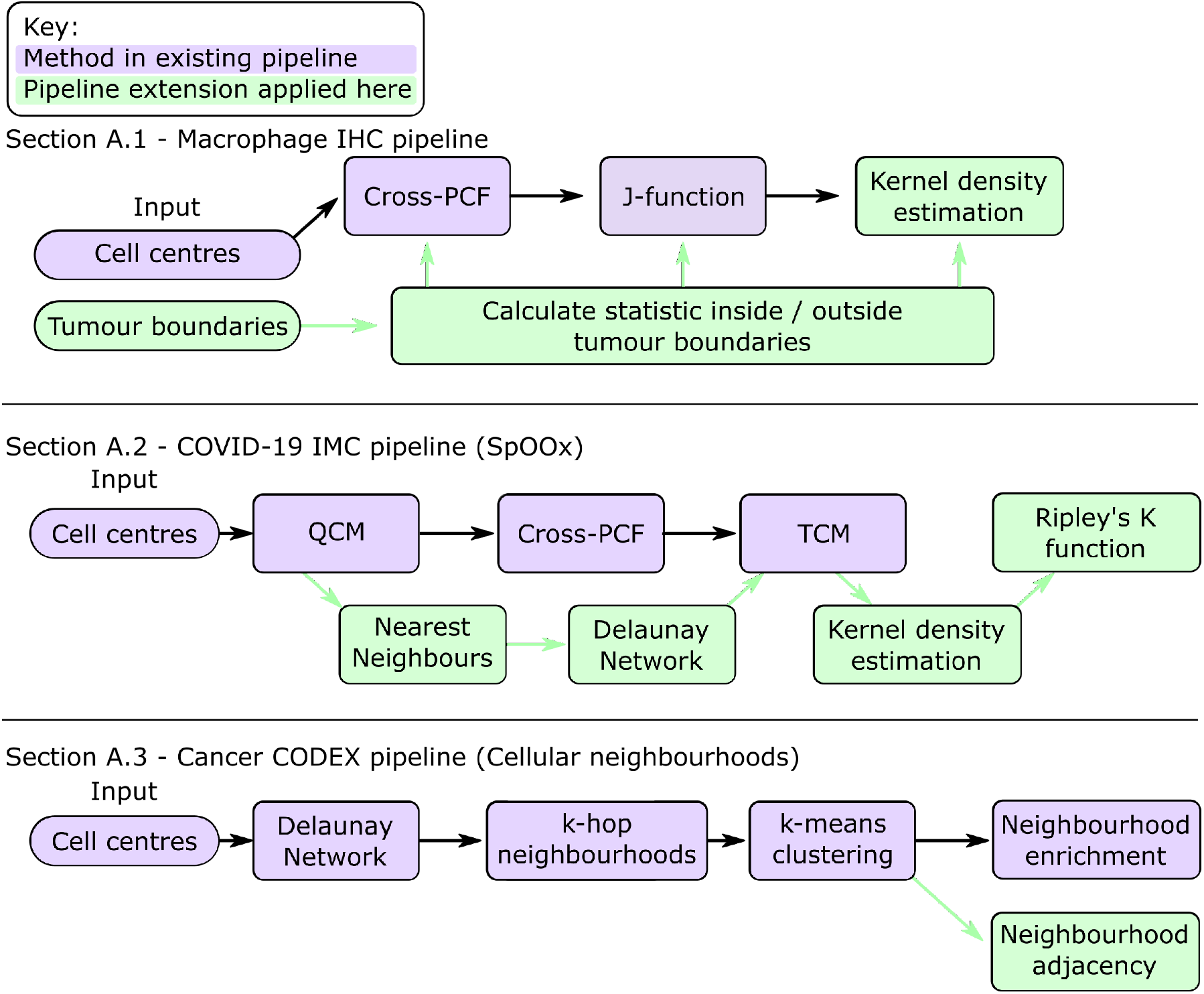
Case studies using MuSpAn to reproduce and extend existing pipelines. Spatial quantification components of three previously published pipelines used for spatial data analysis (A.1: [S1]; A.2: [S2]; A.3: [S3]). Key: Purple boxes - existing analyses. Green boxes - methods used to extend each pipeline here.

We first demonstrate that MuSpAn reproduces the spatial statistics used in the original analysis, specifically the macrophage-macrophage cross-PCF and J-function (Fig. A2D-E). Calculating the cross-PCF across the entire ROI (blue dashed line) reveals short-range exclusion - reflecting the physical spacing between adjacent macrophage cell centres - and clustering at distances beyond around 20 pixels (17.6µm). At this length scale, macrophages are approximately 2.25 times more likely to be found near another macrophage than if they were randomly distributed.

In the original pipeline, the ROI was treated as a uniform square without distinguishing tumour and stroma regions. Using MuSpAn’s `include_boundaries` and `exclude_boundaries` keywords, we extend the analysis to account for these compartments. Both the cross-PCF and J-function show stronger clustering within tumour regions (green line) compared to stroma (red line). Specifically, two stromal macrophages are around 1.75 times more likely than random to be 20µm apart, while tumour macrophages are around 2.75 times more likely - an insight not accessible via the original pipeline. MuSpAn automatically adjusts statistical calculations to account for edge effects introduced by the new boundaries.

We further extend the analysis by computing KDEs to visualise macrophage density across different compartments (Fig. A2F-H). The global KDE (Fig. A2F) highlights a dense macrophage cluster in the upper right of the domain. When restricted to the stroma (excluding PanCK shapes), this cluster dominates the density map (Fig. A2G). In contrast, the KDE within tumour regions (PanCK shapes) reveals that the tumour island with the highest macrophage density is in fact in the lower left (Fig. A2H), which is obscured in the global view.

**Fig. A2:**
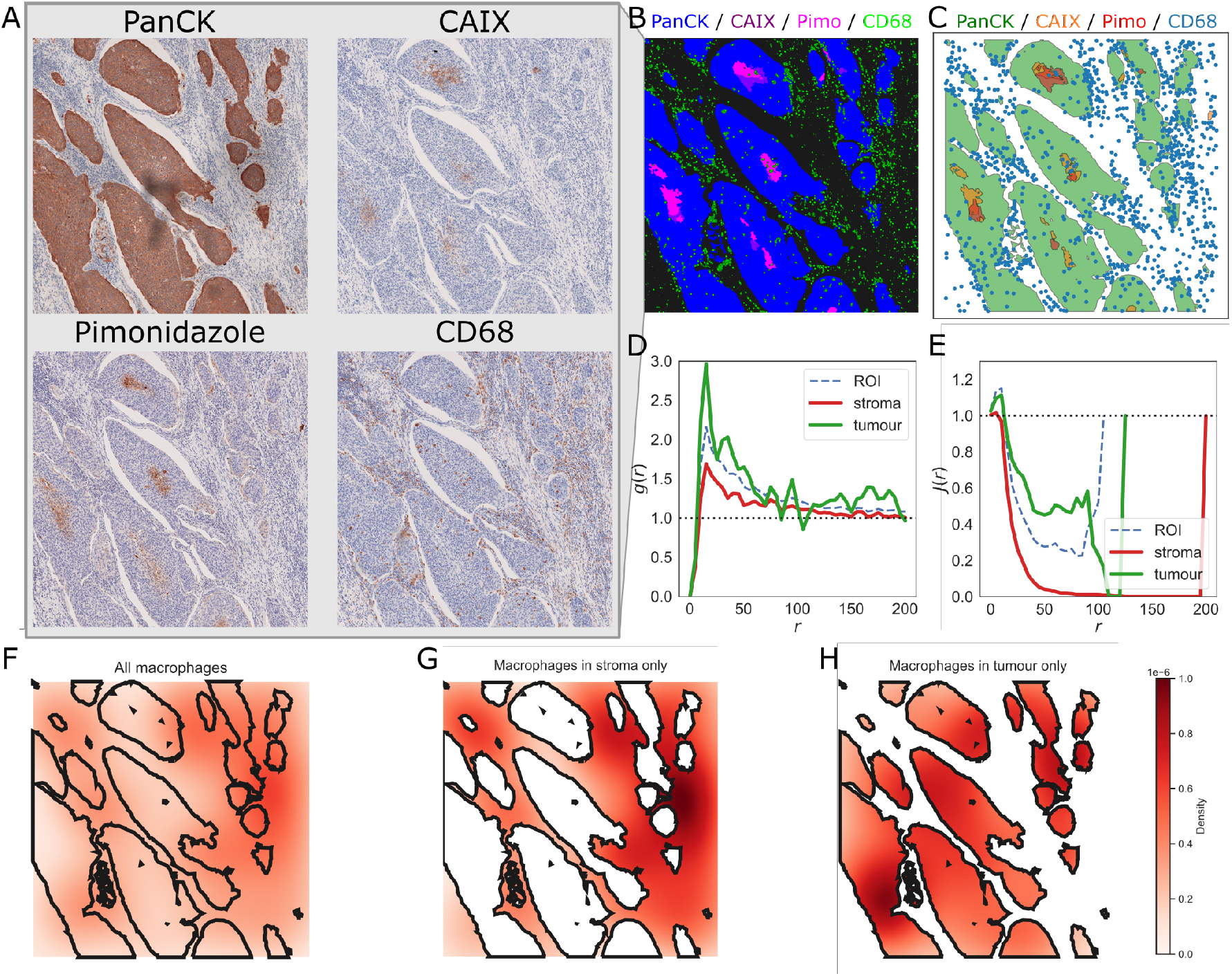
MuSpAn applied to stacked IHC data. A: A 1700 *×* 1700 pixel (1.5mm *×* 1.5mm) ROI from a human head and neck cancer sample, stained on consecutive slides in the tissue stack to show PanCK (tumour cells), CAIX (hypoxia), pimonidazole (severe hypoxia) and CD68 (macrophages). B: Composite image showing the alignment of the four markers. C: Boundaries and points from panel B shown as a MuSpAn domain. D, E: Cross-PCFs (D) and J-function (E) describing macrophage-macrophage correlation within the domain. The dashed blue lines (ROI) show macrophage-macrophage correlation across the whole ROI. The solid green (tumour) and red (stroma) lines show macrophage-macrophage correlation, restricted to inside the PanCK regions (tumour) or outside (stroma). F: Kernel density estimation (KDE) of macrophage localisation within the ROI. Outlines of PanCK shapes are shown for reference, but not used in the estimation. G: KDE of macrophage localisation outside of the PanCK (tumour) regions. H: KDE of macrophage localisation within the PanCK (tumour) regions.

### A.2 COVID-19 infected lung (Imaging Mass Cytometry)

We analyse a 2mm *×* 2mm ROI from an Imaging Mass Cytomtery (IMC) dataset of COVID-19 infected lung tissue, histologically classified as ‘Organising Pneumonia’ [S2] (Fig. A3A-C). The original analysis pipeline involved three key steps: computing the QCM to identify cell pairs of interest, followed by cross-PCF and TCM analyses.

Using MuSpAn, we replicate and extend this pipeline. We first compute the QCM across all cell types (Fig. A3D, quadrat size 100µm) and perform an adjacency permutation test (APT) using a Delaunay network with a 20µm cut-off [S5] (Fig. A3E). We focus on two cell pairs: ‘Blood vessels’-‘UD structural SM’ (high QCM correlation) and ‘UD-structural SM’-’Myofibroblasts’ (low QCM correlation). Interestingly, both pairs show positive spatial association in the APT, despite differing QCM scores.

We compute cross-PCFs and cross-*K* functions for both pairs (Fig. A4A-B, Fig. A5A-B). The ‘UD structural SM’-’myofibroblast’ pair shows no short-range correlation and weak anti-correlation at larger scales. To resolve the apparent contradiction in the negative QCM score, positive APT score, and neutral spatial statistics, we extend the analysis with KDEs (Fig. A4C-D, Fig. A5C-D), revealing that while ‘UD structural SM’ cells are widespread, their densest regions do not overlap with ‘myofibroblast’ clusters. Nearest-neighbour analysis (Fig. A4E-F, Fig. A5E-F) confirms that most ‘myofibroblasts’ are near ‘UD structural SM’ cells, but many ‘UD structural SM’ cells are distant from ‘myofibroblasts’. Finally, TCMs (Fig. A4G, Fig. A5G) show that for this cell pair local positive correlation is confined to specific regions, balanced by areas of exclusion, explaining the neutral global PCF.

This example illustrates the need for caution when interpreting spatial statistics in isolation. The QCM suggests negative correlation due to mismatched distributions across the ROI, while the APT indicates positive interaction driven by a shared region of high cell density. The TCM helps reconcile these findings: local clustering is offset by areas rich in ‘UD structural SM’ cells but lacking ‘myofibroblasts’, resulting in a neutral global PCF. Together, these analyses show how different statistics capture distinct aspects of spatial organisation.

**Fig. A3:**
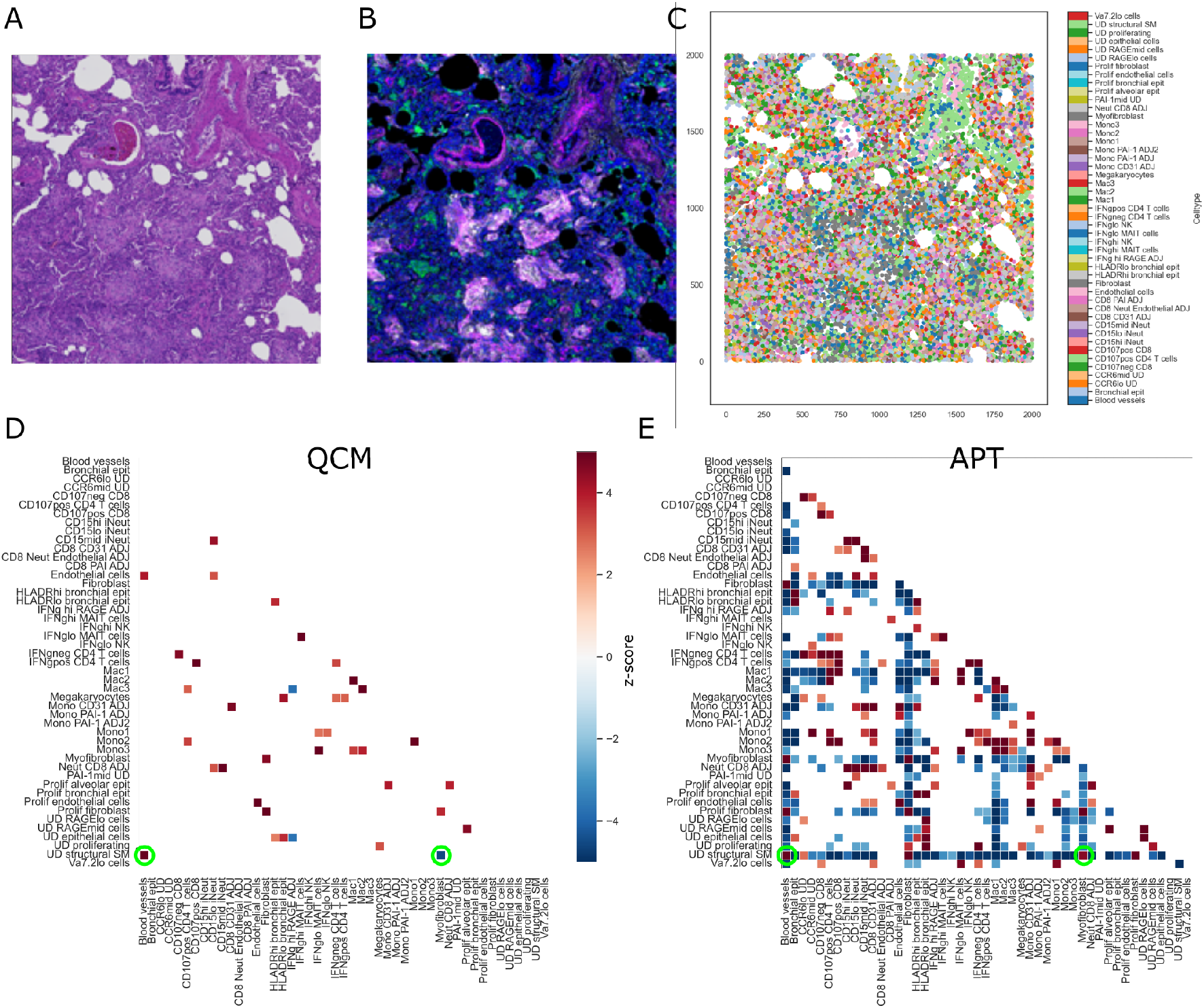
MuSpAn applied to Imaging Mass Cytometry (IMC) data. A: H&E staining of a 2mm *×* 2mm ROI from a human lung resected from a COVID-19 infected patient [S2]. B: False colour IMC staining of the same lung ROI. C: Point cloud of cell centres from the ROI, with annotations from [S2]. D: Quadrat correlation matrix (QCM) for the cells in this ROI (square quadrats, edge length 100µm), showing statistically significant values only and highlighting the cell pairs analysed further. E: Adjacency permutation test (APT) for the cells in this ROI (Delaunay network, maximum edge length 20µm), showing statistically significant values only and highlighting the cell pairs analysed further.

**Fig. A4:**
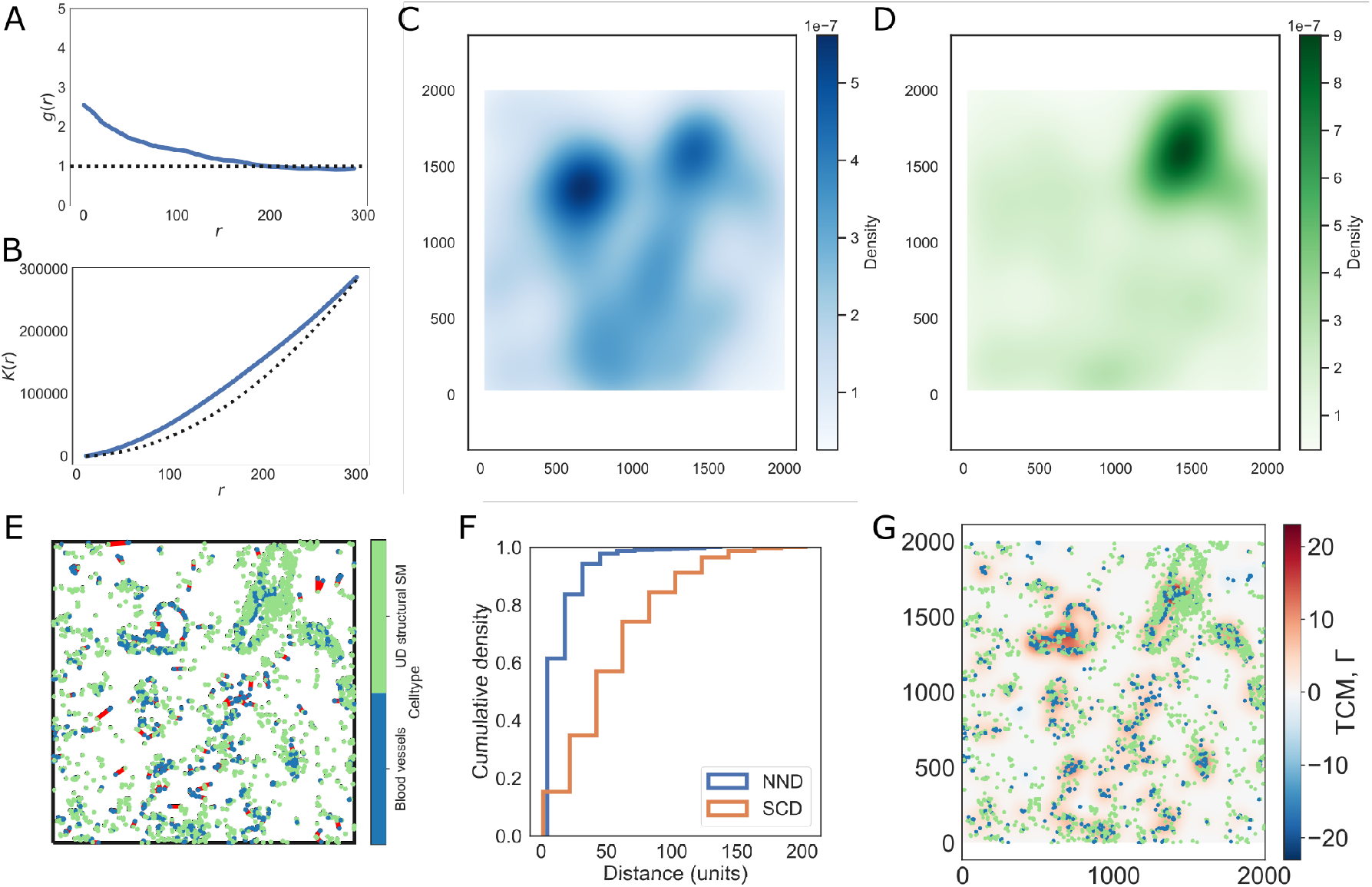
MuSpAn applied to IMC data - positive spatial correlation. A: Cross-PCF function between blood vessels and ‘UD structural SM’ cells, indicating clusters of the two populations with a characteristic size of approximately 180µm. B: Cross-*K* function between the two cell populations, indicating increased association at length scales up to 300µm. C: Kernel Density Estimation (KDE) of blood vessels in the ROI. D: KDE of ‘UD structural SM’ cells in the ROI. E: Nearest neighbour distances (red) from blood vessels to the nearest ‘UD structural SM’ cell, two populations which are highlighted as spatially correlated by the QCM in Fig. A3D. F: Cumulative density function of the nearest neighbour distribution (NND), compared with a null of the spherical contact distribution for blood vessels (SCD). G: Topographical Correlation Map (TCM) showing where blood vessels are locally positively or negatively associated with ‘UD structural SM’ cells.

**Fig. A5:**
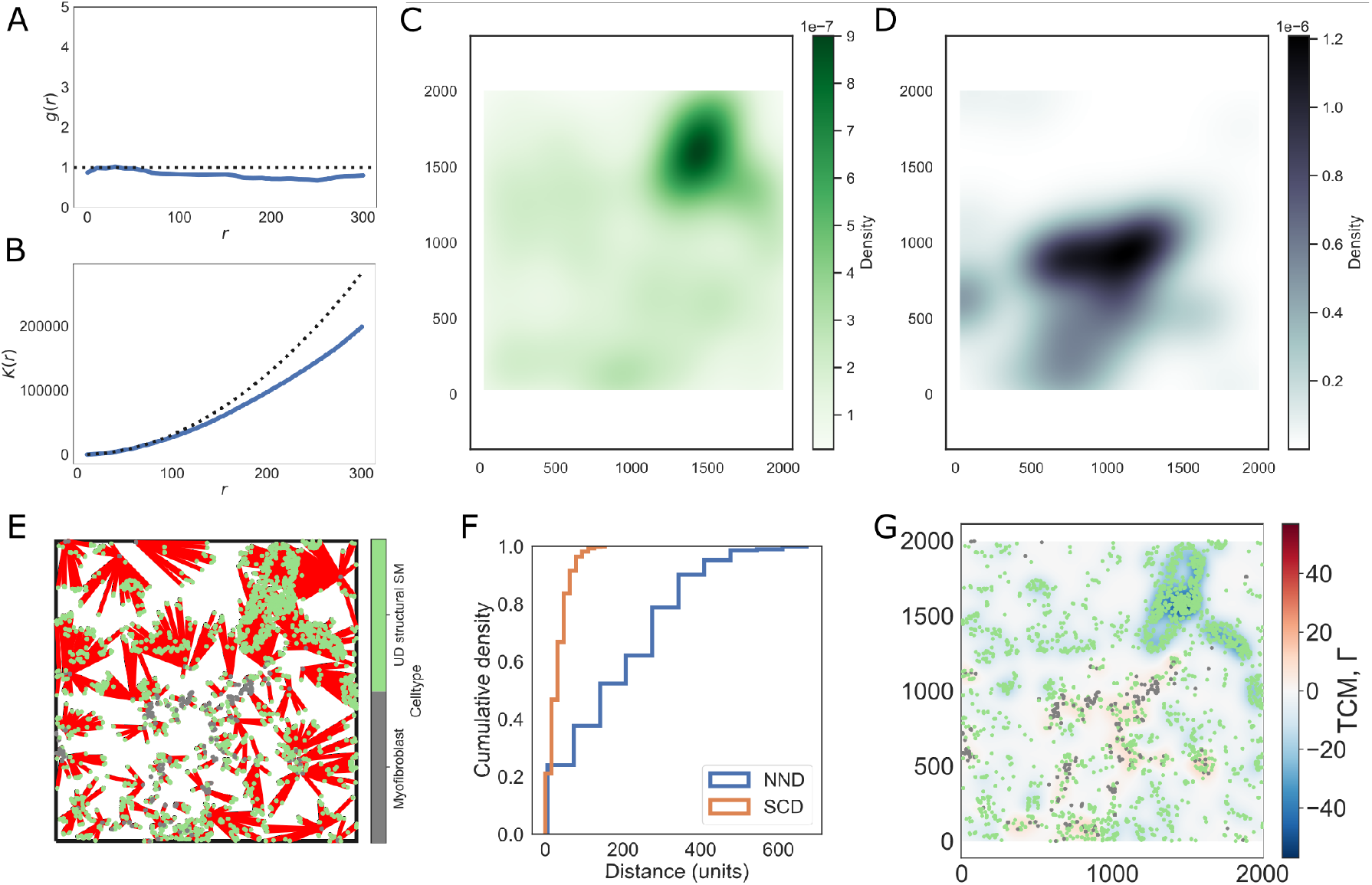
MuSpAn applied to IMC data - spatial anti-correlation. A: Cross-PCF between ‘UD structural SM’ cells and myofibroblasts, two populations which are highlighted as spatially anti-correlated by the QCM in Fig. A3D. This indicates no strong positive or negative correlation between the populations at short length scales, and weak anti-correlation at length scales beyond 100µm. B: Cross-*K* function between the two cell populations, indicating decreased association at length scales up to 300µm and beyond. C: Kernel Density Estimation (KDE) of ‘UD structural SM’ cells in the ROI. D: KDE of myofibroblasts in the ROI. E: Nearest neighbour distances (red) from ‘UD structural SM’ cells to myofibroblasts. F: Cumulative density function of the nearest neighbour distribution (NND), compared with a null of the spherical contact distribution for ‘UD structural SM’ cells (SCD). G: Topographical Correlation Map (TCM) showing where ‘UD structural SM’ cells are locally positively or negatively associated with myofibroblasts.

**Fig. A6:**
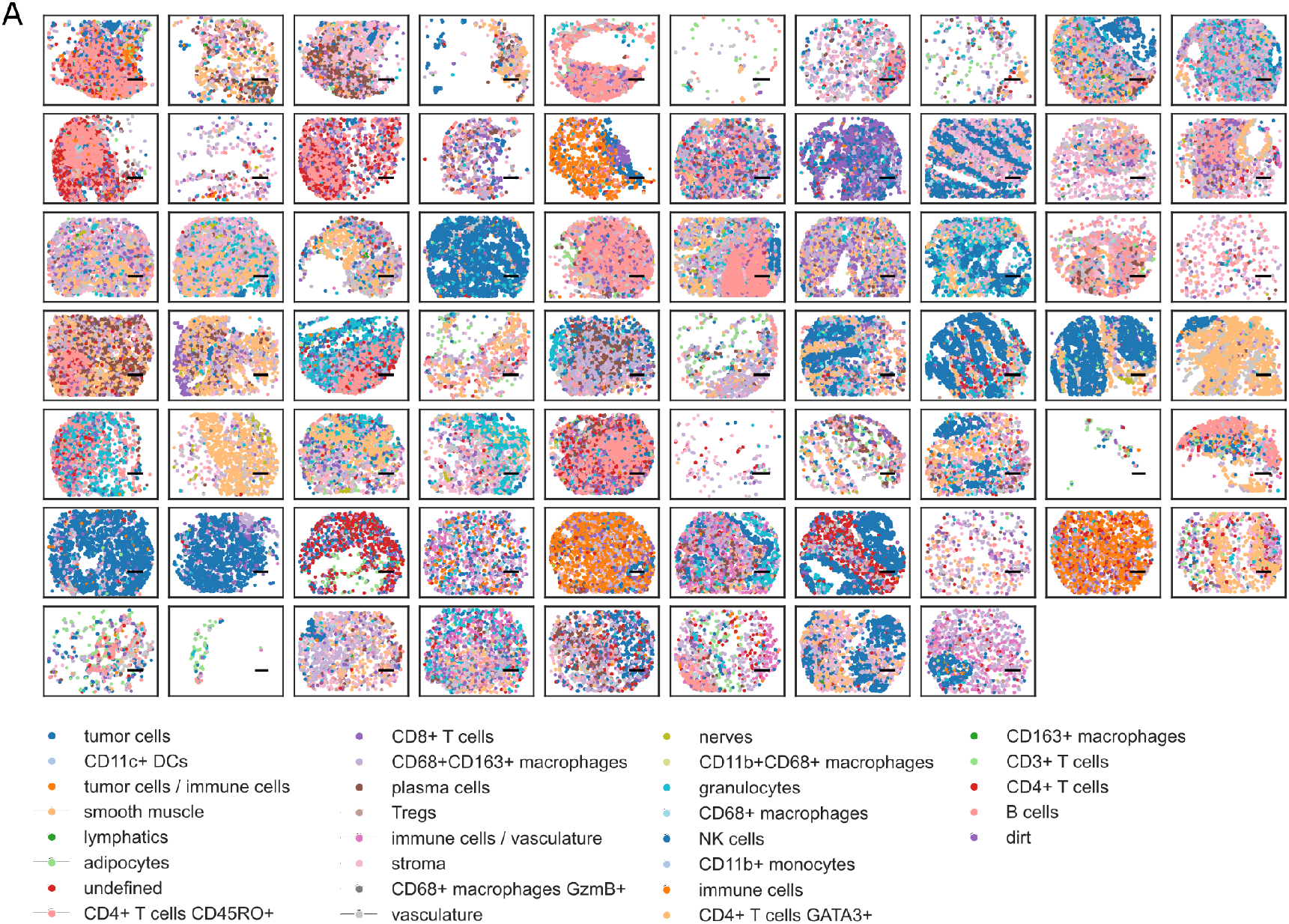
MuSpAn applied to CODEX data. A: MuSpAn visualisation of cell centres from the TMAs used in [S3].

### A.3 CODEX - Schürch *et al*. (2020)

We use MuSpAn to reproduce and extend the analysis of CODEX tissue microarray (TMA) data in [S3]. Fig. A6A shows a selection of 68 TMAs (‘group 1’, regions with “Crohn’s like reaction”, CLR) from this dataset. We use MuSpAn to conduct a neighbourhood enrichment study based on a *k*-nearest neighbours network (and assigned 10 *k*-means clusters using `muspan.networks.cluster_neighbourhoods()` to obtain an enrichment matrix for each cluster, Fig. A7A). The clusters obtained from this analysis are in good qualitative agreement with those obtained by Schürch *et al*. [S3]. MuSpAn assigned clusters are shown in Fig. A7B and D, and the corresponding cluster assignments for the same regions from [S3] are shown in Fig. A7C and E (note that since neighbourhood cluster assignment is conducted on all samples simultaneously, not every cluster is represented in every TMA).

We extend this analysis by merging adjacent Voronoi cells in each TMA which share the same label to form contiguous regions within which all cells belong to the same cellular neighbourhood. This facilitates applying MuSpAn operations directly to the cellular neighbourhoods, treating each contiguous region as a separate shape. We conduct an adjacency permutation test (APT) on each TMA, obtaining a standardised effect size (SES, *z*-score) describing whether any two types of neighbourhood are found in direct contact more or less frequently than would be expected under random labelling. For a given pair of neighbourhoods, we compare the distribution of SES (*z*-score) in ‘group 1’ (Crohn’s like reaction, CLR) and ‘group 2’ (diffuse inflammatory infiltration, DII) TMAs. Fig. A8A shows the SES for interactions between Neighbourhood 4 and Neighbourhood 8 in each TMA, which are significantly higher in the DII regions than the CLR (*p* ≈ 0.0013, unequal variance *t*-test). Similar tests applied to all combinations of Neighbourhoods identify a small number of statistically significant differences in the neighbourhood connectivity between CLR and DII (Fig. A8B).

**Fig. A7:**
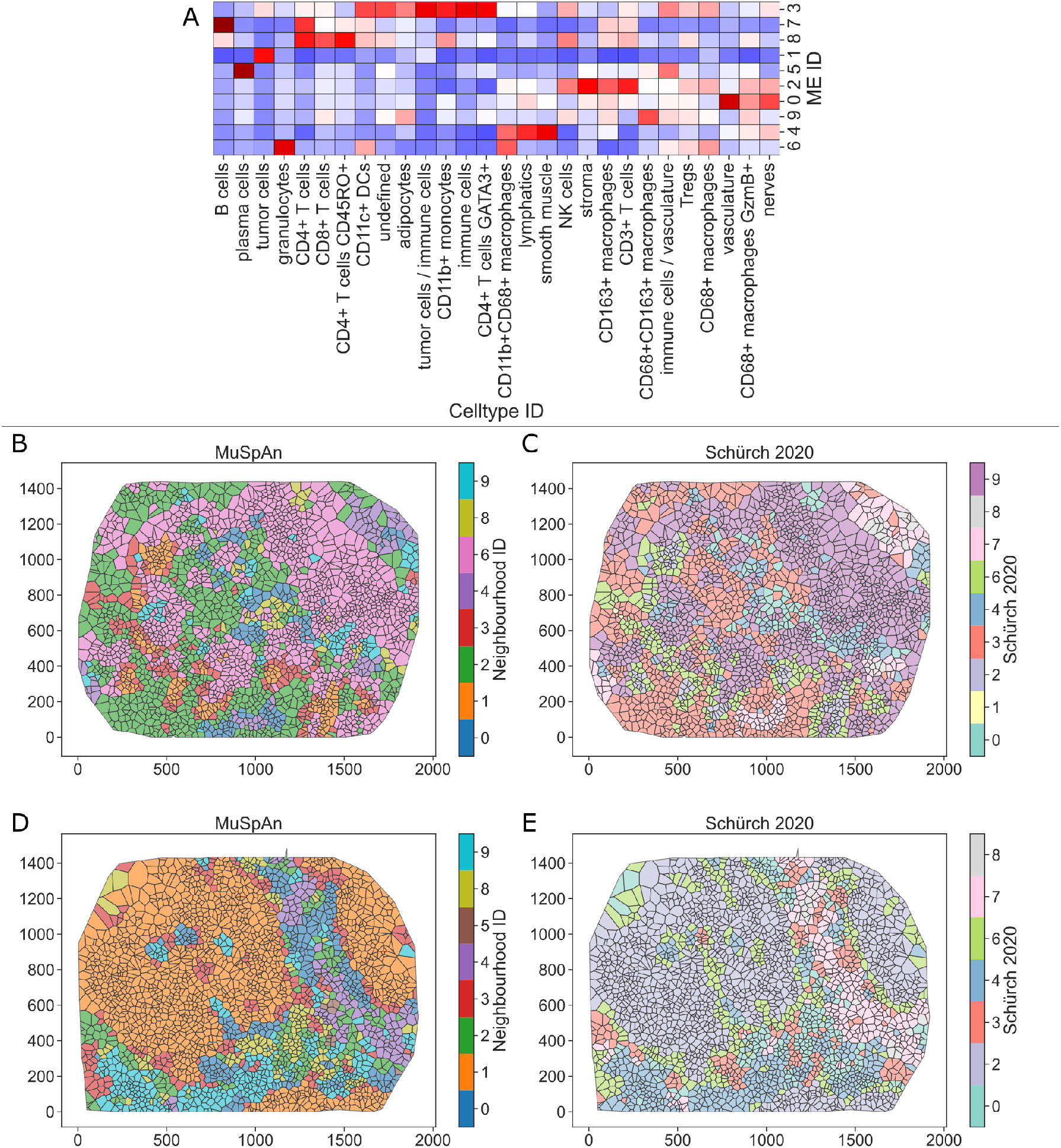
MuSpAn applied to CODEX data. A: A microenvironment enrichment matrix showing the celltypes which are enriched in each of the 10 microenvironment clusters (ME ID 0-9) detected in a MuSpAn reproduction of the Schürch *et al*. pipeline. B-E: Voronoi representations of cell data from two TMAs (B/D and C/E), coloured according to the MuSpAn assigned cellular neighbourhoods (B, D) and to the Schürch *et al*. assigned neighbourhoods (C, E).

**Fig. A8:**
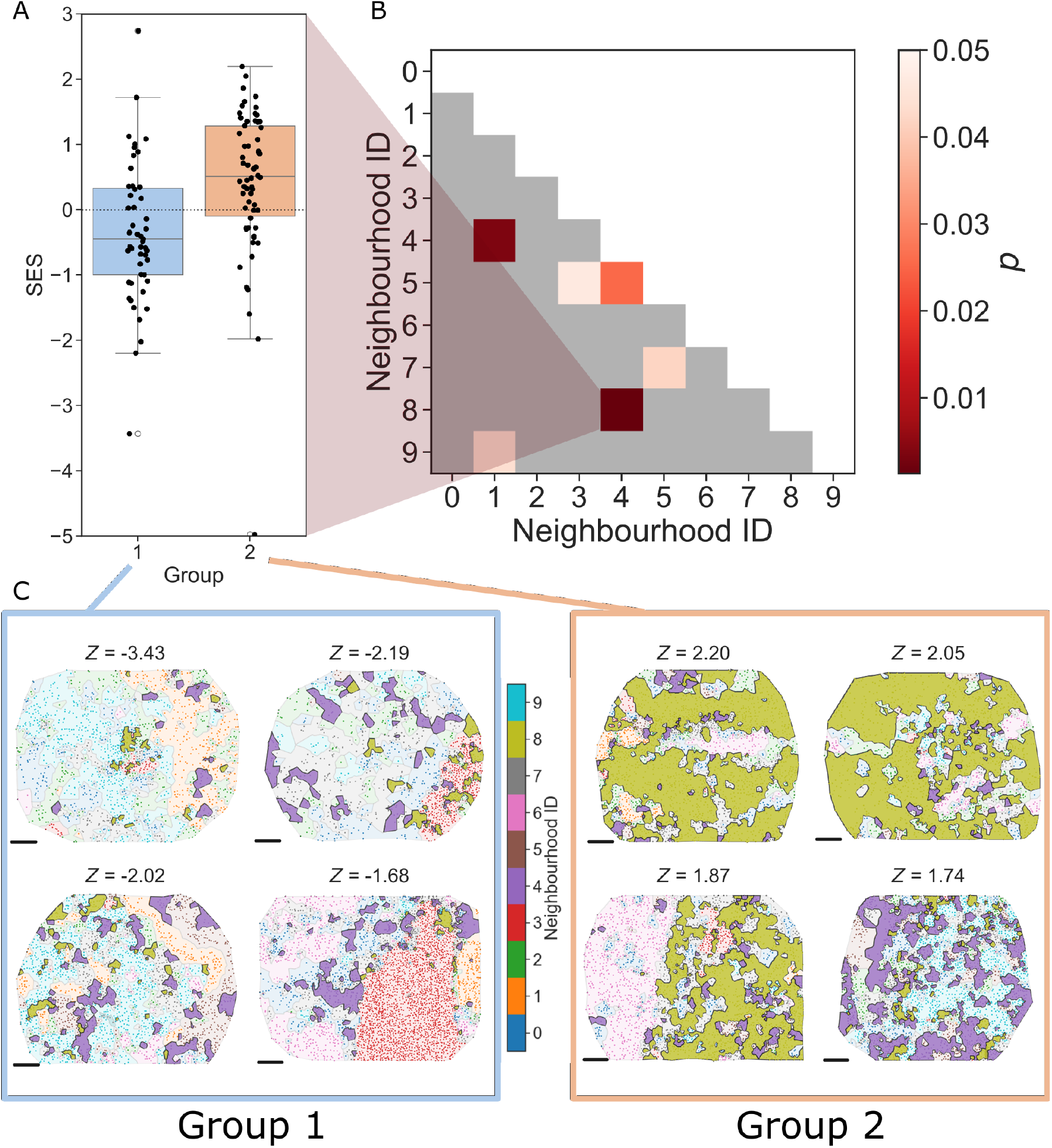
Neighbourhood contacts vary between CLR and DII TMAs. A: Standardised effect sizes (*z*-scores) for adjacency permutation test (APT) interactions between Neighbourhoods 4 and 8, within each TMA from group 1 (CLR) and group 2 (DII) (*p* = 0.0013, unequal variance *t*-test). B: *p*-values showing which neighbourhood-neighbourhood interactions have significantly different APT scores betwen group 1 and group 2 (unequal variance *t*-tests). C: The four TMAs with the lowest SES (group 1) or highest SES (group 2). Regions corresponding to neighbourhoods 4 (purple) and 8 (olive) are highlighted. In group 1, these neighbourhoods are small and rarely in direct contact. In group 2, the neighbourhoods are much larger and frequently share boundaries. Scalebars are 200 pixels.

To better interpret these scores, we show the four TMAs from group 1 (CLR) with the lowest *z*-scores, and those from group 2 (DII) with the highest (Fig. A8C). TMAs with low *z*-scores (CLR) generally have small neighbourhoods of types 4 and 8, which are rarely in direct contact. In contrast, the type 4 and 8 neighbourhoods in group 2 (DII) are much larger, and are found frequently in direct contact with one another. These neighbourhoods approximately correspond to the ‘T cell enriched’ (neighbourhood 8) and ‘smooth muscle’ (neighbourhood 4) cellular neighbourhoods described in [S3], suggesting that there is a significant difference between the localisation of T cell infiltration into stromal regions between CLR and DII regions.

## Appendix B Biologically-driven pipeline generation with MuSpAn

The examples in the main text (Fig. 6) and Appendix A illustrate how MuSpAn enables the construction of spatial analysis pipelines tailored to specific biological hypotheses. Each pipeline applies distinct statistical methods to interrogate different aspects of spatial organisation. Crucially, spatial analysis is not a one-size-fits-all process - methods suitable for investigating one hypothesis may be inappropriate for another.

Effective pipeline design hinges on selecting statistics that capture the biological feature of interest from complementary perspectives. For clarity, we demonstrate how different mathematical approaches highlight distinct spatial features with reference to a simple synthetic dataset in which blue and orange cells form a ring, surrounded by randomly placed green points (Fig. B9). We focus on three commonly-arising spatial features of a dataset: cell-cell contact (Fig. B9A), description of local cellular neighbourhoods (Fig. B9B), and characterisation of emergent tissue architectural features (Fig. B9C). We emphasise that these examples represent just a subset of the perspectives that could analysed with MuSpAn, which is designed to make bespoke pipeline development simple. Our aim here is not to prescribe a fixed set of analyses, but to show how MuSpAn provides a flexible framework adaptable to diverse biological contexts.

### Dataset

The synthetic dataset used in Fig. B9 is available within MuSpAn as the *‘Synthetic-Points-Architecture’* dataset, and can be easily loaded via the command `muspan.datasets.load_example_domain(` *‘Synthetic-Points-Architecture’*). The panels shown represent a subset of the full dataset centred around a ring of orange and blue points, with green points randomly placed throughout the domain; for full details on the dataset, see Figure 1 of [S6]; code for generating the panels shown is available alongside this paper.

#### B.1 Cell-cell contact

Fig. B9A illustrates three complementary approaches for analysing cell-cell contact:

- **Adjacency networks** provide a global view of direct cell-cell interaction that considers all cells simultaneously. They can be constructed from segmentation masks, or inferred from cell centroids using a Delaunay triangulation. Network edges may be weighted by metrics such as intercellular distance, and statistical tests like adjacency permutation [S7] can assess contact patterns across multiple cell types simultaneously.
- **Nearest neighbour analysis** instead focusses on pairwise spatial relationships between specific cell types. For example, measuring the distance from each green cell to the nearest blue cell yields a distribution that can be tested for spatial association using metrics like the Average Nearest Neighbour Index [S8]. This approach is particularly useful when it is hypothesised that direct contact is not required, but proximity is biologically meaningful.
- **Local indicators of spatial association (LISA)** offer a spatially resolved view of co-occurrence. The topographical correlation map (TCM) [S9], for instance, highlights regions where green cells are locally enriched (red) or depleted (blue) relative to the blue cell population. MuSpAn supports several LISA variants, including the Local Getis-Ord statistic (Fig. 5I).

**Fig. B9:**
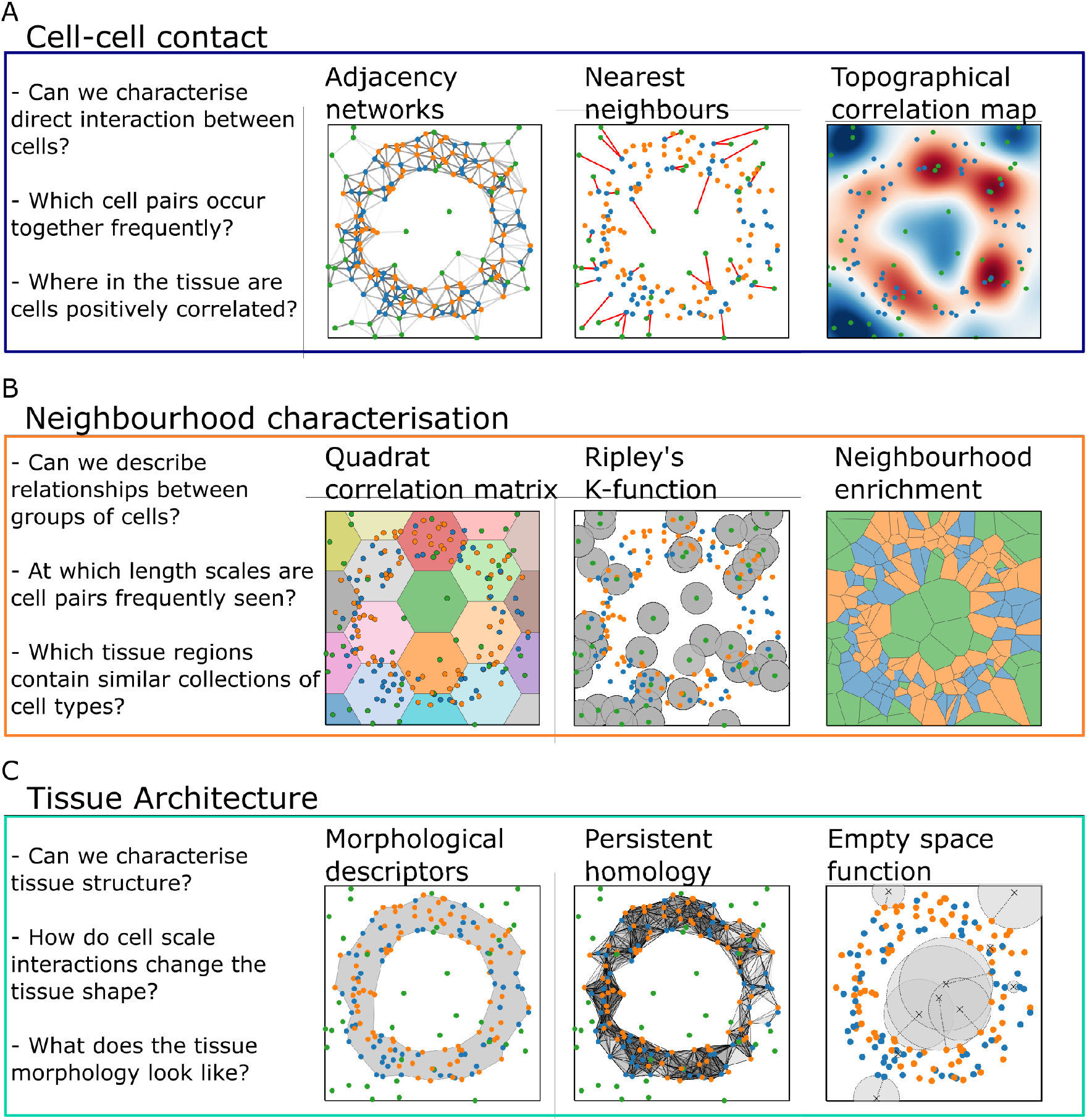
Biologically-driven pipeline development using MuSpAn. Examples of how complementary mathematical methods can be combined to characterise different biological features. In each example we use the same illustrative points, and schematise how the methods considered use different aspects of the data. This synthetic dataset contains three cell types: blue and orange cells form a circle, and green cells are randomly distributed. A: Cell-cell contact could be described using adjacency networks, nearest neighbour approaches, and topographical correlation maps. B: Neighbourhood characterisation can be conducted using region-based approaches such as the QCM, spatial statistics like Ripley’s K function, and neighbourhood enrichment scores. C: Tissue architecture can be described using standard morphological descriptors (such as area/perimeter, circularity, convexity), persistent homology, and the empty-space function.

#### B.2 Neighbourhood characterisation

Fig. B9B presents three strategies for describing local cellular neighbourhoods:

- **Quadrat based methods** divide the ROI into fixed-size tiles, aggregating spatial data within each. Metrics such as the quadrat correlation matrix (QCM) and Morisita-Horn index quantify label diversity and similarity across tiles [S10, S11], offering a coarse-grained view of spatial heterogeneity.
- **Spatial statistics** can examine the local environment around individual cells. For example, Ripley’s cross-*K* function measures the spatial co-occurrence of two cell types within a defined radius [S12]. This approach captures average neighbourhood properties and can be extended to analyse interactions among three or more cell types [S9].
- **Region-based enrichment methods** define neighbourhoods based on locally dominant cell types. In the schematic, the domain is partitioned using Voronoi tessellation; the cellular composition within each region can then be characterised using, for example, *k*-nearest neighbour networks [S3]. MuSpAn includes several alternative methods for identifying and analysing spatially enriched zones.

#### B.3 Tissue architecture

To analyse multicellular structures at larger scales, Fig. B9C shows three complementary approaches:

- **Morphological descriptors** quantify the shape of defined structures. Using *α*-shapes [S13], cell locations can be converted into explicit boundaries, enabling measurement of features such as area, circularity, and convexity (see Fig. 2).
- **Persistent homology (PH)** captures topological features like loops or connected components of point datasets when connected at different length scales, without requiring predefined boundaries. In the schematic, a Vietoris-Rips complex constructed from blue and orange cell positions identifies the ring structure. PH quantifies the persistence of such structures across spatial scales, offering a robust method for detecting architectural motifs.
- **Spatial statistics** including the empty space function (spherical contact distribution) provide a statistical view of voids within the ROI [S1]. Unlike nearest neighbour functions, which measure distances between cells, these functions assess the distance from random points to the nearest cell, characterising the size and distribution of empty regions.

## Appendix C Available quantitative tools

Table C1 contains a list of the data analysis methods available within MuSpAn v1.2.0; since the module is under continual development, subsequent versions contain a wider array of features. For further details about how each method works, we refer the interested reader to the citations in Table C1, or to the documentation available at https://docs.muspan.co.uk/ where quantitative tools available in later versions can be found.

**Table C1:** A summary of the data analysis methods available within each module of MuSpAn v1.2.0, together with a brief summary of their purpose. For further details about these methods, please see the references provided.

| Method name<br>(MuSpAn Module) | Summary | References |
| --- | --- | --- |
| Label Count<br>(summary_Statistics) | Number of objects with a given label, within a given region . | - |
| Label Density<br>(summary_Statistics) | Count, divided by region area. | [S2, S14] |
| Label Entropy<br>(summary_Statistics) | Shannon entropy of a specified label. | [S11, S15] |
| Generate Distribution<br>(distribution) | Convert point data into a distribution using general kernel. | - |
| Kernel Density Estimation<br>(distribution) | Convert point data into a probability density distribution using a Gaussian kernel function. | [S16] |
| Kullback-Leibler Divergence<br>(distribution) | Measures the similarity between two distributions using relative entropy. | [S17] |
| Wasserstein distance<br>(distribution) | Measures the distance between two distributions. | [S6, S18] |
| Summarise Distribution<br>(distribution) | Computes the mean and standard deviational ellipses of spatial point data. | - |
| Area<br>(geometry) | Area of a shape. | - |
| Circularity<br>(geometry) | Circularity of a shape. | - |
| Convexity<br>(geometry) | Convexity of a shape (ratio of the area to the area of its convex hull) | - |
| Perimeter<br>(geometry) | Perimeter of a shape. | - |
| Principle Axis<br>(geometry) | Computes the angle and major axis of a polygon relative to the x-axis. | - |
| Adjacency Permutation Test<br>(networks) | Test the correlation between adjacent labels in a network by randomly permuting them (also referred to as ‘neighbourhood enrichment’). | [S5, S7] |
| Alpha Index<br>(networks) | A measure of connectivity which evaluates the number of cycles in a graph in comparison with the maximum number of cycles. ($\approx 0$ for a Tree, $\approx 1$ for a maximal planar graph). | [S19, S20] |
| Betweenness Centrality<br>(networks) | The sum of the fraction of all shortest paths that pass through each node. | [S20, S21] |
| Closeness Centrality<br>(networks) | The average shortest distance from each node to all other reachable nodes. | [S20, S21] |
| Cluster Neighbourhoods<br>(networks) | Cluster similar neighbourhoods based on the enrichment of labels within the neighbourhoods. | [S3, S22] |
| Community Detection<br>(networks) | Perform community detection based on node connectivity. | [S23] |
| Compactness<br>(networks) | Index in the range $[0, 1]$ that relates the area of a network to the length of its edges. | [S20, S21] |
| Cyclomatic Number<br>(networks) | The number of elementary cycles within a graph. | [S20] |
| Edge Orientation Distribution<br>(networks) | Compute the (symmetric) distribution of edge orientations. | [S20] |
| Gamma Index<br>(networks) | A measure of connectivity that considers the relationship between the number of observed links and the number of possible links. | [S19, S20] |
| Network Generation<br>(networks) | Generate a network based on MuSpAn objects. Available methods include Delaunay, $k$-Nearest Neighbours, and Proximity networks. | [S3, S20, S24–S26] |
| Katz Centrality<br>(networks) | The influence of a node in a network, determined by considering the total number of walks, weighted by their lengths, that connect the node to all other nodes.. | [S20] |
| $k$-Hop Neighbourhood<br>(Networks) | Find the $k$-hop neighbourhood of a set of source objects | - |
| $k$-Nearest Neighbour<br>Label Count<br>(networks) | Count the occurrences of each label within $k$ steps from a set of source objects | - |
| Laplacian Centrality<br>(networks) | A measure of node connectivity in a graph | - |
| Network Distance<br>(networks) | Measure the similarity of two networks, using a range of available distance metrics. | [S27] |
| Node Degree<br>(networks) | Compute the degree of each node. | - |
| Generate Quadrats<br>(region-based) | Create a mesh of quadrats covering the requested space. | - |
| Generate Hexgrid<br>(region-based) | Create a mesh of regular hexagons covering the requested space. | - |
| Morisita Horn Index<br>(region-based) | A measure of the similarity of observations of a label within a set of regions. | [S10, S11] |
| Quadrat Correlation Matrix<br>(region-based) | Describe correlations between the counts of different cell types within a set of regions. | [S2, S28] |
| Average Nearest Neighbour Index<br>(spatial_statistics) | A measure of spatial aggregation based on the average distance between pairs of objects. | [S8, S15] |
| Cross Ripley's K Function<br>(spatial_statistics) | A measure of (cumulative) spatial aggregation between pairs of objects across length scales. | [S29] |
| Cross L Function<br>(spatial_statistics) | A normalised version of Ripley's K-Function. | [S29] |
| Cross Pair Correlation Function<br>(spatial_statistics) | A measure of spatial aggregation between pairs of objects across length scales. | [S9, S29] |
| Getis Ord Statistic<br>(spatial_statistics) | A measure of spatial autocorrelation that identifies hot- and cold-spots in the distribution of a continuous label. | [S30–S32] |
| J-Function<br>(spatial_statistics) | A test that identifies clustering and dispersion in point patterns. Relates the Spherical Contact Distribution and the Nearest Neighbour Distribution. | [S1, S33] |
| Moran's I<br>(spatial_statistics) | Calculate both local and global Moran's I statistic within the domain. The local Moran's I statistic quantifies the spatial autocorrelation for each object in the population. The global Moran's I statistic quantifies the spatial autocorrelation for the entire population. | [S15, S26, S31] |
| Nearest Neighbour Distribution<br>(spatial_statistics) | Calculate the distribution of distances between the closest objects in two populations. | [S22] |
| Spherical Contact Distribution<br>(spatial_statistics) | Calculate the distance from randomly sampled points to the closest objects in a population. Also referred to as the 'Empty Space Distribution'. | [S1] |
| Topographical Correlation Map<br>(`spatial_statistics`) | A spatially-resolved, quantitative summary of colocalisation between two point populations across a region of interest. | [S2, S9] |
| Weighted Pair Correlation Function<br>(`spatial_statistics`) | An extension of the cross PCF which accounts for continuous labels. | [S9, S34] |
| Level Set Filtration<br>(`topology`) | Calculate the level set filtration of a distribution or image. | [S6, S35] |
| Vietoris-Rips Filtration<br>(`topology`) | Calculate the Vietoris-Rips filtration of a set of points. | [S35] |
| Vectorise Persistence<br>(`topology`) | Convert persistence diagram into a vector for subsequent analysis. | [S36] |
| Cluster Data<br>(`helpers`) | Wrapper for clustering methods implemented in scikit-learn. | [S37] |
| Crop Domain<br>(`helpers`) | Return a new domain (or domains) based on a set of new boundary shapes (or label values). | - |
| Combine Shapes<br>(`shape_operations`) | Combine two shapes in the domain using a specified operation. | - |
| Return Difference of Shapes<br>(`shape_operations`) | Compute the difference between two shapes. | - |
| Return Intersection of Shapes<br>(`shape_operations`) | Compute the intersection of two shapes. | - |
| Return Union of Shapes<br>(`shape_operations`) | Compute the union of two shapes. | - |

## Appendix D Further information on spatial analysis methods

Below, we describe the spatial analysis methods used to generate Figs. 2-5 of the main text. The methods are organised to mirror the subsections of the Results section in the main text.

### D.1 Spatial analysis of individual cells

#### D.1.1 Circularity

**Description:** Circularity measures how closely a shape resembles a circle.

**Definition:** Let *O* be a polygon with area, *A*, and perimeter, *P*. Then, the circularity, *C*, of *O* is defined as

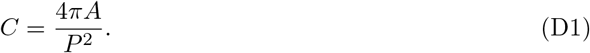

**Application:** In Fig 2F, circularity is computed for all ‘Cell boundary’ objects labelled as SMC or PROG.

#### D.1.2 Principal angle

**Description:** A 2D shape has two principal axes: the principal major axis is the axis of maximum moment of area (maximising bend resistance), and the principal minor axis is the axis of minimum moment of area (minimum bend resistance) [S38].

**Definition:** The principal angle is the angle between the principal major axis and the line *y* = 0. Let *O* be a polygon defined by *n* vertices ***v***_*i*_ = (*x*_*i*_, *y*_*i*_) ∈ ℝ^2^ (1 ≤ *i* ≤ *n*), ordered anti-clockwise.

The second moments of area, *I*_*x*_, *I*_*y*_ and *I*_*xy*_, are given by

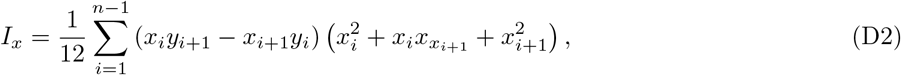

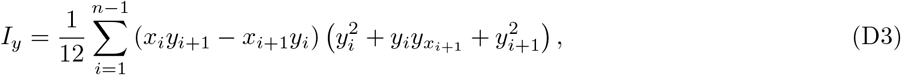

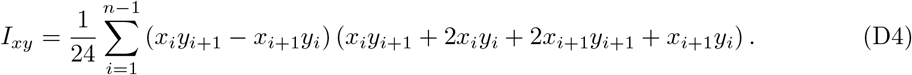

They describe how the vertices of a polygon are distributed with respect the x and y-axis. The principal angle, *θ*_*O*_, of *O* is defined by

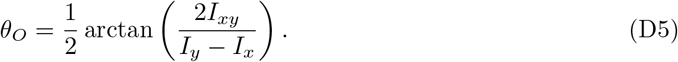

**Application**: In Fig 2G, the principal angle is computed for all ‘Cell boundary’ objects with labels SMC or PROG.

### D.2 Cell proximity analysis

#### D.2.1 Nearest neighbour metrics

**Description:** The Average Nearest Neighbour Distance is the mean minimum distance from an object in one population to an object in another. It is closely related to the Average Nearest Neighbour Index (ANNI), which measures the degree of clustering (or dispersion) of a set of points by comparing the average nearest neighbour distance to the expected nearest neighbour distance in a random distribution of the same density of points, here assumed to be complete spatial randomness (CSR) [S8].

**Definition:**Let Ω be a spatial domain with area *α*. Let *d*_*A,B*_ denote the minimum Euclidean distance between objects *A* and *B*, for *A, B* contained in Ω. Let 𝒪_*A*_ = {*A*_1_, …, *A*_*n*_} and 𝒪_*B*_ = {*B*_1_, …, *B*_*m*_} be two sets with objects *A*_*i*_ (*i* = 1, …, *n*) and *B*_*j*_ (*j* = 1, …, *m*) in Ω. Then the minimum nearest neighbour distance from a specific object *A*_*k*_ to an object in the set 𝒪_*B*_ is

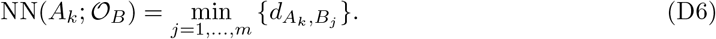

The average nearest neighbour distance for objects from the set 𝒪_*A*_ to those in 𝒪_*B*_ is then

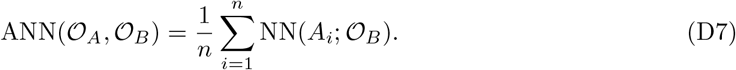

Assuming CSR, the expected average nearest neighbour distance, 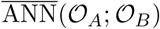, is given by

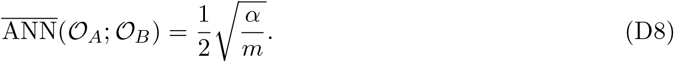

The Average Nearest Neighbour Index, ANNI(𝒪 _*A*_; 𝒪_*B*_) is expressed as the ratio of the observed to the expected average nearest neighbour distance:

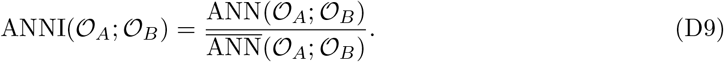

The significance of the difference between the observed nearest neighbour distribution and that expected under CSR is typically quantified using the *z*-score (standard normal variate), *z*, such that

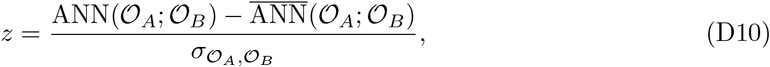

where the variance, 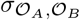, is evaluated as

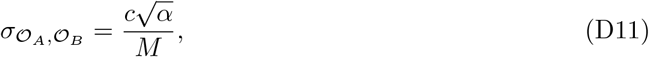

with *c* = 0.26136 [S8].

**Application**: The ANNI was used to investigate the spatial relationships between SMC and STR1 cells, and PROG and COL cells in Figs 3B-3E.

#### D.2.2 Quadrat correlation matrix

**Description:** The quadrat correlation matrix (QCM) describes correlations between counts of two labels of interest (here, Celltype IDs) within a set of ‘quadrats’, a regular tiling of the domain of interest with non-overlapping areas [S2, S6, S28, S39]. For a given pair of labels, the QCM quantifies how different the observed correlation coefficient across the quadrats is from that expected under a random shuffling of the object labels.

**Definition:** The QCM is calculated by comparing the observed matrix of cell counts within each quadrat, **O**, against a null distribution of *K* null matrices, **N**^k^ for *k* ∈ [1, …, *K*], whose row and column sums are the same as those of **O**. The standard effect size (SES / *z*-score) for Celltypes *i* and *j* is given by

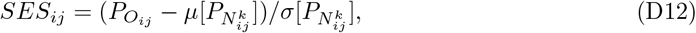

where *P*_*A*_ is the partial correlation matrix associated with matrix A, and *µ* and *σ* are the mean and standard deviation over all *k*. The entry of the QCM corresponding to Celltypes *i* and *j* is *SES*_*ij*_ if the *p*-value associated with *SES*_*ij*_ score is statistically significant after applying the Benjamini-Hochberg correction for multiple comparisons, and 0 otherwise.

**Application:**In Fig. 3J, the QCM was computed using a square lattice of edge length 75µm and shuffled over 1000 iterations (*K* = 1000). For simplicity, values shown in Fig. 3J are not filtered for statistical significance (so, strictly speaking, this is the SES matrix, rather than the QCM).

#### D.2.3 Topographical Correlation Map (TCM)

**Description:** The Topographical Correlation Map (TCM) is a spatial statistic derived from the general class of local indicators of spatial association (LISA) [S40], designed to identify the spatially resolved strength of positive or negative correlation between two cell populations [S2, S9]. Unlike the cross-PCF (see below), which provides a global summary, the TCM assigns a continuous-valued mark to each individual cell and smooths these to produce a spatial map of local associations.

**Definition:** Let *C*_*A*_ and *C*_*B*_ be two cell populations, and let ***x***_*i*_ denote the spatial location of a cell of type *C*_*A*_. To compute the TCM, we first define a mark *m*_*AB*_(*r*, ***x***_*i*_) for each such cell, which quantifies its cumulative contribution to the cross-PCF up to radius *r*:

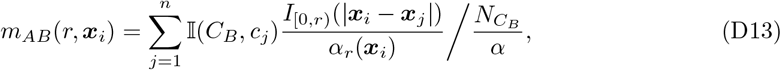

where *α* is the total area of the domain, 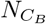 is the number of cells of type *C*_*B*_, *α*_*r*_(***x***_*i*_) is the area of intersection between an annulus of radius *r* centred at ***x***_*i*_ and the domain, and 𝕀 (*C*_*B*_, *c*_*j*_) is an indicator function equal to 1 if *c*_*j*_ ∈ *C*_*B*_, and 0 otherwise. This mark is conceptually similar to the cross-PCF: values *m*_*AB*_ *>* 1 indicate local clustering of *C*_*B*_ around ***x***_*i*_, while values *m*_*AB*_ *<* 1 indicate local exclusion.

To facilitate comparison and aggregation across cells, *m*_*AB*_ is normalised to a bounded, symmetric score *µ*_*AB*_(*r*, ***x***_*i*_), using a tuning parameter *γ*:

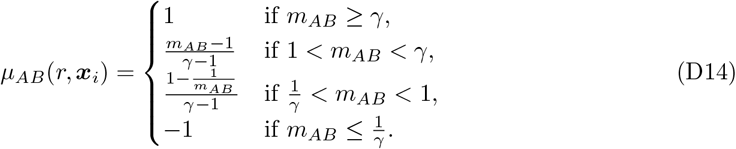

This ensures that *µ*_*AB*_ ∈ [−1, 1], where values near 1 and −1 represent the strongest possible clustering and exclusion respectively (bounded by *γ*), and values near 0 indicate no association. The scaling is symmetric, so *µ*_*AB*_ = 0.5 and *µ*_*AB*_ = −0.5 represent equal strengths of clustering and exclusion, respectively.

To generate a spatial map over the domain, each *µ*_*AB*_(***x***_*i*_) is convolved with a Gaussian kernel of standard deviation *σ*, centred at ***x***_*i*_. The resulting TCM, Γ_*AB*_(*r*, ***x***), is defined as:

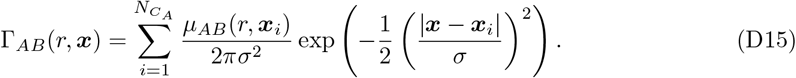

Note that the TCM is not symmetric in its arguments; that is, Γ_*AB*_(*r*, ***x***) / ≠ Γ_*BA*_(*r*, ***x***), since the kernel locations are determined by the *C*_*A*_ population.

**Application:** The TCM was applied in the multiscale spatial profiling of immune–fibroblast interactions in CMS4-like colorectal tumours to quantify the spatial heterogeneity of cell–cell associations within regions of interest (ROIs). In this context, we examine selected fibroblast–immune cell type pairs using *r* = 20µm and *σ* = 25, to highlight localised clustering or exclusion patterns across the tissue microenvironment.

### D.3 Analysis of cell-cell relationships across scales

#### D.3.1 Cross pair correlation function

**Description:** The cross pair correlation function (cross-PCF) is a spatial statistic that relates the observed number of point pairs from populations *A* and *B* that are separated by distance *r* against the expected number under a statistical null distribution.

**Definition:** Let **x**^*A*^ and **x**^*B*^ be the sets of locations of points in populations *A* and *B* respectively, and suppose there are *N*_*A*_ and *N*_*B*_ points in each set. Consider the indicator function

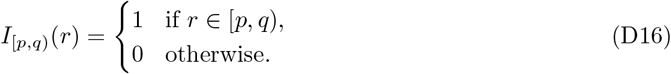

The cross-PCF, *g*_*AB*_(*r*), is given by

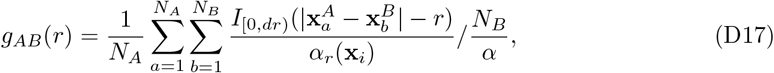

where *α* is the area of the domain of interest, and *α*_*r*_(**x**) is the area of the intersection between the domain and an annulus centred at **x** with inner radius *r* and outer radius *dr*. This provides a boundary correction term for points close to the domain edge; alternative methods for boundary correction can be found in [S29].

**Application:** In Fig. 4B we use *dr* = 15µm and calculate *g*_*AB*_(*r*) at discrete values *r*_*k*_ ∈ [0, 5, 10, …, 250], excluding any points within 20µm of the region boundary from the calculation.

#### D.3.2 Vietoris-Rips filtration

**Description:** A Vietoris–Rips (VR) filtration is a method from persistent homology that can be used to quantify the topological features (e.g. connected components, loops, voids) of point clouds at multiple spatial scales [S35].

**Definition:** Let *X* denote a finite set of points in a domain, Ω ⊂ ℝ^2^, and *d*_*i,j*_ represent the distance between points *x*_*i*_ and *x*_*j*_. For a given scale parameter *ϵ* ≥ 0, the VR complex *R*_*ϵ*_(*X*) is a simplicial complex defined as follows. A finite subset *σ* = {*x*_0_, *x*_1_, …, *x*_*k*_} ⊆ *X* forms a *k*-simplex in *R*_*ϵ*_(*X*) if and only if the pairwise distances between all points in *σ* satisfy *d*_*i,j*_ ≤ *ϵ* for all 0 ≤ *i, j* ≤ *k*. As the scale parameter *ϵ* increases, the VR complexes are nested, satisfying 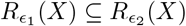 for any 0 ≤ *ϵ*_1_ ≤ *ϵ*_2_. This nested sequence forms a filtration of simplicial complexes, denoted by {*R*_*ϵ*_(*X*)}_*ϵ*≥0_.

To quantify these topological features, the homology groups *H*_*k*_(*R*_*ϵ*_(*X*)) of the VR complexes are computed for each dimension *k* as follows. Each *k*-simplex in *R*_*ϵ*_(*X*) there exists a boundary operator *∂*_*k*_ that maps it to a formal sum of its (*k* − 1)-dimensional faces. The boundary operators satisfy the composition property *∂*_*k*_ ∘ *∂*_*k*+1_ = 0, which ensures that the image of *∂*_*k*+1_ is contained within the kernel of *∂*_*k*_. The *k*-th homology group is then defined as the quotient

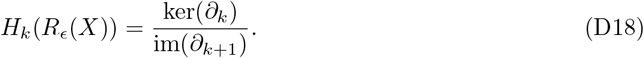

It captures the *k*-dimensional topological features, such as connected components (*k* = 0), loops (*k* = 1), and higher-dimensional voids. Persistent homology extends this computation across the filtration {*R*_*ϵ*_(*X*)} _*ϵ*≥0_. As *ϵ* increases, the boundary maps and the resulting homology groups change, leading to the birth and death of topological features.

**Application:** VR filtrations were used to identify crypt-like structures in Fig. 4F in the PROG population by restricting our attention to 0 and 1-dimensional features (*k* ≤ 1) and defining *d*_*i,j*_ as the minimum Euclidean distance between point objects *i* and *j*.

### D.4 Spatial relationships at tissue scales

#### D.4.1 Neighbourhood clustering

**Description:** Microenvironment clustering seeks to group objects with similar compositions of labels located within a local neighbourhood of an object. These methods are inspired by those deployed in [S3].

**Definition:** Let 𝒢 (*V, E*) be a network with nodes *v*_*i*_ ∈ *V* and edges *e*_*ij*_ ∈ *E* encode spatial relationships between nodes. Each node has a categorical label *c*_*i*_ ∈ {*C*_1_, …, *C*_*m*_} (i.e., its cell type). Let the microenvironment, 𝒩_*i*_, be the set of nodes about node *i* defined by a *k*-hop neighbourhood (see Table 1). For each neighbourhood, 𝒩_*i*_, the composition of each label category is computed to generate an observation matrix ***O*** ∈ ℕ^|*V* |*×m*^ where

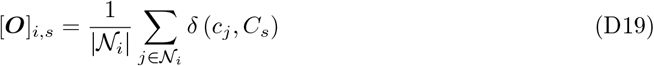

and

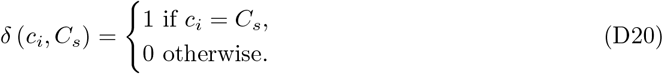

Row-wise clustering is performed on ***O*** to identify similar microenvironment compositions, which partition the nodes *v*_*i*_ ∈ *V* into disjoint sets *S*_*t*_.

Cluster representatives (i.e., the mean or medoid composition vector of the cluster) are extracted from each set *S*_*t*_, denoted by 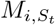.

Define the average composition of label *C*_*i*_ of the sample, *M*_*i*_, by

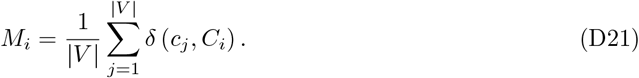

An microenvironment enrichment score, 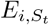, is then calculated by compare the representatives to the average composition,*M*_*i*_, define as a log-fold change

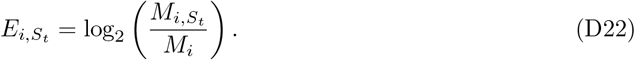

**Application:** In Fig. 5D, 𝒢 (*V, E*) is a Delaunay network (see Table 1) on cell centres with edge distance threshold of 30µm. A 3-hop neighbourhood was used to generate microenvironment sets about each node. Clustering was performed using *k*-means with *k* = 5.

#### D.4.2 Adjacency permutation test

**Description:** The adjacency permutation test quantifies the significance of labelled nodes being adjacent in a network. This non-parametric test involves repeatedly shuffling node labels to generate a null distribution of a test statistic under the hypothesis of no spatial association [S7].

**Definition:** Let 𝒢 (*V, E*) be a network with nodes *v*_*i*_ ∈ *V* and edges *e*_*ij*_ ∈ *E* where each node has a categorical label *c* ∈ {*C*_1_, …, *C*_*m*_}. For every pair of categorical labels *C*_*s*_ and *C*_*t*_, the number of observed edges, 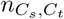, is calculated between nodes with these labels such that

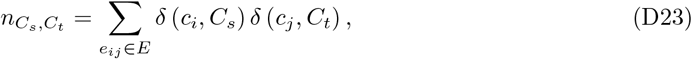

where *δ* (*c*_*i*_, *C*_*s*_) is defined in equation (D20). An observation matrix, 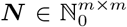, is constructed using each categorical pair count, 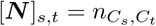, for 1 ≤*s, t* ≤*m*.

The node labels are then shuffled and the adjacent pair count procedure is repeated for *K* iterations, generating a null distribution of count matrices 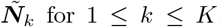 for 1 ≤ *k* ≤ *K*. The adjacency significance score for each pair of categorical labels, *S*_*s,t*_, is calculated as z-score of the observed to the null,

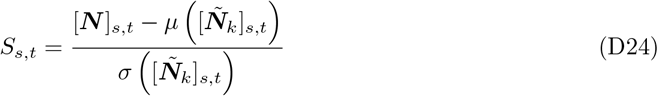

are the mean and standard deviation over all *k*.

**Application:** The Adjacency Permutation Test was used to determine the adjacency significance of the ME regions that are in close proximity in Fig. 5G. After converting spatially contiguous regions of points with identical ME labels into shapes (ME regions, Fig. 5F), a proximity network of these shapes was generated where edges between ME regions were drawn if they less than 5µm apart. This proximity network equipped with ME ID labels was then subjected to a Adjacency Permutation Test with 1000 random samples (*K* = 1000).

#### D.4.3 Local Getis-Ord

**Description:** The local Getis-Ord statistic 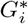 is measures spatial autocorrelation by relating spatial weights (e.g., distances between objects) with label values [S32].

**Definition:** Let ***x***_*i*_ ∈ ℝ^2^ represent the spatial position of the *i*-th object, 1 ≤ *i* ≤ *n*, and associated with a continuous label *l*_*i*_ ∈ ℝ. If *w*_*i,j*_ ∈ ℝ defines the spatial relationship between ***x***_*i*_ and ***x***_*j*_, then the local Getis-Ord statistic, 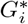, for object *i* is given by

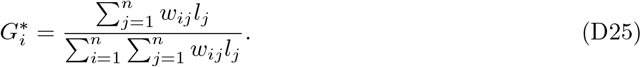

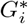 is the weighted average of label values adjacent to object *i*. The value of 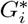 relative to the mean identifies whether an object belongs to a *hotspot* (+ve) or *coldspot* (-ve) for that label is typically reported as a z-score, *z*,

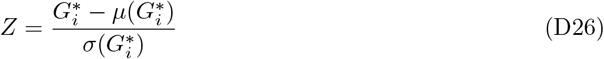

where *µ* and *σ* are the mean and standard deviation over all *i*, respectively.

**Application:** In Fig. 5I, the objects are hexagons, and we assign a weight function for the shapes *i* and *j* of the form

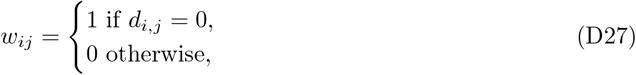

where *d*_*i,j*_ is the minimum distance between objects *i* and *j*. The labels *l*_*i*_ are the number of cells of PROG cells located within hexagon *i*.

#### D.4.4 Gaussian kernel density estimation

**Description:** Kernel density estimation (KDE) is a non-parametric method for estimating a probability density function from observations.

**Definition:** A Gaussian KDE uses Gaussian kernels centred on each observation, ***x***_*i*_, to generate a distribution over a domain Ω ⊂ ℝ^2^. The resultant probability density function, *P* (***x***), is determined by

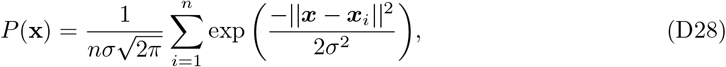

where *n* is the number of observations and *σ* is the bandwidth of the kernel. Increasing *σ* generates smoother representations of the observed point data, ***x***_*i*_.

**Application:** In Fig. 5J, we independently apply Gaussian kernels, centred on SMC, PROG, and STR1 cells, with *σ* = 0.05.

#### D.4.5 Wasserstein distance

**Description:** The 2-Wasserstein distance (referred to here as the Wasserstein distance), describes the minimum distance required to transform one distribution onto another (see [S41] for further details).

**Definition:** Consider two discrete probability distributions ***µ*** = (*µ*_1_, …, *µ*_*n*_) and ***ν*** = (*ν*_1_, …, *ν*_*m*_) with spatial locations ***x*** = (***x***_1_, …, ***x***_*n*_) and ***y*** = (***y***_1_, …, ***y***_*m*_) within a domain Ω ⊂ ℝ^2^, respectively.

Then the 2-Wasserstein distance is defined as

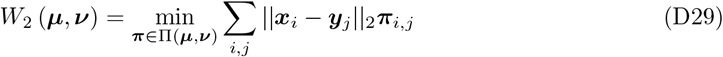

where || · ||_2_ is the *l*_2_-norm and

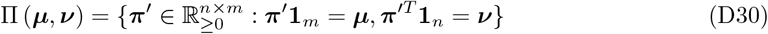

is the set of all possible transport plans from ***µ*** to ***ν***. The mapping ***π*** provides the optimal transport plan that transforms ***µ*** onto ***ν*** with the minimum work.

Solving for ***π*** is computationally inefficient for distributions in more than one dimension and therefore we employ the sliced method to determine ***π*** [S42]. We project ***µ*** and ***ν*** onto a one-dimensional line drawn through the barycentre of the distributions at an angle *θ U ∼* [0, *π*) and then determine the 1-dimensional Wasserstein determined,

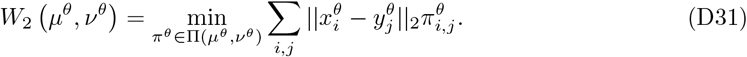

The Wasserstein distance of the full-dimension dataset is then approximated by the root average of these projected distances such that

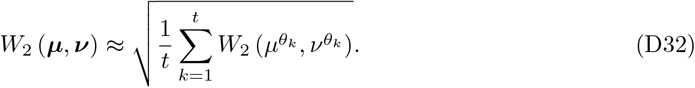

Critically, equation (D32) holds in equality in the limit *t* → ∞ [S41].

**Application:** In Figure 5k, *µ* and *ν* are weighted uniformly such that 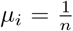 and 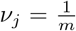 for all 1 ≤*i* ≤ *n* and 1 ≤ *j* ≤ *m*. In addition, *t* = 500 was selected as the number of sliced projections for sufficient coverage of the spatial data.

#### D.4.6 KL-Divergence

**Description:** The Kullback-Leibler (KL) divergence is a method for comparing the statistical distance between two probability distributions by measuring the entropy of a distribution relative to the other [S17].

**Definition:** Let *P* and *Q* be two probability distributions defined over the sample space. The KL divergence is defined as

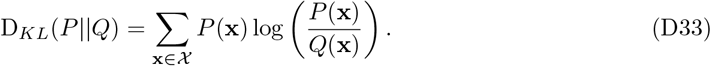

Thus, *D*_*KL*_(*P* || *Q*) measures the expected logarithmic difference between the probabilities *P* and *Q*, with the expectation taken under the distribution defined by *P*.

**Application:** In Figure 5k, *P* and *Q* are the Gaussian KDEs associated with of the point clouds of the corresponding SMC, PROG, and STR1 cells.

## Appendix E Supplementary information for multiscale spatial profiling of fibroblast-immune interactions in CRC

### E.1 Extended mouse information

All procedures were carried out in accordance with Home Office UK regulations and the Animals (Scientific Procedures) Act 1986. All mice were housed in individually ventilated cages at the animal unit either at the Functional Genetics Facility (Wellcome Centre for Human Genetics, University of Oxford). All mice were housed in a specific-pathogen-free (SPF) facility, with unrestricted access to food and water, and were not involved in any previous procedures. All strains used in this study were maintained on a C57BL/6J background for ≥6 generations. Mice were sacrificed when they reached humane-end points (exhibited anaemia, hunching and inactivity).

### E.2 Cell type annotation

Spatial transcriptomic data from four AKPT murine colorectal cancer (CRC) samples were provided as HDF5 files containing regions of interest and processed using Seurat (v5.2.0) in R (v4.3.3). Each sample was converted into a Seurat object, merged, collapsed into a single layer using `JoinLayers()`, and normalized using `LogNormalize` with a scale factor of 500 (Fig. E10A). Dimensionality reduction and clustering were performed using PCA and UMAP (dimensions 1–20), followed by initial clustering at a resolution of 0.1 (Fig. E10B). Clusters expressing epithelial markers (*Epcam, Cdh1*) were identified and removed(Figs. E10C), after which the remaining ‘non-epithelial’ cells were re-clustered at a resolution of 0.6 to identify stromal and immune populations (Fig. E10D).

For each of the resulting 22 clusters, aggregate scaled expression of fibroblast (*Pdgfra*) and immune-associated genes (*Ptprc, Cd68, Ms4a7, Trem2, Spp1, Clec9a, Cxcl2, Foxp3, Cd3e, Cd8a*, and *Mzb1*) was calculated (Figs. E10E-F). Clusters were annotated as fibroblast if *Pdgfra* expression was *>* 0, or immune if *Ptprc* expression was *>* 0. Clusters 3 and 4 expressed both markers but were assigned as fibroblast based on higher *Pdgfra* levels (cluster 3: *Pdgfra* = 2.30, *Ptprc* = 0.25; cluster 4: *Pdgfra* = 2.45, *Ptprc* = 0.03). This enabled a low-resolution classification of fibroblast, immune, and unlabelled populations.

Immune clusters were further subtyped by visualising scaled aggregate expression of immune markers using `ComplexHeatmap` (v2.18.0) (Fig. E10G). Clusters enriched for distinct immune gene signatures were annotated as specific immune cell types, resulting in the identification of 12 immune populations, with the remaining clusters left unlabelled (Fig. E10H).

### E.3 Extended analysis methods

#### Dimension reduction

Dimensionality reduction of transcriptional counts and multiscale spatial feature vectors was performed using UMAP for visualisation. For each two-dimensional embedding, we set *n neighbors*=100 for transcriptional data and *n neighbors*=150 for spatial feature data, using the *Manhattan* distance metric in both datasets.

#### Trajectory inference

To infer a sequential organisation of fibroblast types, we constructed a weighted minimum spanning tree (WMST) connecting the medoids of transcriptionally defined fibroblast clusters [S43]. Pairwise distances between medoids were calculated using the 1-norm (Manhattan distance) of their transcript count vectors. The WMST was computed using the `minimum_spanning_tree` function from `networkx` (v3.4.2), and projected into the UMAP space to visualise inferred cluster trajectories.

**Fig. E10:**
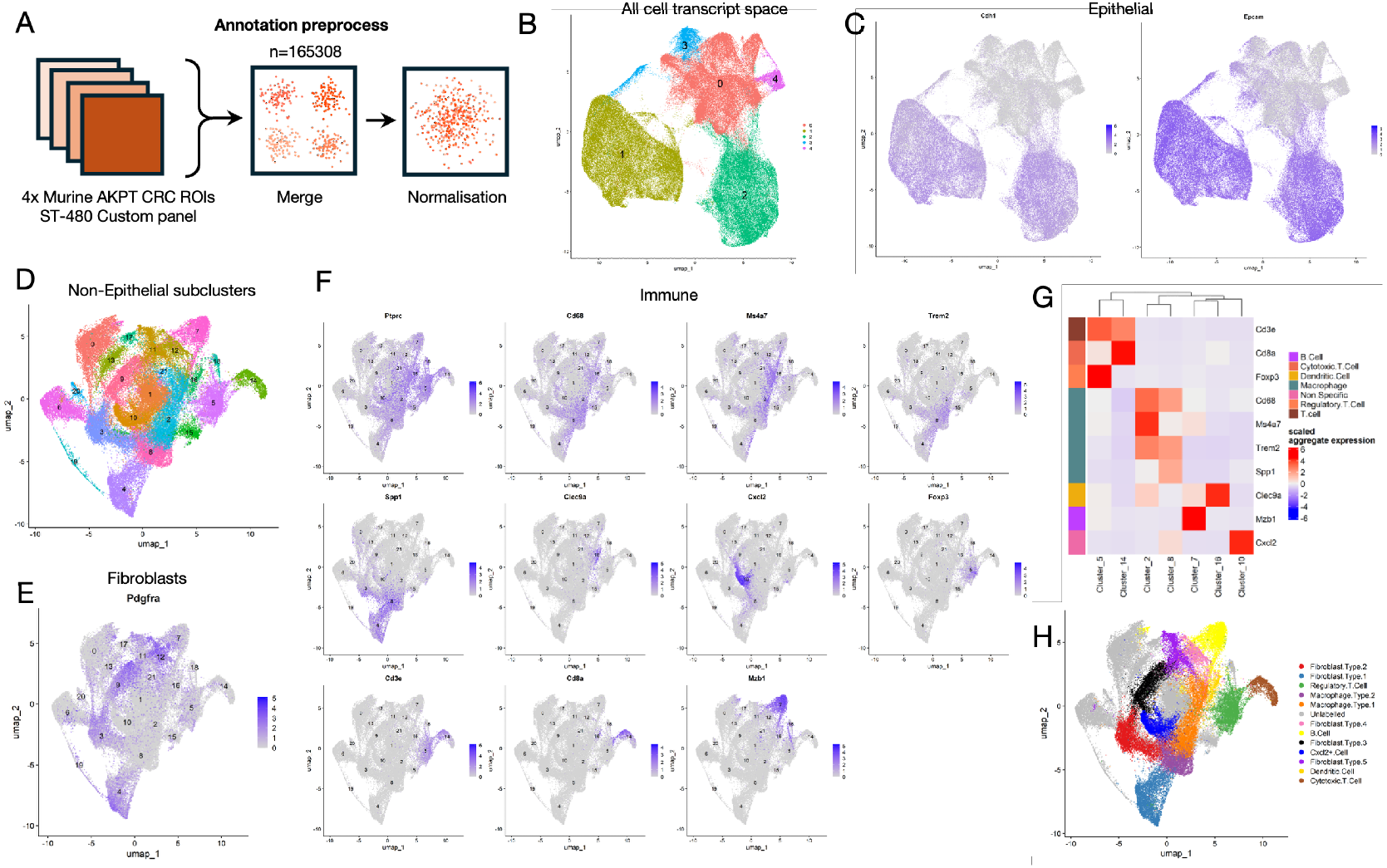
Annotation of murine AKPT colorectal tumours. A–H: Cell type annotation of four Xenium (480-gene custom panel) AKPT colorectal tumour samples. A: Schematic overview of the preprocessing workflow for cell type annotation. B: Two-dimensional UMAP projection of the 480-gene transcript space across all samples, coloured by coarse clustering. C: Expression of epithelial markers *Cdh1* and *Epcam* mapped onto the transcript UMAP from A. D: UMAP projection of transcript expression from non-epithelial cells (clusters 0, 3, and 4 from B), coloured by fine-grained clustering (clusters 0–21). E: Expression of the fibroblast marker *Pdgfra* on the non-epithelial UMAP projection from D. F: Expression of immune cell markers (*Ptprc, Cd68, Ms4a7, Trem2, Spp1, Clec9a, Cxcl2, Foxp3, Cd3e, Cd8a*, and *Mzb1*) on the non-epithelial UMAP projection from D. G: Heatmap showing hierarchical clustering of immune marker expression with supervised immune cell label annotations. H: Final cell type annotations for fibroblasts and immune cells overlaid on the non-epithelial transcript UMAP. Grey points indicate unclassified cells.

#### Generating MuSpAn domains

MuSpAn domains for each AKPT colorectal cancer sample were extracted directly from Xenium Explorer (v3) using the `xenium to domain` helper function from the `io` module of MuSpAn. This function automatically maps cell boundary geometries to their corresponding annotations. A pathology-derived region, annotated in Xenium Explorer (v3), was incorporated into the domain using the `add_shapes` function. Sample boundaries were estimated using an alpha shape (*α* = 100) via the `estimate_boundary` function. Cell centroids were computed using the `convert_objects` function, allowing each cell to be represented either as a shape or a point for downstream spatial analysis.

#### Tumour annotation analysis

To assess whether selected fibroblasts were located within tumour annotations, we used the `is A_contained_by_B` function from the `query` module of MuSpAn. Fibroblasts within tumour regions were assigned to an *in tumour* collection, and the proportion of each fibroblast type within this collection was calculated.

#### Cell-cell contact analysis

To analyse cell–cell contact, we first constructed a proximity network for each sample, representing physical interactions between adjacent cell boundaries. Networks were generated using the *proximity* method within the `generate_network` function of the `networks` module of MuSpAn, with contact defined as a distance of ≤1.5 *µ*m between cell boundaries. Each resulting network was stored within the corresponding sample domain for downstream analysis.

Contact adjacency correlations were assessed using the `adjacency_permutation_test` function in the `networks` module, with statistical significance threshold *α* = 0.05 and 500 randomised permutations on the contact networks previously constructed. This test was applied independently to each sample.

Topographical Correlation Maps (TCMs) were generated to visualise patterns of contact between immune and fibroblast populations within each sample. TCMs were computed using the `topographical correlation map` function from the `spatial_statistics` module of MuS-pAn, with interaction radius *r* = 20 *µ*m, Gaussian kernel variance *σ* = 25, and a maximum correlation threshold of *α* = 3. Further details on the `adjacency_permutation _test` and `topographical_correlation_map` are provided in Appendix D.

#### Neighbourhood analysis

To investigate immune neighbourhood composition surrounding selected fibroblasts, we use the `cluster_neighbourhoods` function from the `networks` module in MuSpAn. Neighbourhoods were defined as all nodes reachable within three edges (3-hop neighbourhoods; *k hop* = 3) from each fibroblast of interest, using the previously constructed cell boundary contact network.

For each fibroblast, the immune cell composition within its 3-hop neighbourhood was computed and square-root transformed across all samples. These transformed profiles were then clustered using *k* -means clustering (*cluster method* = `kmeans`) with *k* = 4 (*n clusters* = 4). Cluster enrichment was quantified by computing the *z*-score of the representative centroid of each cluster (*neighbourhood enrichment as* = `zscore`). Mathematical details for the `cluster_neighbourhoods` function are provided in Appendix D.

#### Spatial correlation at multiple scales

To assess spatial associations between selected fibroblast and immune cell populations across multiple length scales, we used the cross–Pair Correlation Function (cross-PCF). The cross-PCF was computed using the `cross _pair correlation_function` from the `spatial_statistics` module of MuSpAn. For all analyses, correlations were calculated up to a maximum radius of 500 *µ*m (*max R* = 500 *µ*m), with an annulus width of 30 *µ*m (*annulus width* = 30 *µ*m).

As a summary statistic, we computed the cumulative cross-PCF ratio, *C*_*AB*_(*R*), which captures the integrated spatial association between two populations up to radius *R*, defined as

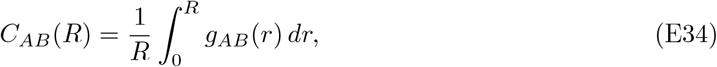

where *g*_*AB*_(*r*) is the cross-PCF between populations *A* and *B*, and *R* = 150 *µ*m. The cumulative ratio is normalised against the expected value under a spatial null model, preserving interpretation of the cross-PCF. Further mathematical details of the cross-PCF computation are provided in Appendix D.

#### Multiscale spatial feature vectors

To enable integrated analysis of the fibroblast spatial context, we constructed multiscale feature vectors for each fibroblast by combining outputs from the aforementioned analyses. For each fibroblast, local contact-based neighbourhood features (1-hop and 3-hop immune subtype counts), TCM value with each immune population at it’s location, and spatial association statistics derived from cross–pair correlation functions (cross-PCFs). Cross-PCF features included the cumulative spatial correlation up to 150 *µ*m, as well as values sampled at radii of 0, 100, 200, 300, and 400 *µ*m, capturing spatial interactions across multiple scales. The resultant pairwise computations between the fibroblast and immune subtypes produced a 87-dimensional vector for each fibroblast.

All features were computed independently for each fibroblast across all samples and combined into a single unified dataframe. To ensure comparability between features, each dimension was z-score normalised across all fibroblasts. For each fibroblast type, we computed the median of its multiscale feature vectors to obtain a representative profile. A WMST was then constructed using the 1-norm distance matrix between these fibroblast type medians. To visualise the inferred progression of fibroblast–immune interactions, both the fibroblast-level feature vectors and the WMST edges were projected into a shared two-dimensional space using the same UMAP embedding.

## References

[1] Dario Bressan, Giorgia Battistoni, and Gregory J. Hannon. “The dawn of spatial omics”. In: Science 381.6657 (Aug. 2023). Publisher: American Association for the Advancement of Science, eabq4964. DOI: 10.1126/science.abq4964. URL: https://www.science.org/doi/full/10.1126/science.abq4964 (visited on 11/18/2024).

[2] Jeffrey R. Moffitt, Emma Lundberg, and Holger Heyn. “The emerging landscape of spatial profiling technologies”. en. In: Nature Reviews Genetics 23.12 (Dec. 2022). Publisher: Nature Publishing Group, pp. 741–759. ISSN: 1471-0064. DOI: 10.1038/s41576-022-00515-3. URL: https://www.nature.com/articles/s41576-022-00515-3 (visited on 11/18/2024).

[3] Christopher R. Merritt et al. “Multiplex digital spatial profiling of proteins and RNA in fixed tissue”. en. In: Nature Biotechnology 38.5(May 2020). Publisher: Nature Publishing Group, pp. 586–599. ISSN: 1546-1696. DOI: 10.1038/s41587-020-0472-9. URL: https://www.nature.com/articles/s41587-020-0472-9 (visited on 11/18/2024).

[4] Guillaume Jacquemet et al. “The cell biologist’s guide to super-resolution microscopy”. In: Journal of Cell Science 133.11 (June 2020), jcs240713. ISSN: 0021-9533. DOI: 10.1242/jcs.240713. (visited on 11/18/2024).

[5] Katy Vandereyken et al. “Methods and applications for single-cell and spatial multi-omics”. en. In: Nature Reviews Genetics 24.8 (Aug. 2023). Publisher: Nature Publishing Group, pp. 494–515. ISSN: 1471-0064. DOI: 10.1038/s41576-023-00580-2. URL: https://www.nature.com/articles/s41576-023-00580-2 (visited on 11/18/2024).

[6] Shahar Alon et al. “Expansion sequencing: Spatially precise in situ transcriptomics in intact biological systems”. In: Science 371.6528 (Jan. 2021). Publisher: American Association for the Advancement of Science, eaax2656. DOI: 10.1126/science.aax2656. URL: https://www.science.org/doi/full/10.1126/science.aax2656 (visited on 11/18/2024).

[7] Jun-Han Su et al. “Genome-Scale Imaging of the 3D Organization and Transcriptional Activity of Chromatin”. English. In: Cell 182.6 (Sept. 2020). Publisher: Elsevier, 1641–1659.e26. ISSN: 0092-8674, 1097-4172. DOI: 10.1016/j.cell.2020.07.032. URL: https://www.cell.com/cell/abstract/S0092-8674(20)30940-5 (visited on 11/18/2024).

[8] Yodai Takei et al. “Integrated spatial genomics reveals global architecture of single nuclei”. en. In: Nature 590.7845 (Feb. 2021). Publisher: Nature Publishing Group, pp. 344–350. ISSN: 1476-4687. DOI: 10.1038/s41586-020-03126-2. URL: https://www.nature.com/articles/s41586-020-03126-2 (visited on 11/18/2024).

[9] Anjali Rao et al. “Exploring tissue architecture using spatial transcriptomics”. en. In: Nature 596.7871 (Aug. 2021). Publisher: Nature Publishing Group, pp. 211–220. ISSN: 1476-4687. DOI: 10.1038/s41586-021-03634-9. URL: https://www.nature.com/articles/s41586-021-03634-9 (visited on 11/18/2024).

[10] John W. Hickey et al. “Spatial mapping of protein composition and tissue organization: a primer for multiplexed antibody-based imaging”. en. In: Nature Methods 19.3 (Mar. 2022). Publisher: Nature Publishing Group, pp. 284–295. ISSN: 1548-7105. DOI: 10.1038/s41592-021-01316-y. URL: https://www.nature.com/articles/s41592-021-01316-y (visited on 11/18/2024).

[11] Yury Goltsev et al. “Deep Profiling of Mouse Splenic Architecture with CODEX Multiplexed Imaging”. In: Cell 174.4 (Aug. 2018), 968–981.e15. ISSN: 0092-8674. DOI: 10.1016/j.cell.2018.07.010. URL: https://www.sciencedirect.com/science/article/pii/S0092867418309048 (visited on 03/08/2024).

[12] Michael Angelo et al. “Multiplexed ion beam imaging of human breast tumors”. en. In: Nature Medicine 20.4 (Apr. 2014). Publisher: Nature Publishing Group, pp. 436–442. ISSN: 1546-170X. DOI: 10.1038/nm.3488. URL: https://www.nature.com/articles/nm.3488 (visited on 11/18/2024).

[13] Charlotte Giesen et al. “Highly multiplexed imaging of tumor tissues with subcellular resolution by mass cytometry”. en. In: Nature Methods 11.4 (Apr. 2014). Publisher: Nature Publishing Group, pp. 417–422. ISSN: 1548-7105. DOI: 10.1038/nmeth.2869. URL: https://www.nature.com/articles/nmeth.2869 (visited on 03/12/2024).

[14] Peter Bankhead et al. “QuPath: Open source software for digital pathology image analysis”. en. In: Scientific Reports 7.1 (Dec. 2017). Publisher: Nature Publishing Group, p. 16878. ISSN: 2045-2322. DOI: 10.1038/s41598-017-17204-5. URL: https://www.nature.com/articles/s41598-017-17204-5 (visited on 11/04/2024).

[15] Noah F. Greenwald et al. “Whole-cell segmentation of tissue images with human-level performance using large-scale data annotation and deep learning”. en. In: Nature Biotechnology 40.4 (Apr. 2022). Publisher: Nature Publishing Group, pp. 555–565. ISSN: 1546-1696. DOI: 10.1038/s41587-021-01094-0. URL: https://www.nature.com/articles/s41587-021-01094-0 (visited on 11/15/2024).

[16] Stuart Berg et al. “ilastik: interactive machine learning for (bio)image analysis”. en. In: Nature Methods 16.12 (Dec. 2019). Publisher: Nature Publishing Group, pp. 1226–1232. ISSN: 1548-7105. DOI: 10.1038/s41592-019-0582-9. URL: https://www.nature.com/articles/s41592-019-0582-9 (visited on 11/15/2024).

[17] Denis Schapiro et al. “histoCAT: analysis of cell phenotypes and interactions in multiplex image cytometry data”. en. In: Nature Methods 14.9 (Sept. 2017). Publisher: Nature Publishing Group, pp. 873–876. ISSN: 1548-7105. DOI: 10.1038/nmeth.4391. URL: https://www.nature.com/articles/nmeth.4391 (visited on 11/15/2024).

[18] Zhiyuan Yuan et al. “Benchmarking spatial clustering methods with spatially resolved transcriptomics data”. en. In: Nature Methods 21.4 (Apr. 2024). Publisher: Nature Publishing Group, pp. 712–722. ISSN: 1548-7105. DOI: 10.1038/s41592-024-02215-8. URL: https://www.nature.com/articles/s41592-024-02215-8 (visited on 11/18/2024).

[19] Luca Marconato et al. “SpatialData: an open and universal data framework for spatial omics”. en. In: Nature Methods (Mar. 2024). Publisher: Nature Publishing Group, pp. 1–5. ISSN: 1548-7105. DOI: 10.1038/s41592-024-02212-x. URL: https://www.nature.com/articles/s41592-024-02212-x (visited on 11/18/2024).

[20] Chi-Li Chiu, Nathan Clack, and the napari community. “napari: a Python Multi-Dimensional Image Viewer Platform for the Research Community”. In: Microscopy and Microanalysis 28.S1 (Aug. 2022), pp. 1576–1577. ISSN: 1431-9276. DOI: 10.1017/S1431927622006328. (visited on 11/18/2024).

[21] Thibaut Goldsborough et al. InstanSeg: an embedding-based instance segmentation algorithm optimized for accurate, efficient and portable cell segmentation. arXiv:2408.15954. Aug. 2024. DOI: 10.48550/arXiv.2408.15954. URL: http://arxiv.org/abs/2408.15954 (visited on 11/04/2024).

[22] Daniela Bruni, Helen K. Angell, and Jérôme Galon. “The immune contexture and Immunoscore in cancer prognosis and therapeutic efficacy”. en. In: Nature Reviews Cancer 20.11 (Nov. 2020). Publisher: Nature Publishing Group, pp. 662–680. ISSN: 1474-1768. DOI: 10.1038/s41568-020-0285-7. URL: https://www.nature.com/articles/s41568-020-0285-7 (visited on 11/29/2024).

[23] Raghu Kalluri and Valerie S. LeBleu. “The biology, function, and biomedical applications of exosomes”. In: Science 367.6478 (Feb. 2020). Publisher: American Association for the Advancement of Science, eaau6977. DOI: 10.1126/science.aau6977. URL: https://www.science.org/doi/full/10.1126/science.aau6977 (visited on 08/11/2025).

[24] Joshua A. Bull et al. Integrating diverse statistical methods to analyse stage-discriminatory cell interactions in colorectal neoplasia. en. Pages: 2024.06.02.597010 Section: New Results. June 2024. DOI: 10.1101/2024.06.02.597010. URL: https://www.biorxiv.org/content/10.1101/2024.06.02.597010v1 (visited on 11/04/2024).

[25] Christian M. Schürch et al. “Coordinated Cellular Neighborhoods Orchestrate Antitumoral Immunity at the Colorectal Cancer Invasive Front”. English. In: Cell 182.5 (Sept. 2020). Publisher: Elsevier, 1341–1359.e19. ISSN: 0092-8674, 1097-4172. DOI: 10.1016/j.cell.2020.07.005. URL: https://www.cell.com/cell/abstract/S0092-8674(20)30870-9 (visited on 11/27/2023).

[26] Eloise Withnell and Maria Secrier. “SpottedPy quantifies relationships between spatial transcriptomic hotspots and uncovers environmental cues of epithelial-mesenchymal plasticity in breast cancer”. In: Genome Biology 25.1 (Nov. 2024), p. 289. ISSN: 1474-760X. DOI: 10.1186/s13059-024-03428-y. (visited on 11/25/2024).

[27] Yuzhou Feng et al. “Spatial analysis with SPIAT and spaSim to characterize and simulate tissue microenvironments”. en. In: Nature Communications 14.1(May 2023). Publisher: Nature Publishing Group, p. 2697. ISSN: 2041-1723. DOI: 10.1038/s41467-023-37822-0. URL: https://www.nature.com/articles/s41467-023-37822-0 (visited on 03/25/2024).

[28] Clarence K. Mah et al. “Bento: a toolkit for subcellular analysis of spatial transcriptomics data”. In: Genome Biology 25.1 (Apr. 2024), p. 82. ISSN: 1474-760X. DOI: 10.1186/s13059-024-03217-7. (visited on 08/22/2024).

[29] Ajit J. Nirmal and Peter K. Sorger. SCIMAP: A Python Toolkit for Integrated Spatial Analysis of Multiplexed Imaging Data. arXiv:2405.02076 [q-bio]. May 2024. DOI: 10.48550/arXiv.2405.02076. URL: http://arxiv.org/abs/2405.02076 (visited on 05/20/2024).

[30] Raman Sethi et al. “ezSingleCell: an integrated one-stop single-cell and spatial omics analysis platform for bench scientists”. en. In: Nature Communications 15.1(July 2024). Publisher: Nature Publishing Group, p. 5600. ISSN: 2041-1723. DOI: 10.1038/s41467-024-48188-2. URL: https://www.nature.com/articles/s41467-024-48188-2 (visited on 08/22/2024).

[31] Praveen Weeratunga et al. “Single cell spatial analysis reveals inflammatory foci of immature neutrophil and CD8 T cells in COVID-19 lungs”. en. In: Nature Communications 14.1 (Nov. 2023). Number: 1 Publisher: Nature Publishing Group, p. 7216. ISSN: 2041-1723. DOI: 10.1038/s41467-023-42421-0. URL: https://www.nature.com/articles/s41467-023-42421-0 (visited on 11/27/2023).

[32] Giovanni Palla et al. “Squidpy: a scalable framework for spatial omics analysis”. en. In: Nature Methods 19.2 (Feb. 2022). Publisher: Nature Publishing Group, pp. 171–178. ISSN: 1548-7105. DOI: 10.1038/s41592-021-01358-2. URL: https://www.nature.com/articles/s41592-021-01358-2 (visited on 05/20/2024).

[33] Isaac Virshup et al. “The scverse project provides a computational ecosystem for single-cell omics data analysis”. en. In: Nature Biotechnology 41.5(May 2023). Publisher: Nature Publishing Group, pp. 604–606. ISSN: 1546-1696. DOI: 10.1038/s41587-023-01733-8. URL: https://www.nature.com/articles/s41587-023-01733-8 (visited on 11/04/2024).

[34] Philip J. Clark and Francis C. Evans. “Distance to Nearest Neighbor as a Measure of Spatial Relationships in Populations”. en. In: Ecology 35.4 (Oct. 1954), pp. 445–453. ISSN: 00129658. DOI: 10.2307/1931034. URL: http://doi.wiley.com/10.2307/1931034 (visited on 11/26/2024).

[35] Yuxuan Hu et al. “Unsupervised and supervised discovery of tissue cellular neighborhoods from cell phenotypes”. en. In: Nature Methods 21.2 (Feb. 2024). Publisher: Nature Publishing Group, pp. 267–278. ISSN: 1548-7105. DOI: 10.1038/s41592-023-02124-2. URL: https://www.nature.com/articles/s41592-023-02124-2 (visited on 11/27/2024).

[36] Doron Haviv et al. “The covariance environment defines cellular niches for spatial inference”. en. In: Nature Biotechnology (Apr. 2024). Publisher: Nature Publishing Group, pp. 1–12. ISSN: 1546-1696. DOI: 10.1038/s41587-024-02193-4. URL: https://www.nature.com/articles/s41587-024-02193-4 (visited on 11/27/2024).

[37] Gaurav Malviya et al. “Noninvasive Stratification of Colon Cancer by Multiplex PET Imaging”. en. In: Clinical Cancer Research 30.8 (Apr. 2024), pp. 1518–1529. ISSN: 1078-0432, 1557-3265. DOI: 10.1158/1078-0432.CCR-23-1063. URL: https://aacrjournals.org/clincancerres/article/30/8/1518/742132/Noninvasive-Stratification-of-Colon-Cancer-by (visited on 08/20/2025).

[38] Yunhe Liu et al. “Conserved spatial subtypes and cellular neighborhoods of cancer-associated fibroblasts revealed by single-cell spatial multi-omics”. English. In: Cancer Cell 43.5(May 2025). Publisher: Elsevier, 905–924.e6. ISSN: 1535-6108, 1878-3686. DOI: 10.1016/j.ccell.2025. URL: https://www.cell.com/cancer-cell/abstract/S1535-6108(25)00083-2 (visited on 08/11/2025).

[39] Justin Guinney et al. “The consensus molecular subtypes of colorectal cancer”. en. In: Nature Medicine 21.11 (Nov. 2015). Publisher: Nature Publishing Group, pp. 1350–1356. ISSN: 1546-170X. DOI: 10.1038/nm.3967. URL: https://www.nature.com/articles/nm.3967 (visited on 08/11/2025).

[40] Sophie Mouillet-Richard et al. “Clinical Challenges of Consensus Molecular Subtype CMS4 Colon Cancer in the Era of Precision Medicine”. In: Clinical Cancer Research 30.11 (June 2024), pp. 2351–2358. ISSN: 1078-0432. DOI: 10.1158/1078-0432.CCR-23-3964. (visited on 08/21/2025).

[41] David Sidak et al. “Interpretable machine learning methods for predictions in systems biology from omics data”. English. In: Frontiers in Molecular Biosciences 9 (Oct. 2022). Publisher: Frontiers. ISSN: 2296-889X. DOI: 10.3389/fmolb.2022.926623. URL: https://www.frontiersin. org/journals/molecular-biosciences/articles/10.3389/fmolb.2022.926623/full (visited on 11/27/2024).

[42] Oliver Vipond et al. “Multiparameter persistent homology landscapes identify immune cell spatial patterns in tumors”. In: Proceedings of the National Academy of Sciences 118.41 (Oct. 2021). Publisher: Proceedings of the National Academy of Sciences, e2102166118. DOI: 10.1073/pnas.2102166118. URL: https://www.pnas.org/doi/abs/10.1073/pnas.2102166118 (visited on 11/27/2023).

[43] Iris H. R. Yoon et al. Deciphering the diversity and sequence of extracellular matrix and cellular spatial patterns in lung adenocarcinoma using topological data analysis. en. Pages: 2024.01.05.574362 Section: New Results. Jan. 2024. DOI: 10.1101/2024.01.05.574362. URL: https://www.biorxiv.org/content/10.1101/2024.01.05.574362v2 (visited on 02/28/2024).

[44] Isaac Virshup et al. “anndata: Access and store annotated data matrices”. en. In: Journal of Open Source Software 9.101 (Sept. 2024), p. 4371. ISSN: 2475-9066. DOI: 10.21105/joss.04371. URL: https://joss.theoj.org/papers/10.21105/joss.04371 (visited on 11/04/2024).

[45] John D. Hunter. “Matplotlib: A 2D Graphics Environment”. In: Computing in Science & Engineering 9.3 (May 2007). Conference Name: Computing in Science & Engineering, pp. 90–95. ISSN: 1558-366X. DOI: 10.1109/MCSE.2007.55. URL: https://ieeexplore.ieee.org/document/4160265 (visited on 11/12/2024).

[46] Ulrich Bauer. “Ripser: efficient computation of Vietoris–Rips persistence barcodes”. en. In: Journal of Applied and Computational Topology 5.3 (Sept. 2021), pp. 391–423. ISSN: 2367-1734. DOI: 10.1007/s41468-021-00071-5. (visited on 03/06/2024).

[47] Pauli Virtanen et al. “SciPy 1.0: fundamental algorithms for scientific computing in Python”. en. In: Nature Methods 17.3 (Mar. 2020). Publisher: Nature Publishing Group, pp. 261–272. ISSN: 1548-7105. DOI: 10.1038/s41592-019-0686-2. URL: https://www.nature.com/articles/s41592-019-0686-2 (visited on 05/17/2024).

[48] Aric A. Hagberg, Daniel A. Schult, and Pieter J. Swart. “Exploring Network Structure, Dynamics, and Function using NetworkX”. In: Proceedings of the 7th Python in Science Conference. Ed. by Gaël Varoquaux, Travis Vaught, and Jarrod Millman. Pasadena, CA USA, 2008, pp. 11–15.

[49] Andrzej Kwinta and Joanna Bac-Bronowicz. “Regular polygons in 2D objects shape description”. en. In: Geomatics, Landmanagement and Landscape 4 (2020). ISSN: 2300-1496. DOI: 10.15576/GLL/2020.4.43. URL: https://repo.ur.krakow.pl/info/article/UR62ad65e95a3a45efb63ae1d773088563/ (visited on 11/29/2024).

[50] Marc Barthélemy. “Spatial networks”. In: Physics Reports 499.1 (Feb. 2011), pp. 1–101. ISSN: 0370-1573. DOI: 10.1016/j.physrep.2010.11.002. URL: https://www.sciencedirect.com/science/article/pii/S037015731000308X (visited on 11/29/2024).

[51] Naia Morueta-Holme et al. “A network approach for inferring species associations from co-occurrence data”. en. In: Ecography 39.12 (2016). eprint: https://onlinelibrary.wiley.com/doi/pdf/10.1111/ecog.01892, pp. 1139–1150. ISSN: 1600-0587. DOI: 10.1111/ecog.01892. URL: https://onlinelibrary.wiley.com/doi/abs/10.1111/ecog.01892 (visited on 11/27/2023).

[52] Baddeley, A., Rubak, E., and Turner, R. Spatial Point Patterns: Methodology and Applications with R. 1st Edition. Boca Raton, London, New York: CRC Press, Taylor & Francis Group, 2016. URL: https://www.routledge.com/Spatial-Point-Patterns-Methodology-and-Applications-with-R/Baddeley-Rubak-Turner/p/book/9781482210200.

[53] Mark M. Fredrickson and Yuguo Chen. “Permutation and randomization tests for network analysis”. In: Social Networks 59 (Oct. 2019), pp. 171–183. ISSN: 0378-8733. DOI: 10.1016/j.socnet.2019.08.001. URL: https://www.sciencedirect.com/science/article/pii/S0378873318303381 (visited on 11/29/2024).

[54] Arthur Getis and J. K. Ord. “The Analysis of Spatial Association by Use of Distance Statistics”. en. In: Geographical Analysis 24.3 (1992). eprint: https://onlinelibrary.wiley.com/doi/pdf/10.1111/j.1538-4632.1992.tb00261.x, pp. 189–206. ISSN: 1538-4632. DOI: 10.1111/j.1538-4632.1992.tb00261.x. URL: https://onlinelibrary.wiley.com/doi/abs/10.1111/j.1538-4632.1992.tb00261.x (visited on 11/26/2024).

[55] S. Kullback and R. A. Leibler. “On Information and Sufficiency”. In: The Annals of Mathematical Statistics 22.1 (Mar. 1951). Publisher: Institute of Mathematical Statistics, pp. 79–86. ISSN: 0003-4851, 2168-8990. DOI: 10.1214/aoms/1177729694. URL: https://projecteuclid.org/journals/annals-of-mathematical-statistics/volume-22/issue-1/On-Information-and-Sufficiency/10.1214/aoms/1177729694.full (visited on 11/19/2024).

[56] Joshua A. Bull et al. “Extended correlation functions for spatial analysis of multiplex imaging data”. en. In: Biological Imaging (Feb. 2024). DOI: 10.1017/S2633903X24000011. (visited on 12/04/2023).

## Supplementary Information References

[S1] Joshua A. Bull et al. “Combining multiple spatial statistics enhances the description of immune cell localisation within tumours”. en. In: Scientific Reports 10.1 (Oct. 2020). Number: 1 Publisher: Nature Publishing Group, p. 18624. ISSN: 2045-2322. DOI: 10.1038/s41598-020-75180-9. URL: https://www.nature.com/articles/s41598-020-75180-9 (visited on 11/27/2023).

[S2] Praveen Weeratunga et al. “Single cell spatial analysis reveals inflammatory foci of immature neutrophil and CD8 T cells in COVID-19 lungs”. en. In: Nature Communications 14.1 (Nov. 2023). Number: 1 Publisher: Nature Publishing Group, p. 7216. ISSN: 2041-1723. DOI: 10.1038/s41467-023-42421-0. URL: https://www.nature.com/articles/s41467-023-42421-0 (visited on 11/27/2023).

[S3] Christian M. Schürch et al. “Coordinated Cellular Neighborhoods Orchestrate Antitumoral Immunity at the Colorectal Cancer Invasive Front”. English. In: Cell 182.5 (Sept. 2020). Publisher: Elsevier, 1341–1359.e19. ISSN: 0092-8674, 1097-4172. DOI: 10.1016/j.cell.2020.07.005. URL: https://www.cell.com/cell/abstract/S0092-8674(20)30870-9 (visited on 11/27/2023).

[S4] Oliver Vipond et al. “Multiparameter persistent homology landscapes identify immune cell spatial patterns in tumors”. In: Proceedings of the National Academy of Sciences 118.41 (Oct. 2021). Publisher: Proceedings of the National Academy of Sciences, e2102166118. DOI: 10.1073/pnas.2102166118. URL: https://www.pnas.org/doi/abs/10.1073/pnas.2102166118 (visited on 11/27/2023).

[S5] Praveen Weeratunga et al. Temporo-spatial cellular atlas of the regenerating alveolar niche in idiopathic pulmonary fibrosis. en. Pages: 2024.04.10.24305440. Apr. 2024. DOI: 10.1101/2024.04.10.24305440. URL: https://www.medrxiv.org/content/10.1101/2024.04.10.24305440v1 (visited on 04/29/2024).

[S6] Joshua A. Bull et al. Integrating diverse statistical methods to analyse stage-discriminatory cell interactions in colorectal neoplasia. en. Pages: 2024.06.02.597010 Section: New Results. June 2024. DOI: 10.1101/2024.06.02.597010. URL: https://www.biorxiv.org/content/10.1101/2024.06.02.597010v1 (visited on 11/04/2024).

[S7] Mark M. Fredrickson and Yuguo Chen. “Permutation and randomization tests for network analysis”. In: Social Networks 59 (Oct. 2019), pp. 171–183. ISSN: 0378-8733. DOI: 10.1016/j.socnet.2019.08.001. URL: https://www.sciencedirect.com/science/article/pii/S0378873318303381 (visited on 11/29/2024).

[S8] Philip J. Clark and Francis C. Evans. “Distance to Nearest Neighbor as a Measure of Spatial Relationships in Populations”. en. In: Ecology 35.4 (Oct. 1954), pp. 445–453. ISSN: 00129658. DOI: 10.2307/1931034. (visited on 11/26/2024).

[S9] Joshua A. Bull et al. “Extended correlation functions for spatial analysis of multiplex imaging data”. en. In: Biological Imaging (Feb. 2024). DOI: 10.1017/S2633903X24000011. (visited on 12/04/2023).

[S10] Carlo C. Maley et al. “An ecological measure of immune-cancer colocalization as a prognostic factor for breast cancer”. en. In: Breast Cancer Research 17.1 (Sept. 2015), p. 131. ISSN: 1465-542X. DOI: 10.1186/s13058-015-0638-4. (visited on 11/19/2024).

[S11] Muhammad Shaban et al. “A Novel Digital Score for Abundance of Tumour Infiltrating Lymphocytes Predicts Disease Free Survival in Oral Squamous Cell Carcinoma”. en. In: Scientific Reports 9.1 (Sept. 2019). Publisher: Nature Publishing Group, p. 13341. ISSN: 2045-2322. DOI:10.1038/s41598-019-49710-z. URL: https://www.nature.com/articles/s41598-019-49710-z (visited on 11/19/2024).

[S12] B. D. Ripley. “Modelling Spatial Patterns”. en. In: Journal of the Royal Statistical Society: Series B (Methodological) 39.2 (1977). eprint: https://onlinelibrary.wiley.com/doi/pdf/10.1111/j.2517-6161.1977.tb01615.x, pp. 172–192. ISSN: 2517-6161. DOI: 10.1111/j.2517-6161.1977.tb01615.x. URL: https://onlinelibrary.wiley.com/doi/abs/10.1111/j.2517-6161.1977.tb01615.x (visited on 02/22/2024).

[S13] H. Edelsbrunner, D. Kirkpatrick, and R. Seidel. “On the shape of a set of points in the plane”. In: IEEE Transactions on Information Theory 29.4(July 1983). Conference Name: IEEE Transactions on Information Theory, pp. 551–559. ISSN: 1557-9654. DOI: 10.1109/TIT.1983.1056714. URL: https://ieeexplore.ieee.org/document/1056714 (visited on 03/06/2024).

[S14] Clarence K. Mah et al. “Bento: a toolkit for subcellular analysis of spatial transcriptomicsdata”. In: Genome Biology 25.1 (Apr. 2024), p. 82. ISSN: 1474-760X. DOI: 10.1186/s13059-024-03217-7. (visited on 08/22/2024).

[S15] Yuzhou Feng et al. “Spatial analysis with SPIAT and spaSim to characterize and simulate tissue microenvironments”. en. In: Nature Communications 14.1(May 2023). Publisher: Nature Publishing Group, p. 2697. ISSN: 2041-1723. DOI: 10.1038/s41467-023-37822-0. URL: https://www.nature.com/articles/s41467-023-37822-0 (visited on 03/25/2024).

[S16] Pauli Virtanen et al. “SciPy 1.0: fundamental algorithms for scientific computing in Python”. en. In: Nature Methods 17.3 (Mar. 2020). Publisher: Nature Publishing Group, pp. 261–272. ISSN: 1548-7105. DOI: 10.1038/s41592-019-0686-2. URL: https://www.nature.com/articles/s41592-019-0686-2 (visited on 05/17/2024).

[S17] S. Kullback and R. A. Leibler. “On Information and Sufficiency”. In: The Annals of Mathematical Statistics 22.1 (Mar. 1951). Publisher: Institute of Mathematical Statistics, pp. 79–86. ISSN: 0003-4851, 2168-8990. DOI: 10.1214/aoms/1177729694. URL: https://projecteuclid.org/journals/annals-of-mathematical-statistics/volume-22/issue-1/On-Information-and-Sufficiency/10.1214/aoms/1177729694.full (visited on 11/19/2024).

[S18] Cédric Villani. “The Wasserstein distances”. en. In: Optimal Transport: Old and New. Ed. by Cédric Villani. Grundlehren der mathematischen Wissenschaften. Berlin, Heidelberg: Springer, 2009, pp. 93–111. ISBN: 978-3-540-71050-9. DOI: 10.1007/978-3-540-71050-96. (visited on 03/06/2024).

[S19] M. G. Sreelekha, K. Krishnamurthy, and M. V. L. R. Anjaneyulu. “Interaction between Road Network Connectivity and Spatial Pattern”. In: Procedia Technology. International Conference on Emerging Trends in Engineering, Science and Technology (ICETEST - 2015) 24 (Jan. 2016), pp. 131–139. ISSN: 2212-0173. DOI: 10.1016/j.protcy.2016.05.019. URL: https://www.sciencedirect.com/science/article/pii/S2212017316301025 (visited on 11/29/2024).

[S20] Marc Barthélemy. “Spatial networks”. In: Physics Reports 499.1 (Feb. 2011), pp. 1–101. ISSN: 0370-1573. DOI: 10.1016/j.physrep.2010.11.002. URL: https://www.sciencedirect.com/science/article/pii/S037015731000308X (visited on 11/29/2024).

[S21] Akrati Saxena and Sudarshan Iyengar. Centrality Measures in Complex Networks: A Survey. arXiv:2011.07190. Nov. 2020. DOI: 10.48550/arXiv.2011.07190. URL: http://arxiv.org/abs/2011.07190 (visited on 11/29/2024).

[S22] Caleb R. Stoltzfus et al. “CytoMAP: A Spatial Analysis Toolbox Reveals Features of Myeloid Cell Organization in Lymphoid Tissues”. en. In: Cell Reports 31.3 (Apr. 2020), p. 107523. ISSN: 22111247. DOI: 10.1016/j.celrep.2020.107523. URL: https://linkinghub.elsevier.com/retrieve/pii/S221112472030423X (visited on 11/27/2023).

[S23] Vincent D. Blondel et al. “Fast unfolding of communities in large networks”. en. In: Journal of Statistical Mechanics: Theory and Experiment 2008.10 (Oct. 2008), P10008. ISSN: 1742-5468. DOI: 10.1088/1742-5468/2008/10/P10008. (visited on 11/29/2024).

[S24] Zhenqin Wu et al. “Graph deep learning for the characterization of tumour microenvironments from spatial protein profiles in tissue specimens”. en. In: Nature Biomedical Engineering 6.12 (Dec. 2022). Publisher: Nature Publishing Group, pp. 1435–1448. ISSN: 2157-846X. DOI: 10.1038/s41551-022-00951-w. URL: https://www.nature.com/articles/s41551-022-00951-w (visited on 03/25/2024).

[S25] Ajit J. Nirmal and Peter K. Sorger. SCIMAP: A Python Toolkit for Integrated Spatial Analysis of Multiplexed Imaging Data. arXiv:2405.02076 [q-bio]. May 2024. DOI: 10.48550/arXiv.2405.02076. URL: http://arxiv.org/abs/2405.02076 (visited on 05/20/2024).

[S26] Giovanni Palla et al. “Squidpy: a scalable framework for spatial omics analysis”. en. In: Nature Methods 19.2 (Feb. 2022). Publisher: Nature Publishing Group, pp. 171–178. ISSN: 1548-7105. DOI: 10.1038/s41592-021-01358-2. URL: https://www.nature.com/articles/s41592-021-01358-2 (visited on 05/20/2024).

[S27] Peter Wills and François G. Meyer. “Metrics for graph comparison: A practitioner’s guide”. en. In: PLOS ONE 15.2 (Feb. 2020). Publisher: Public Library of Science, e0228728. ISSN: 1932-6203. DOI: 10.1371/journal.pone.0228728. URL: https://journals.plos.org/plosone/article?id=10.1371/journal.pone.0228728 (visited on 11/29/2024).

[S28] Naia Morueta-Holme et al. “A network approach for inferring species associations from co-occurrence data”. en. In: Ecography 39.12 (2016). eprint: https://onlinelibrary.wiley.com/doi/pdf/10.1111/ecog.01892, pp. 1139–1150. ISSN: 1600-0587. DOI: 10.1111/ecog.01892. URL: https://onlinelibrary.wiley.com/doi/abs/10.1111/ecog.01892 (visited on 11/27/2023).

[S29] Baddeley, A., Rubak, E., and Turner, R. Spatial Point Patterns: Methodology and Applications with R. 1st Edition. Boca Raton, London, New York: CRC Press, Taylor & Francis Group, 2016. URL: https://www.routledge.com/Spatial-Point-Patterns-Methodology-and-Applications-with-R/Baddeley-Rubak-Turner/p/book/9781482210200.

[S30] Eloise Withnell and Maria Secrier. “SpottedPy quantifies relationships between spatial transcriptomic hotspots and uncovers environmental cues of epithelial-mesenchymal plasticity in breast cancer”. In: Genome Biology 25.1 (Nov. 2024), p. 289. ISSN: 1474-760X. DOI: 10.1186/s13059-024-03428-y. (visited on 11/25/2024).

[S31] Oscar E. Ospina et al. spatialGE: A user-friendly web application to democratize spatial transcriptomics analysis. en. Pages: 2024.06.27.601050 Section: New Results. July 2024. DOI: 10.1101/2024.06.27.601050. URL: https://www.biorxiv.org/content/10.1101/2024.06.27.601050v1 (visited on 08/22/2024).

[S32] Arthur Getis and J. K. Ord. “The Analysis of Spatial Association by Use of Distance Statistics”. en. In: Geographical Analysis 24.3 (1992). eprint: https://onlinelibrary.wiley.com/doi/pdf/10.1111/j.1538-4632.1992.tb00261.x, pp. 189–206. ISSN: 1538-4632. DOI: 10.1111/j.1538-4632.1992.tb00261.x. URL: https://onlinelibrary.wiley.com/doi/abs/10.1111/j.1538-4632.1992.tb00261.x (visited on 11/26/2024).

[S33] M. N. M. van Lieshout and A. J. Baddeley. “A nonparametric measure of spatial interaction in point patterns”. en. In: Statistica Neerlandica 50.3 (1996). eprint: https://onlinelibrary.wiley.com/doi/pdf/10.1111/j.1467-9574.1996.tb01501.x, pp. 344–361. ISSN: 1467-9574. DOI: 10.1111/j.1467-9574.1996.tb01501.x. URL: https://onlinelibrary.wiley.com/doi/abs/10.1111/j.1467-9574.1996.tb01501.x (visited on 02/29/2024).

[S34] Joshua A. Bull and Helen M. Byrne. “Quantification of spatial and phenotypic heterogeneity in an agent-based model of tumour-macrophage interactions”. en. In: PLOS Computational Biology 19.3 (Mar. 2023). Publisher: Public Library of Science, e1010994. ISSN: 1553-7358. DOI: 10.1371/journal.pcbi.1010994. URL: https://journals.plos.org/ploscompbiol/article?id=10.1371/journal.pcbi.1010994 (visited on 11/27/2023).

[S35] Ulrich Bauer. “Ripser: efficient computation of Vietoris–Rips persistence barcodes”. en. In: Journal of Applied and Computational Topology 5.3 (Sept. 2021), pp. 391–423. ISSN: 2367-1734. DOI: 10.1007/s41468-021-00071-5. (visited on 03/06/2024).

[S36] Dashti Ali et al. “A Survey of Vectorization Methods in Topological Data Analysis”. In: IEEE Transactions on Pattern Analysis and Machine Intelligence 45.12 (Dec. 2023). Conference Name: IEEE Transactions on Pattern Analysis and Machine Intelligence, pp. 14069–14080. ISSN: 1939-3539. DOI: 10.1109/TPAMI.2023.3308391. URL: https://ieeexplore.ieee.org/abstract/document/10235748 (visited on 03/06/2024).

[S37] Fabian Pedregosa et al. “Scikit-learn: Machine Learning in Python”. en. In: Journal of Machine Learning Research 12 (2011), pp. 2825–2830.

[S38] Andrzej Kwinta and Joanna Bac-Bronowicz. “Regular polygons in 2D objects shape description”. en. In: Geomatics, Landmanagement and Landscape 4 (2020). ISSN: 2300-1496. DOI: 10.15576/GLL/2020.4.43. URL: https://repo.ur.krakow.pl/info/article/UR62ad65e95a3a45efb63ae1d773088563/ (visited on 11/29/2024).

[S39] Chandler D. Gatenbee et al. “Immunosuppressive niche engineering at the onset of human colorectal cancer”. en. In: Nature Communications 13.1 (Apr. 2022). Number: 1 Publisher: Nature Publishing Group, p. 1798. ISSN: 2041-1723. DOI: 10.1038/s41467-022-29027-8. URL: https://www.nature.com/articles/s41467-022-29027-8 (visited on 02/21/2024).

[S40] Luc Anselin. “Local Indicators of Spatial Association—LISA”. en. In: Geographical Analysis 27.2 (1995). eprint: https://onlinelibrary.wiley.com/doi/pdf/10.1111/j.1538-4632.1995.tb00338.x, pp. 93–115. ISSN: 1538-4632. DOI: 10.1111/j.1538-4632.1995.tb00338.x. URL: https://onlinelibrary.wiley.com/doi/abs/10.1111/j.1538-4632.1995.tb00338.x (visited on 11/05/2024).

[S41] Jiaqi Xi and Jonathan Niles-Weed. “Distributional Convergence of the Sliced Wasserstein Process”. en. In: Advances in Neural Information Processing Systems 35 (Dec. 2022), pp. 13961–13973. URL: https://proceedings.neurips.cc/paperfiles/paper/2022/hash/5a5e9197ea547141b4977a5a198bbaac-Abstract-Conference.html (visited on 03/05/2024).

[S42] Nicolas Bonneel et al. “Sliced and Radon Wasserstein Barycenters of Measures”. en. In: Journal of Mathematical Imaging and Vision 51.1 (Jan. 2015), pp. 22–45. ISSN: 1573-7683. DOI: 10.1007/s10851-014-0506-3. (visited on 03/05/2024).

[S43] Wouter Saelens et al. “A comparison of single-cell trajectory inference methods”. en. In: Nature Biotechnology 37.5(May 2019). Publisher: Nature Publishing Group, pp. 547–554. ISSN: 1546-1696. DOI: 10.1038/s41587-019-0071-9. URL: https://www.nature.com/articles/s41587-019-0071-9 (visited on 09/15/2025).

